# Interpretable Inflammation Landscape of Circulating Immune cells

**DOI:** 10.1101/2023.11.28.568839

**Authors:** Laura Jiménez-Gracia, Davide Maspero, Sergio Aguilar-Fernández, Francesco Craighero, Maria Boulougouri, Max Ruiz, Domenica Marchese, Ginevra Caratù, Jose Liñares-Blanco, Miren Berasategi, Ricardo O Ramirez Flores, Angela Sanzo-Machuca, Ana M. Corraliza, Hoang A. Tran, Rachelly Normand, Jacquelyn Nestor, Yourae Hong, Tessa Kole, Petra van der Velde, Frederique Alleblas, Flaminia Pedretti, Adrià Aterido, Martin Banchero, German Soriano, Eva Román, Maarten van den Berge, Azucena Salas, Jose Manuel Carrascosa, Antonio Fernández Nebro, Eugeni Domènech, Juan Cañete, Jesús Tornero, Javier P. Gisbert, Ernest Choy, Giampiero Girolomoni, Britta Siegmund, Antonio Julià, Violeta Serra, Roberto Elosua, Sabine Tejpar, Silvia Vidal, Martijn C. Nawijn, Ivo Gut, Julio Saez-Rodriguez, Sara Marsal, Alexandra-Chloé Villani, Juan C. Nieto, Holger Heyn

**Affiliations:** Centro Nacional de Análisis Genómico, C/Baldiri Reixac 4, 08028 Barcelona, Spain; Universitat de Barcelona (UB), Barcelona, Spain; Signal Processing Laboratory 2 (LTS2), École Polytechnique Fédérale de Lausanne (EPFL), Lausanne, Switzerland; Heidelberg University, Faculty of Medicine, and Heidelberg University Hospital, Institute for Computational Biomedicine, Heidelberg, Germany; Department of Statistics. University of Granada, 18071, Granada, Spain; Inflammatory Bowel Disease Group, Institut d’Investigacions Biomèdiques August Pi i Sunyer (IDIBAPS), Barcelona, Spain; Centro de Investigación Biomédica en Red de Enfermedades Hepáticas y Digestivas (CIBEREHD), Spain; Center for Immunology and Inflammatory Diseases, Massachusetts General Hospital, Charlestown, 02129, Massachusetts, USA; Broad Institute of MIT and Harvard, Cambridge, 02142, Massachusetts, USA; Harvard Medical School, Boston, Massachusetts, USA; Digestive Oncology, Department of Oncology, Katholieke Universiteit Leuven, Leuven, Belgium; Groningen Research Institute for Asthma and COPD (GRIAC), University Medical Center Groningen, Groningen, Netherlands; Department of Pulmonary Diseases, University of Groningen, University Medical Center Groningen, Groningen, Netherlands; Department of Pathology and Medical Biology, University of Groningen, University Medical Center Groningen, Groningen, Netherlands; Experimental Therapeutics Group, Vall d’Hebron Institute of Oncology, Barcelona, Spain; Rheumatology Research Group, Vall d’Hebron Research Institute, Barcelona, Spain; IMIDomics, Inc; Department of Gastroenterology, Biomedical Research Institut Sant Pau (IIB Sant Pau), Barcelona, Spain; Biomedical Research Network on Hepatic and Digestive Diseases (CIBEREHD), Instituto de Salud Carlos III. Madrid, Spain; Dermatology Department, Hospital Universitari Germans Trias i Pujol. Badalona, Spain; Rheumatology Department, Hospital Regional Universitario Carlos Haya. Málaga, Spain; Gastroenterology Department, Hospital Universitari Germans Trias i Pujol, Badalona, Spain; Rheumatology Department, Fundació Clínic per a la Recerca Biomèdica. Barcelona, Spain; Rheumatology Department, Hospital Universitario Guadalajara. Guadalajara, Spain; Gastroenterology Unit, Hospital Universitario de La Princesa, Instituto de Investigación Sanitaria Princesa (IIS-Princesa). Madrid, Spain; Universidad Autónoma de Madrid (UAM); Section of Rheumatology, Cardiff University, Cardiff, United Kingdom; Section of Dermatology and Venereology, University of Verona, 37129 Verona, Italy; Department of Gastroenterology, Rheumatology and Infectious Diseases, Charité-Universitätsmedizin Berlin, Humboldt-Universität zu Berlin and Berlin Institute of Health, Berlin, Germany; Hospital del Mar Research Institute (IMIM). Barcelona, Catalonia, Spain; CIBERCV, Instituto de Salud Carlos III. Madrid, Spain; Faculty of Medicine, University of Vic-Central University of Catalonia. Vic, Catalonia, Spain; Group of Immunology-Inflammatory Diseases, Biomedical Research Institut Sant Pau (IIB Sant Pau), Barcelona, Spain; European Molecular Biology Laboratory, European Bioinformatics Institute (EMBL-EBI), Hinxton, Cambridgeshire, U.K; On behalf of IMID-Consortium; ICREA, Barcelona, Spain

## Abstract

Inflammation is a biological phenomenon involved in a wide variety of physiological and pathological processes. Although a controlled inflammatory response is beneficial for restoring homeostasis, it can become unfavorable if dysregulated. In recent years, major progress has been made in characterizing acute and chronic inflammation in specific diseases. However, a global, holistic understanding of inflammation is still elusive. This is particularly intriguing, considering the crucial function of inflammation for human health and its potential for modern medicine if fully deciphered. Here, we leverage advances in the field of single-cell genomics to delineate the full spectrum of circulating immune cell activation underlying inflammatory processes during infection, immune-mediated inflammatory diseases and cancer. Our single-cell atlas of >6.5 million peripheral blood mononuclear cells from 1047 patients and 19 diseases allowed us to learn a comprehensive model of inflammation in circulating immune cells. The atlas expanded our current knowledge of the biology of inflammation of immune-mediated diseases, acute and chronic inflammatory diseases, infection and solid tumors, and laid the foundation to develop a precision medicine framework using unsupervised as well as explainable machine learning. Beyond a disease-centered analysis, we charted altered activity of inflammatory molecules in peripheral blood cells, depicting discriminative inflammation-related genes to further understand mechanisms of inflammation. Finally, we have laid the groundwork for developing precision medicine diagnostic tools for patients experiencing pathologic inflammation by learning a classifier for inflammatory diseases, presenting cells in circulation as a powerful resource for patient diagnosis.

## Introduction

Inflammation is a biological response or state of the immune system that serves to protect the human body from environmental challenges, thereby preserving homeostasis and structural integrity of tissues and organs^1^. Inflammatory processes are activated in response to various triggers, such as infection or injury, and involve a multistep defensive mechanism aimed at eliminating the source of perturbation^2–4^. Thus, inflammation represents an altered state within the immune system, which can manifest as either a protective or pathological response^5^. The cellular and molecular mediators of inflammation play pivotal roles in nearly every human disease, encompassing a wide array of biological processes, including the complex interplay of cytokines, myeloid and lymphoid cells^6^.

The initiation of inflammatory processes is driven by cellular stimulation, triggered by the release of proinflammatory cytokines^7,8^. These cytokines exert autocrine and paracrine effects, activating endothelial cells, subsequently increasing vascular permeability. This allows immune cells to infiltrate tissues at the site of infection, facilitated by chemokines. Chemokines are essential for recruiting additional immune cells, playing a crucial role in phagocytosis and pathogen eradication ^9^. In the bloodstream, activated immune cells release cytokines and travel to various tissues. Inflammation is a central driver in cardio-vascular^10^, autoimmune^11,12^, infectious diseases^13,14^ and even cancer^15^. The success of therapies targeting inflammation underscores the importance of understanding the underlying pathways^16–18^. Thus, categorizing patients based on their specific inflammatory cell states in the bloodstream has significant potential for advancing disease management^19^.

Single-cell RNA sequencing (scRNA-seq) is becoming a conventional method for detecting altered cell states in blood, enabling the comparison of transcriptional profiles during perturbations, including inflammation^20^. Previous works revealed cellular profiles across diverse conditions, creating a shared phenotypic space that facilitates comparisons among patients and conditions, and generating a comprehensive view of inflammation^21^. Consequently, differential analysis of cell states and gene expression programs can now guide a holistic understanding of inflammation in acute and chronic diseases to form the basis for future precision medicine tools in diagnostics and novel treatments. In this regard, interpretable machine learning will play a pivotal role to extract disease-driving features from large healthy and disease single-cell references. Eventually, comprehensive models will allow the classification of patients for precise diagnostics and the patient stratification for tailored treatments.

Our study initially defined common immune cell types in peripheral blood, before capturing disease-specific inflammatory cell states that exhibit functional specialization within the inflammatory landscape. Beyond a disease-centered classification, we modeled the expression profiles of inflammatory molecules to define discriminative genes driving immune cell activation, migration, cytotoxic responses, and antigen presentation activities. Ultimately, we proposed a classifier framework based on the peripheral blood mononuclear cell (PBMC) reference, establishing inflammatory immune cell features as a precision medicine diagnostic tool for patients suffering from severe acute or chronic inflammation.

## Results

### An inflammation landscape of circulating immune cells

To chart a comprehensive landscape of immune cells in circulation of healthy individuals and patients suffering from inflammatory diseases, we analyzed the transcriptomic profiles of >6.5 million (6,340,934 million after filtering) PBMCs, representing 1047 patients and 19 diseases, split into a main Inflammation atlas and two validation datasets (**Fig. 1a,b**). Diseases broadly classified into five distinct groups: 1) Immune-mediated inflammatory diseases (IMIDs), 2) acute and 3) chronic inflammation, 4) infection and 5) solid tumors, which were profiled along with healthy donor samples (**Fig. 1a**). We completed our dataset with additional studies to generate a comprehensive resource of immune cell states across inflammatory diseases and beyond (**Fig. 1a**; **Extended Data Fig. 1a; Supplementary Table 1**). Our cohort included various scRNA-seq chemistries (10x Genomics 3’ and 5’ mRNA) and experimental designs (CellPlex and genotype multiplexing), as well as individuals of both sexes and across age groups, to comprehensively capture technical and biological variability (see **Methods**). To learn a generative model of circulating immune cells of inflammatory diseases, we applied probabilistic modeling of the single-cell data using scVI^22^ and scANVI^23^, considering clinical diagnosis, sex and age. scANVI generates a lower-dimensional cell embedding space, before reconstructing gene expression data. Batch effects are removed based on gene-specific parameters, learned during the integration. Its generative probabilistic models proved superior performances in integrating complex datasets compared to other approaches^24^, particularly if cell annotations are available (**Extended Data Fig. 1b-c**). Applied here, the resulting gene expression profiles and the cell embedding space were batch effect corrected, while preserving biological heterogeneity (i.e. previously annotated cell types and states; **Supplementary Table 2**). From the joint embedding space, we initially assigned cells to major immune cell lineages and 15 subpopulations (*Level 1*; **Fig. 1c; Extended Data Fig. 1d**). Then, following a recursive, top-down clustering approach, we obtained a total of 64 immune populations ( *Level 2*), comprehensively resembling immune cell states of the innate and adaptive compartments and allowing a fine-grained description of immune cell types with distinct activation-related transcriptomes ( **Extended Data Fig. 2**; **Supplementary Table 2,3**). High-level compositional analysis (*Level 1*) across diseases revealed significant changes of cell type distributions (scCODA^25^; **Extended Data Fig. 1d**) and validated previously described alterations in peripheral blood cells of patients suffering from inflammatory diseases. For example, we confirmed low levels of Unconventional T Cells (UTCs), Innate Lymphoid Cells (ILCs) and Naive CD4 T cells, together with high proportions of B cells and Monocytes in Systemic Lupus Erythematosus (SLE)^26–28^. Inflammatory Bowel Disease (IBD) patients showed lower levels of UTCs and ILCs^29^ and we observed lower proportions of UTCs accompanied by a larger fraction of Monocytes and B cells in Rheumatoid Arthritis (RA)^30–32^, in line with previous observations. Lymphopenia, a common event during the development of Sepsis^33^, and lymphocytosis, typical of HIV infection^34^, were also confirmed.

**Figure 1.**
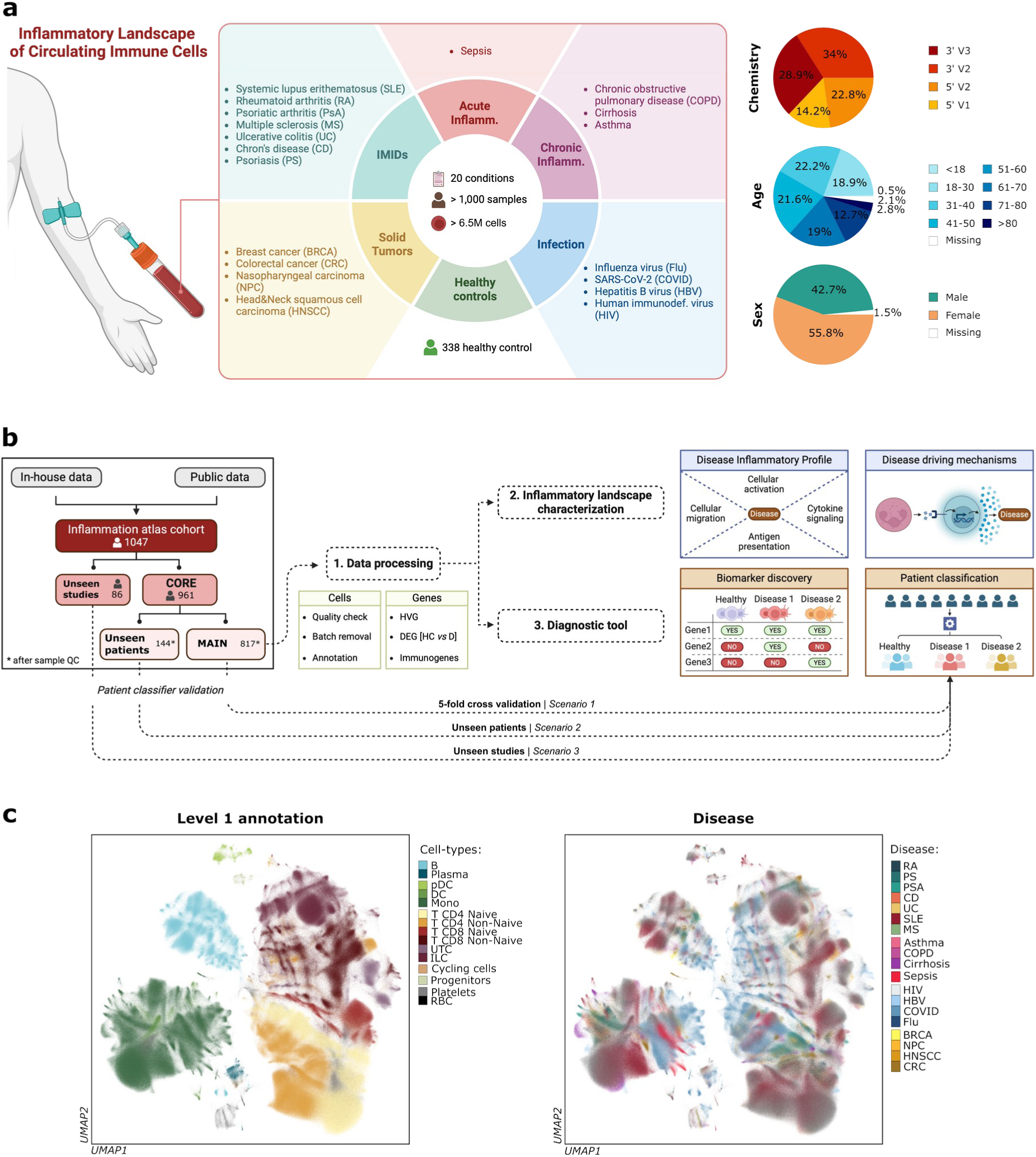
**Inflammation Landscape of Circulating Immune Cells**. (**a**) Left: Schematic overview illustrating the number of cells, samples, and conditions (diseases and disease groups) analyzed. Right: Pie charts displaying metadata related to the scRNA-seq chemistry (10x Genomics assay and version) and patient demographics (age and gender). (**b**) Schematic overview of the analysis workflow followed, detailing the division of the overall dataset into main, unseen patients, and unseen studies. The figure illustrates the specific tasks and analyses performed with each dataset. ( **c**) Uniform manifold approximation and projection (UMAP) embedding for the scANVI-corrected latent space considering the Main dataset (4,435,922 cells) across patients and diseases colored by the major cell lineages (left, *Level 1*) and diseases (right).

Diving deeper into genes and gene programs to characterize inflammatory diseases, our subsequent analysis followed three complementary strategies 1) to identify disease-driving mechanisms (gene signature and Gene Regulatory Network (GRN) activity), 2) to capture discriminative inflammation-related genes (feature extraction) and 3) to classify patients based on their disease-specific signatures (projection). Therefore, we looked at gene expression profiles holistically, but also delineated the inflammatory process by focusing on molecules that trigger immune cell activation, immune cellular migration and extravasation, antigen presentation and cytokine responses (**Supplementary Table 4**)^35–41^. These strategies jointly allowed us to enlarge our understanding of inflammatory processes and their contribution to inflammatory diseases, but also form the basis for precision medicine tools by establishing immune cells in circulation as valuable assets for disease diagnostics.

### Inflammation-related signatures across diseases and cell types

To identify gene expression programs and regulatory mechanisms across cell types and diseases, we first grouped inflammatory molecules into 21 gene signatures that delineate multiple processes, including immune cell adhesion-activation, cellular migration (chemokines), antigen presentation, and cytokine-related signaling (**Supplementary Table 4**). To tailor these signatures to reflect the inflammation landscape of circulating immune cells, we refined these using Spectra, yielding a comprehensive set of 119 cell type-specific factors (**Supplementary Table 4**). We then ran a Univariate Linear Model (ULM) analysis using DecoupleR^42^ on the scANVI-corrected gene expression data, providing an inflammation signature activity score for each group. Finally, we ran a Linear Mixed Effect Model (LMEM) between diseased and healthy samples to highlight disease-specific alterations (**Supplementary Table 5**).

We observed a general trend of increased activity in immune-relevant signatures as compared to healthy donors (>50% increased average signature scores; **Fig. 2a**). For Immune-mediated inflammatory diseases (IMIDs), we found the characteristic upregulation of adhesion molecule signatures, TNF via NFkB signaling, antigen cross-presentation and antigen presenting signatures^43,44^. IFN type 1 and 2 signatures were significantly downregulated in most IMIDs and cell types, except for Non-Naive CD8 T cells that showed an upregulation, pointing to a common cell type-specific mechanism^45^. Remarkably, SLE showed a uniquely strong upregulation of the IFN-induced signature in all immune cell types, accompanied by an upregulation of chemokines and chemokine receptors. Multiple Sclerosis (MS) showed a decreased IFN-induced signature and increased chemokine receptor activity, in line with the migratory capacity of blood cells to infiltrate the brain during the course of the disease^46^. As previously reported, we captured the upregulation of the TNF receptor/ligand signature mainly in Non-Naive CD8 T cells for Sepsis (together with an increase in IFNG response in Monocytes), with a decrease in the other inflammatory signals (adhesion molecules and cytokines) ^47^. In contrast, all chronic inflammatory diseases upregulated the activity of antigen presenting molecules and increased IFN-induced signaling. This IFN-induced signature was also increased in viral infections, such as Flu and COVID-19, while we found a decreased activity in Human Immunodeficiency Virus (HIV) and Hepatitis B virus (HBV). Finally, within solid tumors, Colorectal Cancer (CRC) and Nasopharyngeal Carcinoma (NPC) presented a strong upregulation of TNF via NFkB signaling. Intriguingly, only RA, Psoriatic Arthritis (PSA), Ulcerative Colitis (UC) and Crohn’s Disease (CD) showed an enrichment in the T follicular helper (Tfh) signature in Non-Naive CD4 T cells, highlighting the role of circulating Tfh cells in these diseases. In IMIDs more generally, both Naive and Non-Naive CD4 T cell populations were enriched in T helper signatures, pointing to an early priming of Naive T towards helper T cell-driven inflammation^48,49^. Finally, to assess the similarity of the inflammatory profiles among diseases, we performed hierarchical clustering of the inflammation signature activity score across all cell types ( *Level 1;* **Extended Data Fig. 3a**).

**Figure 2.**
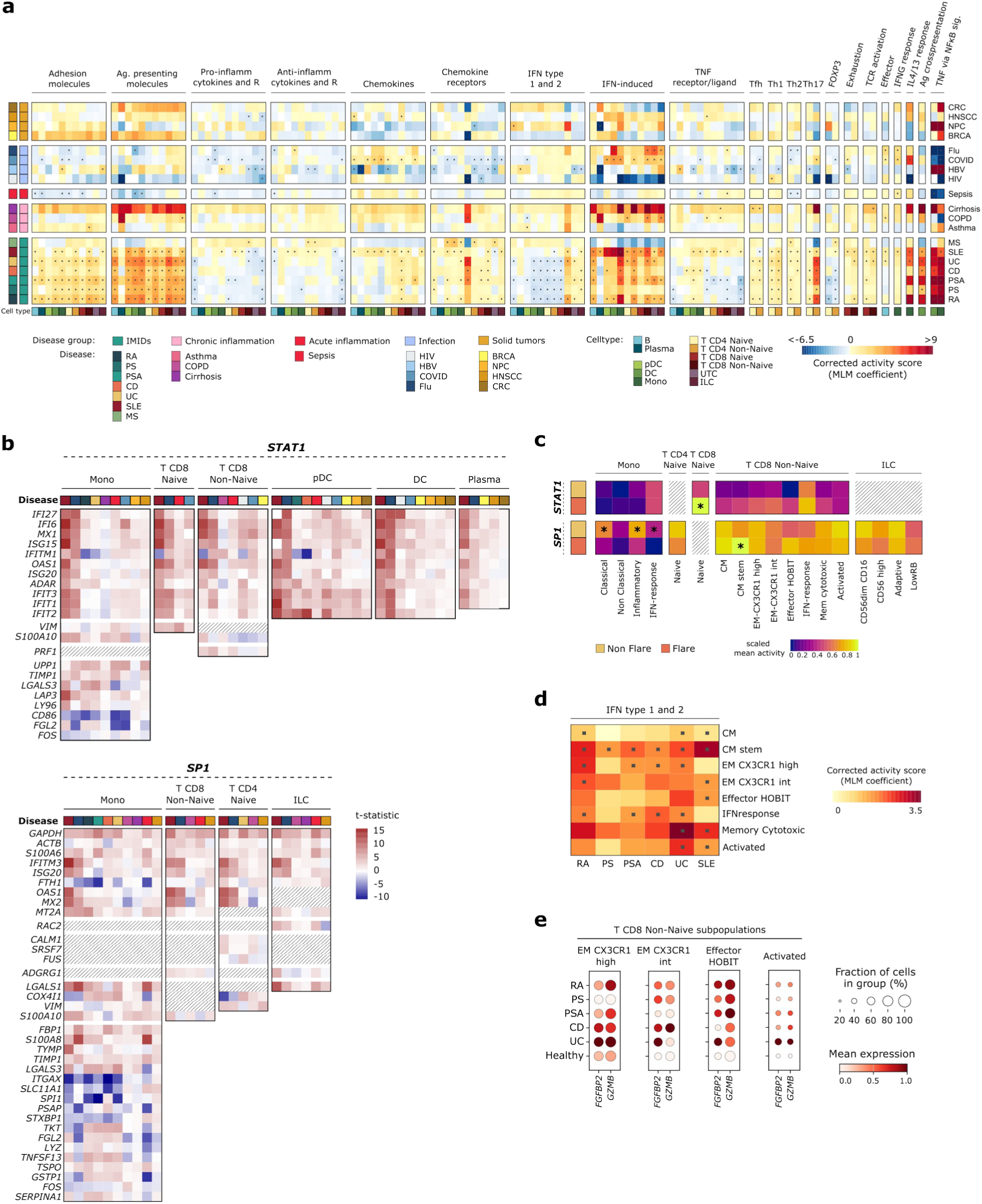
**Inflammation-related signatures across cell types and diseases**. (**a**) Heatmap displaying the corrected signature activity score of the 119 cell type-specific Spectra factors across diseases and cell types (*Level 1*). Here, the corrected signature activity score represents the coefficient value after running a Linear Mixed Effect Model (LMEM) comparing diseases versus Healthy Control (HC) on the Univariate Linear Model (ULM) estimates computed using the cell-type (*Level1*) and patient pseudobulk on the corrected count matrix. The *x-axis* represents the Spectra cell type-specific factor associated with a given function (*top annotation*). The *y-axis* represents the diseases grouped by disease group. (**b**) Heatmap displaying the transcription factor (TF) specificity of STAT1 and SP1 across different cell types and diseases. The t-statistic represents the differential expression of genes between diseased and healthy samples, highlighting shared genes between TF target genes and IFN-induced cell-type signatures. (**c**) Heatmap representing the average scaled TF activity of STAT1 and SP1 across *level 2* population for Flare and non flare patients from *Perez et al*^27^. Asterisk (*) indicates statistical significant changes using the Wilcoxon signed-rank test, FDR adjusted p-value < 0.05. (**d**) Heatmap displaying the corrected IFN Type 1 and 2 signature activity score across Non-Naive CD8 T cells (*Level 2*) across IMID diseases. Here, the corrected signature activity score represents the coefficient value after running a LMEM comparing diseases versus Healthy Control (HC) on the ULM estimates computed using the cell-type (*Level2*) and patient pseudobulk on the corrected count matrix. (**e**) Dotplot showing the uncorrected average expression of the *FGFBP2* and *GZMB* genes from IFN Type 1 and 2 signature (*x-axis*) across different subpopulations of Non-Naive CD8 T cells (*Level 2*) on IMID diseases and health (SCGT00 study). The dot size reflects the percentage of cells of each disease expressing each gene, and the color represents the average expression level. For panels (**a**) and (**d**), significant signature activity differences between disease and HC are marked with a dot ( **·**) (Linear Mixed Effect Model (LMEM), FDR Adjusted p-value < 0.05).

Expanding previous discoveries of a pan-immune IFN-response in SLE patients^27^, especially in the myeloid compartment^50^, we identified a pronounced IFN-induced in SLE across all cell types ( **Figure 2a**). To elucidate the regulatory mechanisms and, specifically, transcription factors (TF) driving the IFN activity, we conducted a GRN analysis of IFN-related factors (see **Methods**). This identified STAT1 and SP1 as the primary regulators of the IFN-induced signature, each TF with distinct cell type specificities, suggesting distinct GRNs to regulate the IFN activity in a cell type-specific manner ( **Supplementary Table 6**). Monocytes and Non-Naive CD8 T cells showed enrichment for both STAT1 and SP1. In contrast, Naive CD8 T cells, plasmacytoid Dendritic Cells (pDC), DC, and Plasma cell populations were uniquely enriched for STAT1 activity, whereas Naive CD4 T and ILC populations showed specificity for SP1 activity (**Fig. 2b**). Further examination of STAT1 and SP1 target genes revealed their pronounced upregulation across various cell types and most pronounced in SLE samples (**Fig. 2b; Extended Data Fig. 3b**; **Supplementary Table 6**). Further dissection of the regulatory dynamics in SLE patients at cell state level (*Level 2*) confirmed an increased activity of STAT1 and decreased activity of SP1 in IFN-response Monocytes and IFN-response CD8 T cells compared to other myeloid and CD8 T cell populations (**Extended Data Fig. 3c**)^51^. Interestingly, SP1 was upregulated in classical and regulatory Monocytes, as well as central-memory (CM) and Activated CD8 T cells, which have been described to play a particular role in the development of SLE (Wilcoxon signed-rank test, FDR Adjusted p-value < 0.05**; Supplementary Table 6**). Finally, STAT1 and SP1 showed distinct activity during disease progression (flare and non-flare patients; **Fig. 2c**; **Supplementary Table 6**). STAT1 activity increased during flares, most notably in CD8 T cell populations, while SP1 activity increased mostly in myeloid populations in the absence of flares. These results provide insights into the distinct regulatory mechanisms driving disease activity in SLE, further highlighting the cell type-specific roles and disease progression dynamics of STAT1 and SP1.

Considering distinct cell types as unique contributors to the inflammatory immune landscape, IFN signatures have been used as a biomarker to define disease activity in autoimmune diseases ^52^. However, it remains elusive which immune subpopulations contribute to these signatures to guide the selection of specific therapeutic interventions. Observing an enriched IFN type 1 and 2 activity in Non-Naive CD8 T cells in IMIDs, we next seeked to discover subpopulations as the signature driver. Here, we observed a significant upregulation across almost all CD8 T cell populations, however, with a differential pattern across diseases. We then decomposed the signal to gene-level, based on the gene weights, to identify the most relevant contributors (**Extended Data Fig. 3d,e)**. Intriguingly, *FGFBP2 and GZMB* showed increased expression levels, with restriction to specific effector-memory (EM) CD8 T cell subtypes (EM CX3CR1 high, EM CX3CR1 int, Eff HOBIT and Activated; **Fig. 2e**). Of note, *FGFBP2* and *GZMB* were recently described as markers of CD8 T cells localized to areas of epithelial damage, where they potently express IFN, driving the IFN response in the DCs within this niche^45^.

### Functional gene selection through interpretable modeling

Gene discovery using linear models (such as the above applied ULM) or standard differential expression analysis suffers from the limitation that genes are considered independently. Thus, we considered the possibility of categorizing cells to their respective disease origin through an interpretable machine learning pipeline, to guide the selection of functional disease discriminatory genes (see **Methods; Supplementary Table 4**). Therefore, we next applied a supervised classification approach, together with a post-hoc interpretability method, to allow the inference of the gene-wise importance, stratified by disease and cell type (*Level1)*. We based our strategy on Gradient Boosted Decision Trees (GBDTs), a state-of-the-art machine learning technique proven to be effective in complex tasks with noisy data and non-linear feature dependencies^53^. GBDTs iteratively build an ensemble of decision trees, by trading the complexity of the model (i.e. the number of trees) with its generalization power. Here, we used the XGBoost library^54^ and tuned its hyperparameters through the Optuna library^55^ (see **Methods**). We executed the analysis considering each cell type (*Level 1*) independently, mitigating the impact of cell-specific expression profiles. We applied the pipeline on the corrected gene expression profiles after scANVI integration, but also tested uncorrected log-normalized data as an input. Overall, we achieved high performance to assign each cell to the correct disease label with a Balanced Accuracy Score (BAS) of 0.87 and a Weighted F1 score (WF1) of 0.90, computed on a test set of 20% cells, when starting from scANVI-corrected data (**Fig. 3a**). Instead, log-normalized counts resulted in decreased BAS and WF1, highlighting the improvement provided after batch correction (0.65 and 0.78; **Fig. 3a**). Performances were consistent among cell types, with less abundant cell populations obtaining generally lower scores (e.g., Plasma cells, BAS = 0.78, WF1 = 0.80; **Extended Data Fig. 4**). Noteworthy, Flu obtained the lowest recall (0.03), mainly being classified as COVID-19 (74% of cells), likely due to the low cell numbers (Flu accounts for ∼0.004% of total cells).

**Figure 3.**
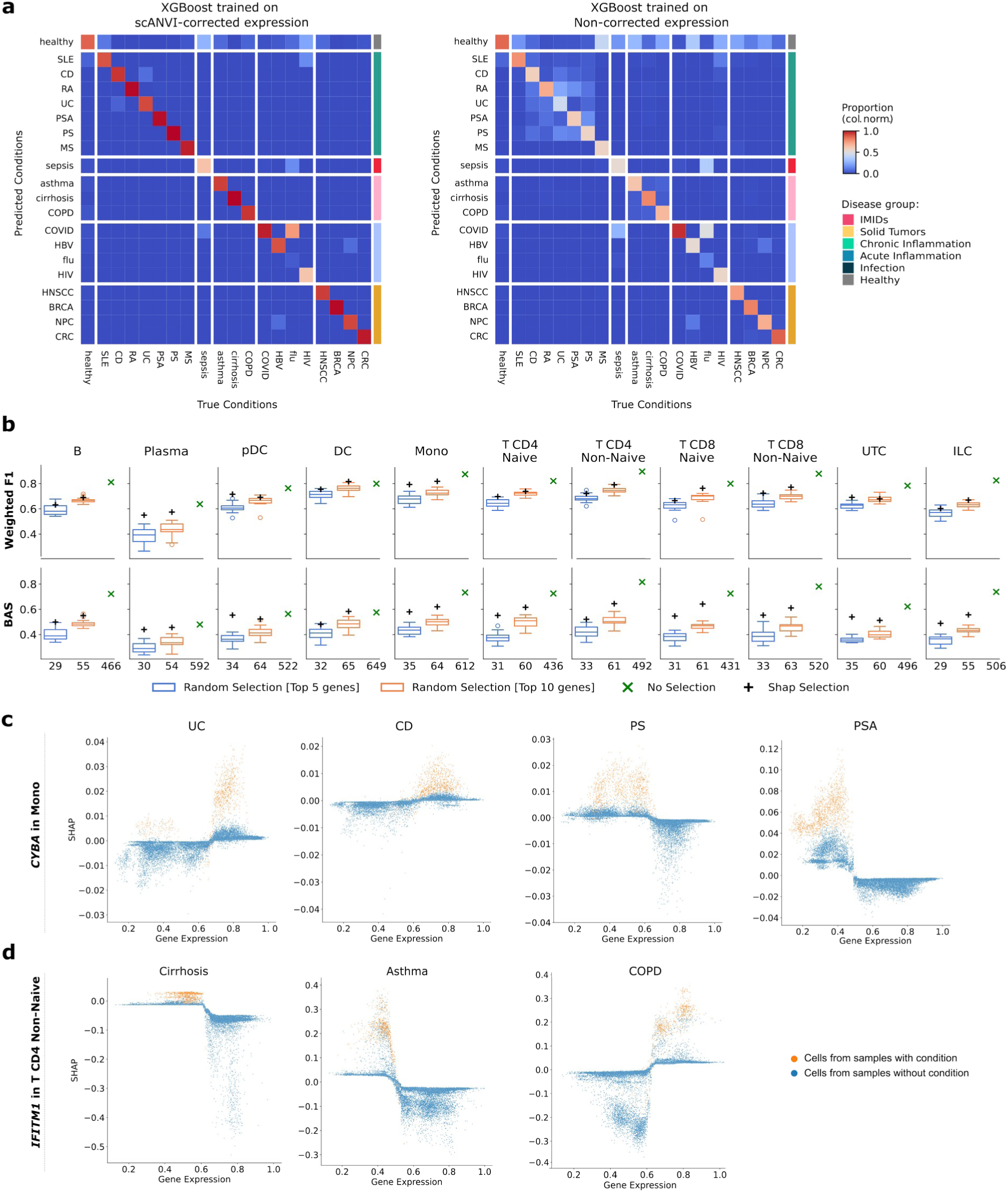
Functional gene discovery using interpretable machine learning. (**a**) Normalized confusion matrices displaying proportion of predictions belonging to each true condition. Diagonal values correspond to the Recall metric. XGBoost was trained on the scANVI batch corrected **(left)** or batch-uncorrected **(right)** log-scaled cell expression profiles. (**b**) Validation of SHAP-based gene selection using XGBoost trained with a nested cross-validation on unseen studies’ cells. Each point corresponds to the average left-out fold performance, for each best configuration of each fold combination. The boxplots plot report the Weighted F1-score (WF1, **top**) and the Balanced Accuracy Score (BAS, **bottom**) computed considering top 5 and 10 genes (among the ones expressed in at least 5% of the total cells), for each inflammatory condition present within the unseen studies dataset (i.e., healthy, sepsis, CD, SLE, HIV, cirrhosis, RA, COVID) according to the SHAP values, across cell-types (*Level 1*). For the same number of genes, we report the performance scores of 20 random selected genes. The performance of the classifier when trained on the whole gene set, consisting of the genes expressed in at least 5% of the total cells is also reported. (**c**) Scatter plot of max-normalized gene expression against SHAP values computed for *CYBA* gene on Monocyte population (annotation *Level1*) and considering the output of XGBoost for a given disease (UC, CD, PS and PSA, from left to right). ( **d**) Scatter plot of max-normalized gene expression against SHAP values computed for *IFITM1* gene on T Non-Naive CD4 population (annotation *Level1*) considering the output of the XGBooost for a given disease (Cirrhosis, Asthma and COPD, from left to right). In Panels (**c**) and (**d**), we limited the visualization to up to 60,000 cells, sampling an equal percentage from each patient corresponding to 5% and 7.5% of Monocytes, T Non-Naive CD4 cells, respectively. Cells belonging to samples with or without the given condition (disease) are marked in orange or blue respectively.

As GBDTs require post-hoc interpretability tools in order to infer explanations, we computed SHAP (SHapley Additive exPlanation) values^56^, shown to provide attributions that are locally consistent and that can be aggregated into global explanations. SHAP values explained the output of the classifier, in our case the predicted conditions, as a sum of contributions of each feature, that is, the gene expression profiles, while also considering its interactions with other genes. By combining the two approaches, we obtained a rich resource of gene rankings based on their ability to discriminate inflammatory conditions across different cell types. When validating the top-ranked genes by predicting the inflammatory disease label of unseen studies, we confirmed that the SHAP gene rankings provide a better feature selection for disease-discrimination than random gene sets (**Fig. 3b**). Moreover, ordering the genes by their importance within immune cell types (*Level1*) and diseases, identified previously described biomarkers, such as *STAT3* in CD4 T cells for RA samples^57^ and IFN genes in multiple cell types for SLE samples^50^ (**Extended Data Fig. 5a**).

The SHAP values of *CYBA*, stood out as a strong candidate marker to classify diseases affecting barrier tissue, PSA, Psoriasis (PS), UC, CD, Chronic Obstructive Pulmonary Disease (COPD) and Asthma (**Fig. 3c; Extended Data Fig. 5b, 6**). *CYBA* encodes the primary component of the microbicidal oxidase system of phagocytes. In line, the importance of the gene across diseases was seen mainly in Monocytes (**Extended Data Fig. 5b, 6**). Interestingly, high expression of *CYBA* drove the model to classify intestinal and pulmonary inflammatory diseases (UC, CD, COPD and Asthma), whereas reduced levels of expression were relevant to classify skin-related diseases (PS and PSA) (**Fig. 3c; Extended Data Fig. 5c**). Mutations in *CYBA* cause an autosomal recessive chronic granulomatous disease with patients showing an impaired phagocyte activation and failing to generate superoxide. Consequently, patients show recurrent bacterial and fungal infections in barrier tissues, including the skin^58^. Thus, we hypothesize that reduction of *CYBA* in skin-related IMIDs leads to an impaired immune barrier function and frequent recurrent infections causing localized, symptomatic flares of PS and PSA. On the other hand, reactive Oxidative Species (ROS) produced by mucosa-resident cells or by newly recruited innate immune cells are essential for antimicrobial mucosal immune responses and defense against pathogenic attack^59^. In UC, CD, COPD and Asthma, an upregulation of *CYBA* may result in the accumulation of superperoxide and ROS through its oxidase function, a hallmark of the diseases ^60^. Importantly, UC mouse models treated with superoxide dismutase showed significantly attenuated UC disease burden in a dose-dependent manner and reduced lipid peroxidation in colonic tissue. Simultaneously, leukocyte rolling and adhesion in colonic venules of colitis rats were significantly reduced, contributing to strongly reduced inflammatory phenotypes^61^.

Further exploring SHAP value ranks, highlighted the importance of *IFITM1* for all chronic diseases (COPD, Asthma and Cirrhosis; **Extended Data Fig. 5d, 6**). *IFITM1* inhibits viral entry into host cells by preventing the fusion of the viral membrane with the host cell membrane^62^. The importance of *IFITM1* was mainly observed in lymphoid cells, specifically CD4 Non-Naive T cells and ILCs ( **Extended Data Fig. 5d, 6**). In CD4 Non-Naive, higher *IFITM1* expression drives the model towards classifying COPD, whereas lower *IFITM1* expression shifts the classification towards cirrhosis and asthma (**Fig. 3d**). ILC populations showed a similar profile, with *IFITM1* emerging as a key disease-discriminative gene (**Extended Data Fig. 5e**). In line, T and ILC cell accumulation is associated with the decline of lung function and severity in COPD patients. T cells also mediate autoimmune responses by inducing the production of IgG autoantibodies in B cells of COPD patients^63^. We hypothesize that chronic inflammation triggers higher expression of *IFITM1* in lymphoid cells, thereby facilitating their accumulation^64^.

### Classifying unseen patients by reference-mapping

The ability to accurately classify cells belonging to different cell types according to their respective diseases prompted us to classify patients based on their disease of origin, creating the basis for a universal classifier as a precision medicine tool for inflammatory diseases. Single-cell information laid a foundation for better understanding the diversity and traits within the populations, but classifying new patients remains a challenge due to data sparsity, noise, and batch-effects across studies. By considering each patient as an ensemble of expression profiles across all circulating immune cells, we learned a generative model while integrating the single-cell reference as a basis to project new patients from a query dataset into the same embedding space. Such strategy allowed us to map unseen and unlabeled query patient data into our reference embedding space, providing a common ground for downstream classification.

Projecting expression data into a lower dimensional space is a common strategy to reduce noise ^65^ and to map query data into a reference atlas^66^. Here, we introduce a novel computational framework to exploit the cell embeddings for classification of patients into conditions (e.g. inflammatory diseases), thus, turning the single-cell reference into a diagnostic tool (**Fig. 4a, Extended Data Fig. 7**). Therefore, we first generated the embeddings with scANVI of both the reference and the unseen query datasets, while also transferring the cell labels to the latter. Then, we defined a cell type pseudobulk profile per patient by averaging the embedded features of the corresponding cells (*Level1*; see **Methods**). Next, we trained an independent classifier to assign correct disease labels, considering one cell type at a time. We handled uncertainty at cell type level via a majority-voting system to determine most frequent conditions. To assess the performance of our framework, we proposed three Scenarios: 1) a 5-fold cross-validation splitting the full reference atlas into five balanced sets, 2) a dataset with unseen patients, 3) an dataset with unseen studies (**Fig. 4b**). Samples from unseen studies have been excluded from our data processing pipeline before quality control, while those corresponding to unseen patients immediately after. We consider these scenarios a representation of the challenges of data integration, where a classifier is expected to account for the unseen batch-effects.

**Figure 4.**
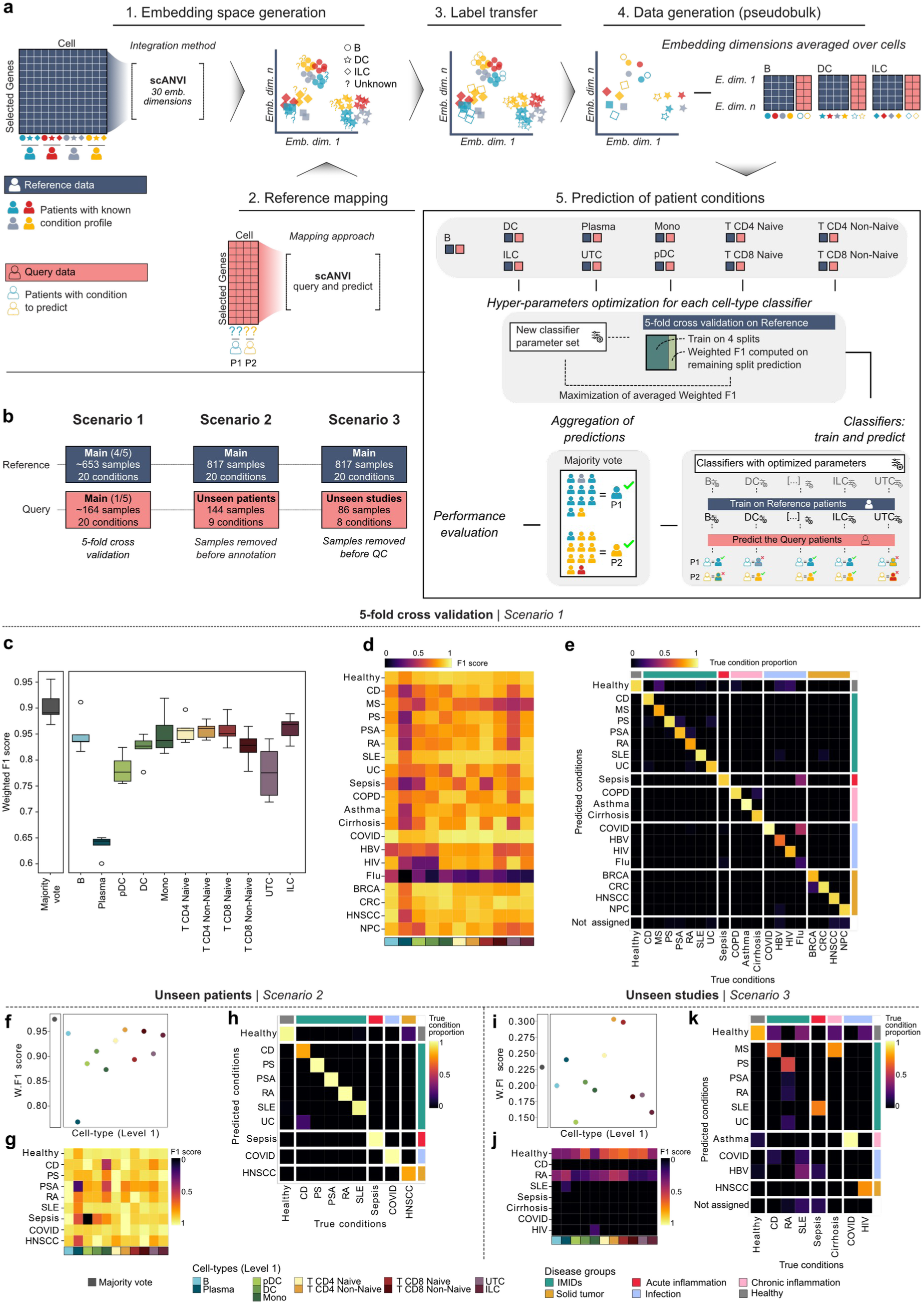
**Schematic representation of the patient classifier pipeline and performance evaluation**. (**a**). Schematic representation of the patient classifier pipeline. (**b**) Description of the three performance evaluation scenarios. In our datasets we always have only one sample for each patient. (**c-e**) Performance evaluation in Scenario 1 (5-fold cross validation), showing (**c**) Distribution of Weighted F1-scores for each left-out split; **(d)** F1-score for each combination of cell-type and disease, after aggregating all the predictions of the left-out folds; (**e**) Normalized confusion matrices displaying proportion of predictions belonging to each true condition after aggregating all the predictions of the left-out folds. Main diagonal values correspond to the Recall metric. ( **f-g**) Performance evaluation in Scenario 2, showing (**f**) Weighted F1-scores for unseen patients’ observation; (**g**) F1-score for each combination of cell-type and disease; **(h)** Normalized confusion matrices displaying proportion of predictions belonging to each true condition. Main diagonal values correspond to the Recall metric. ( **i-k**) Performance evaluation in Scenario 3, showing (**i**) Weighted F1-scores for unseen studies’ observation; (**j**) F1-score for each combination of cell-type and disease; **(k)** Normalized confusion matrices displaying proportion of predictions belonging to each true condition. Main diagonal values correspond to the Recall metric.

Our classification strategy achieved high performance in the cross-validation scenario (Scenario 1), resulting in 0.90±0.03 WF1 (minimum 0.87) and 0.85±0.07 BAS (minimum 0.79), averaged across five independent runs (**Fig. 4c; Extended Data Fig. 8**). Consistently with results obtained from functional gene selection analysis, Flu was the only disease that failed to be classified (Recall: 0.18). Training a classifier for each cell type separately allowed us to assess their relevance in distinguishing inflammatory diseases, particularly for diseases with lower overall performance scores (**Fig. 4d,e**). While certain diseases (COVID, COPD and Asthma) were particularly well classified by lymphoid and myeloid cell types, HIV was best classified by naive lymphoid cells (i.e., Naive CD4 and CD8 T cells and B with F1 of 0.83) in line with the tropism of the virus infecting mainly CD4 T cells^67,68^, while dendritic cell types (i.e. DC, and pDC) did not allow correct disease assignment (0.29). Overall, Plasma and UTC showed the lowest BAS (0.53 and 0.67) and WF1 (0.64 and 0.78), highlighting the strength of our majority voting approach. Increasing the complexity by classifying unseen patient samples (Scenario 2), the performance remained very high, with a BAS of 0.95 and a WF1 of 0.98 (**Fig. 4f-g**; **Extended Data Fig. 9a**). However, the classification of samples from unseen studies (Scenario 3) resulted in a strongly decreased BAS of 0.12 and a WF1 of 0.23 (**Fig. 4i-j**; **Extended Data Fig. 9a**).

The largest performance drop has been observed between Scenario 2 and Scenario 3, the latter classifying patients from unseen studies. We hypothesized that confounding factors, such as variations in assay chemistry or research centers, combined with the inherent complexity of predicting multiple diseases, hindered the classifier’s ability to generalize. To validate our hypothesis and to provide a path towards a generalizable patient classifier, we next considered only data from diseases generated in the same center with a single assay chemistry (SCGT00 data; **Supplementary Table 1**; **Extended Data Fig. 7**). In contrast to Scenario 2, we stratified the samples by sequencing pool and disease, ensuring that reference and query patients belong to distinct cohorts. Moreover, unlike Scenario 3, the primary sources of batch effects are the independent pools, rather than the chemistry protocols and the studies. This new approach included an independent annotation of the reference patients’ cells (see **Methods; Supplementary Table 3**) and new scANVI integration of the reference data, before projecting cells of the query patients. Importantly, in this context, WF1 and BAS increased to 0.56 and 0.53, respectively, pointing to a highly improved generalization performance when classifying query patients as compared to Scenario 3 (**Fig. 5a-c; Extended Data Fig. 10**). Consequently, we hypothesize immune cells in circulation to serve as a source for building a universal classifier for inflammatory diseases, when developing a single-chemistry diagnostic product to control for assay and center batch effects. While the here used subset of the Inflammation Atlas was limited in cell and patient numbers to achieve thresholds for regulatory approval, future efforts for commercialisation are required to develop large single-chemistry training datasets and respective models to further increase the classification accuracy.

**Figure 5.**
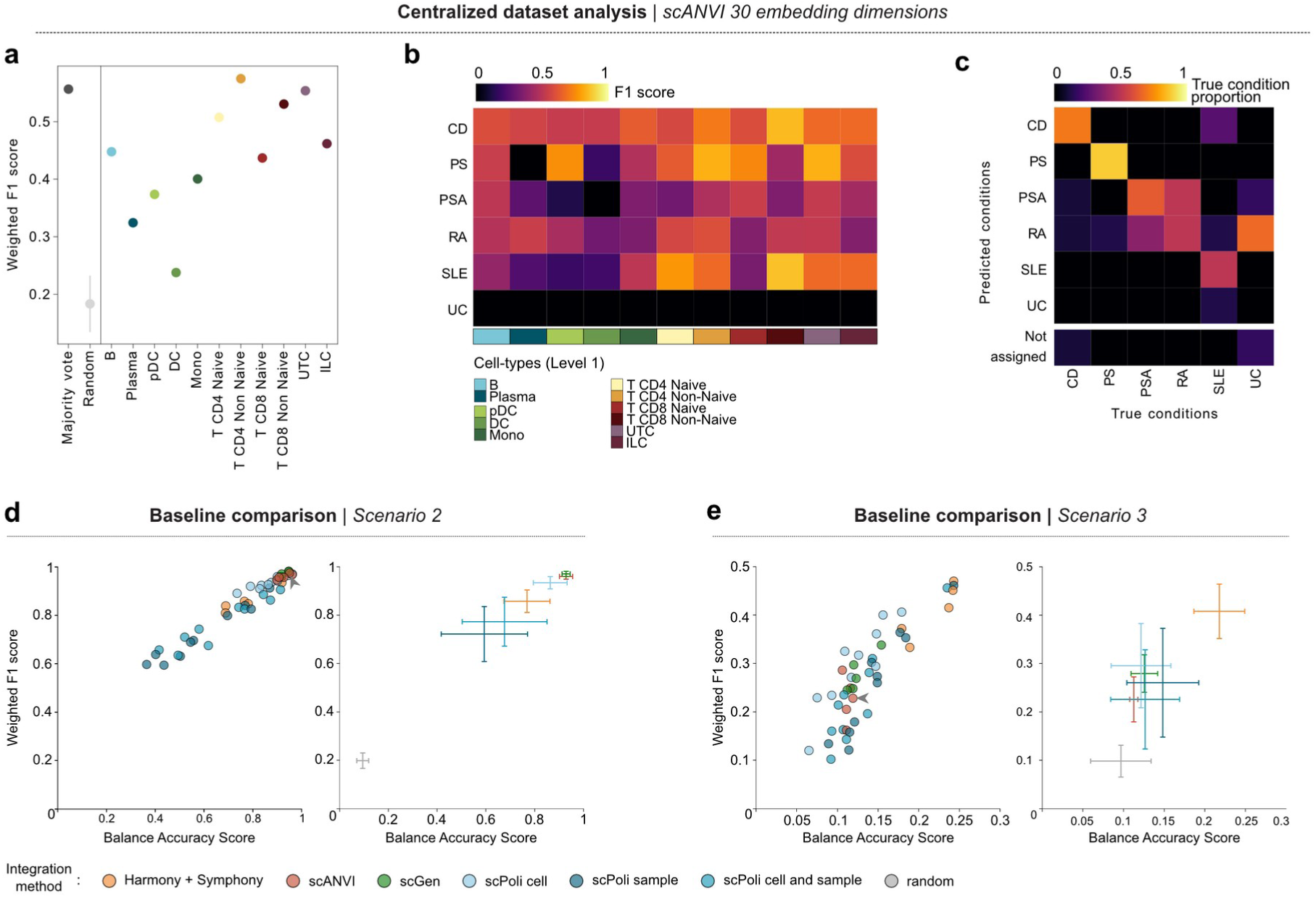
Evaluating patient classifier performance on a centralized dataset and comparison with the state-of-the-art data integration approaches. (**a-c**) Performance evaluation in a Centralized dataset, showing (**a**) Weighted F1-scores for left out pool observation. Mean and standard deviation of weighted F1 score of 100 random condition assignments is reported; ( **b**) F1-score for each combination of cell-type and disease; (**c**) Normalized confusion matrices displaying proportion of predictions belonging to each true condition. Main diagonal values correspond to the Recall metric. ( **d-e**) Performance evaluation in Scenario 2 (**d**) and Scenario 3 (**e**), showing (**left**) the distribution of Weighted F1 and Balanced Accuracy Score for all the configurations of each data integration approach, and (**right**) the mean and standard-deviation of each data integration method, including random assignment. Arrows highlight scANVI configuration applied in Scenario 1.

### Comparing patient classification with state-of-the-art data integration approaches

We selected scANVI as our Inflammation Atlas integration method for its top-ranked performance in data integration benchmarks^24^. To further assess classification performance for the task at hand, we next compared scANVI against other approaches (i.e., Harmony/Symphony^69,70^, scGen^71^, and scPoli^72^) and hyperparameter configurations in Scenario 2 and 3, representing complex classification scenarios with large query datasets. In concordance with our previous results for scANVI, all newly introduced methods achieve high performance in the dataset with unseen patients (Scenario 2); indeed scANVI, scPoli and scGen achieved similar high BAS (>0.94) and WF1 (>0.97), followed by Harmony (BAS: 0.92 and WF1: 0.94), considering the best performing configuration ( **Fig. 5d; Extended Data Fig. 9a,b, 11**). While all the approaches lost predictive power on the unseen studies datasets (Scenario 3), Harmony performed best with a BAS of 0.24 and a WF1 of 0.47 (**Fig. 5e; Extended Data Fig. 9a,c**). Interestingly, scPoli obtained the best result on unseen studies when the learned patient-wise embedding was used to classify patients (BAS: 0.24 and WF1: 0.46), but the same model underperforms on unseen patients (BAS: 0.37 and WF1: 0.60). Since we evaluated different hyperparameters for each approach, we were able to estimate a method’s sensitivity to the chosen configuration. Here, Harmony excels as the best performing and tunable method on samples from unseen studies, with a BAS of 0.22±0.03 and a WF1 of 0.41±0.06, among the considered hyperparameters. In conclusion, all tools showed limited generalization in the most challenging task to classify unseen studies’ patients (Scenario 3), even after hyperparameter adjustments. While linear approaches have less representation power than VAEs, they are also less prone to overfitting and more robust to the hyperparameter choice. Hence, in settings where hyperparameter tuning and validation is not possible due to the lack of condition labels, as in clinical diagnostics, tools like Harmony/Symphony might be preferable to more complex VAEs.

## Discussion

Mapping the plasticity of the immune cells in circulation is now possible by using sequencing technologies that allow an unbiased immuno-phenotyping of single cells^73,74^. Recent technologies enable the sampling of thousands of cells per sample and hundred-thousands per patient cohort, pushing the resolution towards fine-grained cellular maps and increasing the power to identify disease-specific states and programs^75^. To date, single-cell sequencing has been applied to a multitude of inflammatory diseases to determine alterations in cell type composition and to pinpoint disease-driving mechanisms as potential therapeutic targets^20^. However, a complete map of immune cell states across diseases, holistically charting immune plasticity in inflammatory diseases, has been elusive. We reasoned that integrating single-cell transcriptome maps of cells in circulation across a variety of diseases and millions of cells would allow us to extract the full spectrum of features representing inflammatory processes and to generate a comprehensive model of inflammation in circulating immune cells. Our strategy split the analysis into three phases, starting with the supervised extraction of inflammation-related patterns, followed by the discovery of discriminative inflammation-related genes and, finally, a patient classification framework.

Our analysis used complex immune inflammatory signals, such as cytokines, chemokines, activation and antigen presenting signals, that are expressed by immune cells during inflammation, generating gene signatures capturing the intricate inflammatory processes associated with specific diseases^76^. We demonstrated that these inflammatory signatures frequently align with established disease pathways, showing consistency across studies and reinforcing recognized disease mechanisms ^77^. For instance, we confirmed elevated levels of pro-inflammatory cytokines like TNF-α as a hallmark of RA^78^, whereas Type I IFN signature genes were highly active in SLE^79^. We acknowledge though that cell types not captured in this work could further complement the discovery of disease-defining signatures. Here, neutrophils, the most abundant white blood cells in circulation, play important roles in inflammatory diseases^80^. However, the here applied single-cell profiling technologies did not capture neutrophils, presenting a limitation of this work.

The concept of using immune cells as a sensor for diseases is highly intriguing and opens the door for the development of future universal diagnostic tools^81^. For some of the tested diseases, current biomarker strategies may already provide sufficient sensitivity considering the pathology and clinical manifestation of the disease (i.e. lymphopenia as a hallmark of sepsis^47^, or CD4 T lymphocyte count in HIV^82^). Nevertheless, for diseases such as in rheumatology and IBD, many patients are undiagnosed or diagnosed as false positive, and more accurate universal tools are needed^83,84^. Our approach using GBDT, together with SHAP-based interpretability, and a tailored list of functional immune cell molecules, provided explainable outcomes and serves as a rich resource for identifying disease-discriminative genes^53,56^. High absolute SHAP values highlight genes that are important to characterize the inflammatory landscape, and top ranked genes have a high discriminative power between diseases. Additional analysis showed that the top-ranked genes generalize better than a random selection, even when applied on data from unseen studies, confirming the value of SHAP-based ranking. However, given that SHAP values rank genes based on non-linear interactions with other genes, disease– and gene-specific validation in large, independent patient cohorts is required to confirm genes also as biomarkers for clinical diagnostic applications.

We further tested the utility of an Inflammation Atlas as a diagnostic tool by developing a patient classifier based on the latent embeddings after integration. Our approach aimed at leveraging circulating immune cells as a liquid biopsy for diagnostics^72^, by learning to discriminate inflammatory conditions before classifying patients from query datasets. To test such a classification framework, we simulated three scenarios with different degrees of separation of the query patients from the reference cohort. The results obtained for Scenarios 1 and 2 demonstrated that our computational framework can effectively classify patients, indicating that the latent embeddings carry signals associated with inflammatory diseases. To the best of our knowledge, existing patient classifiers have evaluated settings similar to Scenario 1 and 2 (scPoli^72^; MultiMIL^85^). In Scenario 3, we then queried patients belonging to studies excluded from our reference atlas, simulating the application of the Inflammation Atlas as a diagnostic tool. Here, our approach initially failed to generalize to unseen patients, indicating that further optimization was needed to build a generalizable model for more accurate disease diagnostics. To explore the reasons for limited generalization, we performed additional analyses on a centralized dataset, for which all samples were processed at the same center with the same assay chemistry. Here, the improved performance compared to Scenario 3 highlighted the impact of batch effects introduced through differing assay chemistries and centers. We concluded that towards the development of diagnostic tools with regulatory approval, such variability has to be controlled through rigorous standardization efforts during product design and application.

Another important aspect for the classification tasks is the selection of the data integration and reference-to-query mapping tools. While existing atlas-level integration benchmarks guided our selection of scANVI, the tools’ performance for query-to-reference mapping tasks may differ significantly. However, such benchmarking efforts still remained elusive, as they require large datasets to allow cross-validation of both study– and disease-level stratified data. To fill this gap and to facilitate decision-making for building classification pipelines in disease diagnostics, we next provided an evaluation of query-to-reference mapping tools (i.e., scANVI, scGen, scPoli, and Harmony/Symphony) with multiple hyperparameter configurations. Although none of the tools achieved high classification performance in Scenario 3, likely related to uncorrected batch effects between chemistries and studies, our results pointed to Harmony/Symphony as the most robust method. Our evaluation further pointed at the importance of a comprehensive comparison of integration methods for patient classification, since the results are different from existing benchmarks focusing on cell-type annotation, batch correction and biological conservation^24^. While we considered covariates such as gender and age, including additional variables (e.g., treated status and type, comorbidities, disease subtype and duration) could further improve classification performance. Moreover, expanding the feature space to additional data modalities, like cell composition or epigenetic states, could also improve patient classification.

Bringing reference atlases into the clinics remains a complex task, particularly without clear implementation strategies. We contributed to this roadmap by generating a comprehensive landscape of circulating immune cells across inflammatory diseases and healthy controls. Using advanced machine learning pipelines, we demonstrated the usability of interpretable models for discriminating gene importance by cell type and disease, which can be further validated for stand-alone or combinatorial diagnostic tests. Additionally, by classifying patients through generative models that learned the full inflammatory feature space across cell types and diseases, we have laid the groundwork for a universal diagnostic tool for inflammatory diseases. Towards leveraging single-cell technologies in diagnostics, we call for the definition of best practices and quality control standards to reduce batch effects, alongside generating large, controlled training datasets. To allow data integration methods to fully generalize, we need to reduce the confounding factors, as demonstrated by our centralized approach, or largely increase training data size and variability. For the former, diagnostic approval guidelines require the use of predefined, controlled assay chemistries and standard operating procedures, rendering the sequencing pools the main source of batch effects. Alternatively, a reference training data set is generated by multiple centers and diverse chemistries to define a large, heterogeneous atlas, enabling the definition of a foundation model^86^ to pave the way to a universal disease classifier, robust to batch effects.

## Declarations

### Ethics approval and consent to participate

Human blood processed in-house for this project was pre-selected and included within other ongoing studies. All the studies included were conducted in accordance with ethical guidelines and all patients provided written informed consent. Ethical committees and research project approvals for the different studies included in this manuscript are detailed in the following text.

**SCGT00** and **SCGT00val** were approved by Hospital Universitari Vall d’Hebron Research Ethics Committee (PR(AG)144/201). **SCGT01** received the IRB approval by the Parc de Salut Mar Ethics Committee (2016/7075/I). **SCGT02** received the ethics approval by the Medisch-Etische Toetsingscommissie (METc) committee; for asthma patients (ARMS and ORIENT projects – NL53173.042.15 and NL69765.042.19 respectively), for COPD patients (SHERLOck project, NL57656.042.16), and finally, healthy controls (NORM project, NL26187.042.09). **SCGT03** was approved by the Comité Ético de Investigación con Medicamentos del Hospital Universitario Vall d’Hebron (654/C/2019). **SCGT04** and **SCGT06** was approved by the Comitè d’Ètica d’Investigació amb medicaments (CEim) del Hospital de la Santa Creu i Sant Pau (EC/21/373/6616 and EC/23/258/7364). **SCGT05** was approved by the institutional review boards of the Commissie Medische Ethiek UZ KU Leuven/Onderzoek (S66460 and S62294).

### Data and code availability

Single-cell RNA-sequencing (scRNA-seq) in-house generated data and associated count matrices will be available upon publication. The processed scRNA-seq datasets and metadata analyzed in the current study have been deposited at Zenodo: https://zenodo.org/uploads/14851902, and will be available upon publication.

The code to reproduce the full analysis is presented in this article is hosted in the currently private Github repository: https://github.com/Single-Cell-Genomics-Group-CNAG-CRG/Inflammation-PBMCs-Atlas, and will be available upon publication.

### Author information

LJG, JCN and HH conceived the project. JCN and HH supervised the project. LJG, DM, SAF, FC, MBoulougouri and JLB performed the computational and statistical analysis. RORF, HAT, RN and JN provide strategic advice on computational tasks and data interpretation. AA and MBanchero generated VCF files for data demultiplexing. ASM and AMC provided processed objects for validation analysis. MR, DM, GC, MB, and YH generated datasets. TK, PvdV, FA, FP, GS, ER, MvdB, AS, JMC, AFN, ED, JC, JT, JPG, EC, GG, BS, AJ, VS, RE, ST, SV, MCN and SM provided patient samples. LJG, DM, SAF, FC, JSR, PV, ACV, JCN and HH interpreted the results. LJG, DM, SAF, FC, JCN and HH wrote the manuscript with input from all the authors. All authors read and approved the current version of the manuscript.

## Supporting information

JimenezGracia_et_al_2023_SUPPLEMENTARYMATERIAL_v2

JimenezGracia_et_al_2023_SUPPLEMENTARYMATERIAL_Tables_v2

## Acknowledgements

The authors would like to thank the helpful support received by the authors from publicly available data^87,88^ used in the current study by providing processed data in the optimal format. Additionally, we appreciate the great effort put in creating the DISCO database^89^ with multiple-tissue atlases using single-cell datasets. The authors would like to thank the CNAG Scientific IT Unit and the maintainers of the CNAG compute cluster for providing assistance with essential computing resources. Also, the authors would like to thank all the team members, in particular Paula Nieto, Helena L. Crowell, Marc Elosua-Bayes, Mohamed Abdalfatah, and Ramon Massoni-Badosa, for their contributions during brainstorming and discussion sessions.

## Methods

### Atlas of Circulating Immune Cells

The Inflammation Landscape of Circulating Immune Cells atlas has been conceived as a comprehensive resource to expand the current knowledge of physiological and pathological inflammation through the study of circulating immune cells. With this aim, we have included data representing both acute and chronic inflammatory processes, as well as healthy donors. Further details about the included datasets are available (**Supplementary Table 1**).

The project includes in-house single-cell RNA-sequencing data generation from samples shared by our collaborators from several research institutions. Samples were collected with written informed consent obtained from all participants and comply with the ethical guidelines for human samples. Specifically, we generated data from patients suffering Rheumatoid Arthritis (RA), Psoriatic Arthritis (PSA), Crohn’s Disease (CD), Ulcerative Colitis (UC), Psoriasis (PS), Systemic Lupus Erythematosus (SLE) and healthy controls in collaboration with the Vall d’Hebron Research Institute within the DoCTIS consortia [https://doctis.eu/] (**SCGT00** and **SCGT00val**). Additionally, we processed and obtained data from healthy controls in collaboration with the Institut Hospital del Mar d’Investigacions Mèdiques ( **SCGT01**); Asthma, Chronic Obstructive Pulmonary Disease (COPD) and healthy control samples in collaboration with the University Medical Center Groningen (**SCGT02**); Breast Cancer (BRCA) samples in collaboration with the Vall d’Hebron Institute of Oncology (**SCGT03**); cirrhosis samples in collaboration with the Biomedical Research Institut Sant Pau (**SCGT04**); samples of patients suffering Colorectal Cancer (CRC) in collaboration with the Katholieke Universiteit Leuven (**SCGT05**) and, finally, COVID and healthy control samples also in collaboration with Biomedical Research Institut Sant Pau (**SCGT06**).

Moreover, we also included publicly available datasets to complete our cohort. Specifically, we considered data from patients suffering sepsis from **Reyes *et al*.**^47^ and **Jiang *et al*.**^90^, Head and Neck Squamous Cell Carcinoma (HNSCC) from **Cillo *et al*.**^91^, Hepatitis B Virus (HBV) from **Zhang *et al***.^87^, Multiple Sclerosis (MS) from **Schafflick *et al*.**^92^, NasoPharyngeal Cancer (NPC) from **Liu *et al*.**^93^, Human Immunodeficiency Virus (HIV) from **Palshikar *et al*.**^88^ and **Wang *et al*.**^94^, SLE from **Perez *et al*.**^27^, **Savage *et al*.**^95^ and **Mistry *et al*.**^96^, cirrhosis from **Ramachandran *et al*.**^97^, CD from **Martin *et al*.**^98^, COVID-Flu-Sepsis from **COMBAT** (Ahren *et al*.)^99^ as well as COVID from **Ren *et al***.^100^ and healthy controls from **Terekhova *et al*.**^101^ and **10X Genomics** together with the available healthy samples from all the cited studies. The data access identifiers for each project can be found in the **Supplementary Table 1 – Sheet 1**. When raw data was available, we downloaded FASTQ files; otherwise, we retrieved the raw count matrices from the NCBI Gene Expression Omnibus (GEO) [https://www.ncbi.nlm.nih.gov/geo/] or Sequence Read Archieve (SRA) [https://submit.ncbi.nlm.nih.gov/about/sra/], the BioStudies Array Express [https://www.ebi.ac.uk/biostudies/arrayexpress], Broad Institute DUOS [https://duos.broadinstitute.org/], Synapse [https://www.synapse.org], Genome Sequence Analysis (GSA) [https://ngdc.cncb.ac.cn/gsa-human/], CellXGene Data Portal [https://cellxgene.cziscience.com/datasets] and 10X Genomics [https://www.10xgenomics.com/datasets] resources. For all studies, we also collected clinical metadata.

#### Cell barcodes

The *‘cellID’* barcodes assigned were inspired by the TCGA project [https://docs.gdc.cancer.gov/Encyclopedia/pages/TCGA_Barcode/]. Each barcode unequivocally identifies a cell, and it is composed by the *studyID* (project), *libraryID* (10X GEM channel), *patientID*, *chemistry* (only when 3’ and 5’ GEX available for the same sample), *timepoint* (if multiple observations for a patient) and the 10X Genomics cell barcode, respectively [e.g., SCGT00_L046_P006.3P_T0_AAACCCAAGGTGAGAA].

### Sample collection

Human blood samples were collected in EDTA tubes (BD Biosciences). Peripheral blood mononuclear cells (PBMCs) from the **SCGT00, SCGT00val, SCGT02, SCGT04, SCGT05**, and **SCGT06** datasets were isolated using Ficoll density gradient centrifugation (Lymphoprep^TM^, Stem Cell Technologies; Ficoll-Plus, GE Healthcare Biosciences AB). PBMCs belonging to the **SCGT01** and **SCGT03** datasets were isolated using Vacutainer® CPT tube (BD Biosciences). Subsequently, all aliquots were centrifuged following the manufacturer’s protocol. After centrifugation, PBMCs were washed and resuspended in freezing media. Aliquots were gradually frozen using a commercial freezing box (Mr. Frosty, Nalgene, Thermo Fisher Scientific) at –80 °C for 24 h before being transferred to liquid nitrogen for long-term storage.

### Cell thawing and preprocessing

Cryopreserved PBMCs were thawed in a water bath at 37°C and transferred to a 15 ml Falcon tube containing 10 ml of pre-warmed RPMI media supplemented with 10% FBS (Thermo Fisher Scientific). Samples were centrifuged at 350 x g for 8 min at RT, supernatant was removed and pellets resuspended with 1 ml of cold 1X PBS (Thermo Fisher Scientific) supplemented with 0.05% BSA (PN 130-091-376, Miltenyi Biotec). Samples were incubated during 10 min at RT with 0.1 mg/ml of DNAse I (PN LS002007, Worthington-Biochem) in order to eliminate ambient DNA and favor the resuspension of the pellet. Cells were filtered with a 40 µm strainer (PN 43-10040-70, Cell Strainer) to remove eventual clumps and washed by adding 10 ml of cold PBS+0.05% BSA. Samples were centrifuged at 350 x g for 8 min at 4°C and resuspended in an adequate volume of PBS+0.05% BSA in order to reach the desired concentration. Cells concentration and viability were verified with a TC20™ Automated Cell Counter (Bio Rad) upon staining of the cells with Trypan Blue.

### Sample multiplexing by genotyping

PBMCs samples were evenly mixed in pools of 8 donors per library following a multiplexing approach based on donor’s genotype, as done by Kang *et al*.^102^ for a more cost and time-efficient strategy. Importantly, in the case of **SCGT00** libraries were designed to pool samples together from the same disease with different response to treatment (not relevant in this study) whereas in the case of the **SCGT02 Asthma+HC** cohort, 6 samples belonging to patients were pooled with 2 samples derived from non-smoking healthy control individuals. With this approach, we aimed to avoid technical artifacts that could mask subtle biological differences.

### 3’ Cell Plex

PBMCs samples belonging to the **SCGT01**, **SCGT02 COPD+HC**, **SCGT04** and **SCGT06** cohort were multiplexed with 10X Genomics Cell Plex kit following the Cell Multiplexing Oligo Labeling for Single Cell RNA Sequencing Protocol (10x Genomics). While for **SCGT02 COPD+HC** and **SCGT06** projects we pooled 8 samples from patients with healthy controls together, for **SCGT01** and **SCGT04** we only included samples from the condition of interest. Briefly, 0.2-1 million cells were centrifuged at 350x g at RT with a swinging-bucket rotor, resuspended in 100 µl of Cell Multiplexing Oligo (3’ CellPlex Kit Set A PN-1000261, 10x Genomics) and incubated at RT for 5 mins. Cells were washed 3 times with cold 1X PBS (Thermo Fisher Scientific) supplemented with 1% BSA (MACS Miltenyi), all centrifugations being performed at 350x g at 4C. Cells were finally resuspended in an appropriate volume of 1X PBS-1% BSA in order to obtain a final cell concentration of approximately 1600 cells/ul and counted using a TC20™ Automated Cell Counter (Bio-Rad Laboratories, S.A). An equal number of cells of each sample was pooled and filtered with a 40 µm strainer to remove eventual clumps, final cell concentration and viability of the pools were verified before loading onto the Chromium for cell partitioning.

### Cell encapsulation and single cell RNA-sequencing library preparation

Multiplexed samples were loaded for a Target Cell Recovery between 20000 and 60000 cells (corresponding to 5000-7500 cells per sample within each plex). More specifically, samples belonging to **SCGT00**, **SCGT00val** and **SCGT01** cohorts were encapsulated using standard throughput Chromium Next GEM Single Cell 3’ Reagent Kit v3.1, while multiplexes samples belonging to **SCGT02 Asthma+HC and COPD+HC, SCGT04** and **SCGT06** were encapsulated using the high throughput Chromium Next GEM Single Cell 3’ HT Reagent Kit v3.1 in combination with the Chromium X instrument. On the other hand, **SCGT03** and **SCGT05** cohort were loaded in a standard assay with a target recovery of 6-8000 cells per sample using the Chromium Next GEM Single Cell 5’ Reagent Kit v2 (10X Genomics, PN-1000263).

Libraries were prepared following manufacturer’s instructions of protocols CG000315 or CG000390, for the standard assay without and with sample multiplexing, and protocols CG000416 and CG000419 for the high throughput assay without and with sample multiplexing. Protocol CG000331 was instead followed for the **SCGT03** and **SCGT05** cohort. Between 20-200 ng of cDNA were used for preparing libraries and final library size distribution and concentration were determined using a Bioanalyzer High Sensitivity chip (Agilent Technologies). Sequencing was carried out on a NovaSeq6000 system (Illumina) and NextSeq500 using the following sequencing conditions: 28 bp (Read 1) + 10 bp (i7 index) + 10 bp (i5 index) + 90 bp (Read 2), to obtain approximately 40.000 read pairs per cell for the Gene Expression (GEX) library and 2000-4000 read pairs per cell for the CellPlex library.

### Data processing

To profile the cellular transcriptome, we processed the sequencing reads with the 10X Genomics Inc. software package CellRanger (v6.1) [https://support.10xgenomics.com/single-cell-gene-expression/software/overview/welcome] and mapped them against the human GRCh38 reference genome (GENCODE v32/Ensembl 98). This step was applied to the sequencing reads obtained from in-house processed samples and from published projects, when available.

### Genotype processing

Genome-wide genotyping data for patients from the **SCGT00, SCGT00val** and **SCGT02 Asthma+HC** studies were generated from PBMC samples. For **SCGT00,** 184 patients were distributed in four genotyping cohorts (N1=64 samples, N2=32 samples, N3=40 samples and N4=48) and for **SCGT00val,** 32 patients were processed with the Illumina Omni2.5-8 and Illumina GSA MG v3-24 arrays (Illumina, USA), respectively. For **SCGT02 Asthma+HC**, 16 patients were distributed in two genotyping cohorts (N1=8 and N2=8 samples) using Infinium Global Screening Array-24 v3.0 (GSAMD-24v3.0) with the A1 array. Genotyping was done using GRCh37 human genome reference. Data pre-processing and quality control analysis have been separately performed for each genotyping batch of samples at IMIDomics, Inc. (Barcelona, Spain) and Erasmus MC (Rotterdam, The Netherlands). Quality control (QC) analysis was performed using PLINK software (v1.9 and v2). In the **SCGT00** and **SCGT00val** QC analysis, we have identified autosomal SNPs and, using those SNPs from chromosome X, we have confirmed the consistency between SNP-estimated and clinically-reported genders. Then, we have quantified the percentage of SNPs with a minor allele frequency (MAF) higher than 5%. Next, we have computed the percentage of missingness both at the SNP-wise and sample-wise levels. Finally, we have assessed the heterozygosity rate (F) of each sample in order to evaluate if any of the genotyped samples could be contaminated. In the **SCGT02 Asthma+HC** QC analysis, we excluded samples with SNP calling rate lower than 98%.

Patient genotypes (VCF format) were simplified by removing Single Nucleotide Variants (SNVs) that were unannotated (chr 0), located in the sexual Y (chr 24), pseudo-autosomal XY (chr 25), or mitochondrial chromosomes (chr 26). As genotypes were obtained using the human hg19 reference genome, we converted their coordinates to the same reference genome used to mapped the sequencing reads (GRCh38), via the UCSC LiftOver tool [https://genome.ucsc.edu/cgi-bin/hgLiftOver]. LiftOver requires an input file in BED format. Thus, we used a python script [https://github.com/single-cell-genetics/cellsnp-lite/blob/master/scripts/liftOver/liftOver_vcf.py] to convert our VCF file accordingly.

### Library demultiplexing

Multiplexed libraries from **SCGT00, SCGT00val** and **SCGT02 Asthma+HC** cohorts were demultiplexed with cellsnp-lite (v1.2.2) in Mode 1a^103^, which allows us to genotype single-cell GEX libraries by piling-up the expressed alleles based on a list of given SNPs. To do so, we used a list of 7.4 million common SNPs in the human population (MAF > 5%) published by the 1000 Genome Project consortium, and compiled by the authors [https://sourceforge.net/projects/cellsnp/files/SNPlist/]. Then, we performed the donor deconvolution with vireo (v0.5.6)^104^, which assigns the deconvoluted samples to its donor identity using known genotypes, while detecting doublets and unassigned cells. Finally, we discarded detected doublets and unassigned cells before moving on to the downstream processing steps. For CellPlex libraries, we followed a joint deconvolution strategy combining CMO-hashing and genotype-based deconvolution; we generated pools of cells belonging to different samples based on the individual SNPs, and traced back to its donor of origin based on the CMO-hashing. When no genotype is available, the use of this dual approach minimizes the discarded cells.

## Data analysis

All analyses presented in this manuscript were carried out using mainly Python, unless specified otherwise. In particular, we structured our data in Anndata objects^105^ compatible with Scanpy suite^106^, which allowed us to apply single-cell data processing and visualization best practices. The downstream analysis was performed with widely adopted machine learning and single-cell libraries, including scikit-learn^107^ and scvi-tools^108^. All experiments and panels are reproducible with the code released in the project’s GitHub repository.

### Data standardization

Considering the diversity of the datasets included in the reference of circulating immune cells, a standardization step was needed.

#### Gene name harmonization

All datasets were mapped using human GRCh38 genome reference, but the annotation file version might differ, resulting in gene names with multiple aliases or deprecated symbols. To avoid gene redundancy or mismatching, we used ENSEMBL symbols instead of gene names. Then, for datasets without the ENSEMBL symbols, we compared all gene names with the HUGO Gene Nomenclature Committee (HGNC) database (latest version, February 2024) [https://www.genenames.org/], in order to convert them to the latest official HUGO name, merging possible duplicates, and retrieving the corresponding ENSEMBL symbol. For non-official genes, we used the MyGene python interface [https://mygene.info/]^109^ to query the ENSEMBL symbol. Finally, we removed 16 genes categorized as “*artifact*” or “*TEC (To be Experimentally Confirmed)”*.

#### Metadata harmonization

Patient metadata was unified across datasets, using common variable names and values for those present in multiple sources; specifically, we homogenized these variables of interest such as sex, age, disease, diseaseStatus, smokingStatus, ethnicity, or institute. For instance, ‘*M*’, *‘Male’*, and ‘*Hombre’* entries were replaced with ‘*male’*. Additionally, we created a new variable *‘binned_age’* to group patients within a range of 10 years, considering that for the **SCGT01**, **SCGT04**, and **SCGT11** datasets the specific age information was not available. As detailed below, the datasets missing sex and age information were considered as data from **unseen studies** and used to evaluate the patient classifier.

### Data splitting

Some studies in our cohort included patients with samples collected in multiple replicates, different time points, or using different chemistry protocols. In studies with multiple replicates, i.e., **Zhang2022** and **Terekhova2023**, we select samples with the largest number of cells. When multiple timepoints or disease status for the same patient are available, i.e., **Perez2022**, **COMBAT2022** and **Ren2021**, we kept only the samples associated with higher disease severity.

The filtered inflammation atlas cohort has been then split in two datasets, namely: **CORE** and **unseen studies**. Data from **unseen studies** include 86 samples and is used as an independent validation of our patient classifier pipeline. For this dataset, we selected studies that either involve diseases with a large support in our full cohort or lack metadata on sex and age. These chosen studies are: **SCGT00val**, **SCGT06**, **Palshikar2022**, **Ramachandran2019**, **Martin2019**, **Savage2021**, **Jiang2020**, **Mistry2019**, and **10XGenomics**.

After performing data quality control (removing low quality libraries and cells, see section below), the **CORE** dataset includes 961 samples, and was further split into **Main** and data from **Unseen patients**, with 817 and 144 samples each, respectively. We first stratified samples based on the following metadata: *studyID*, *chemistry*, and *disease*. From each of those groups, we randomly selected 20% of samples to be part of the **unseen patients**, provided that they amounted to at least 5 samples. In the patient classifier pipeline, the **Main** dataset is used as a reference, while data from **unseen patients** and **unseen studies** are used as query dataset in two independent scenarios.

The **Centralized Dataset** included samples from **SCGT00** and **SCGT00val**. Since all healthy patients were sequenced in the same pool, we did not take them into account. Then, since multiple samples were multiplexed and sequenced together, we split them stratifying by sequencing *patientPool* to generate both the reference and the query datasets that include at least one pool for all the IMID diseases (i.e., RA, PS, PSA, CD, UC, and SLE). Further information about the samples classified in each group are detailed in the **Supplementary Table 1**.

### Data quality control

We performed data Quality Control (QC) on the **CORE** dataset by computing the main metrics (i.e. library size, library complexity, and percentage of mitochondrial, ribosomal, hemoglobin and platelet-related gene expression) on the count matrix. Metric distributions were visualized grouping cells by library (10X Genomics) and by considering their chemistry (3’ or 5’ prime, and their version). Consequently, we removed low quality observation using permissive thresholds, while the robust cleaning process was performed during cell annotation tasks. In particular, we initially excluded the low quality libraries across datasets (<500 cells or <500 median genes recovered). Next, we removed low quality cells with a very low number of UMIs (<500) and genes (<250), or with a high percentage of mitochondrial expression (>20% for 3’ V3 and 25% for 5’ and 3’ V2), as it is indicative of lysed cells. Then, we removed barcodes with a high library size (>50000 UMIs for 3’ V3 and 5’ V1, > 40000 UMIs for 5’ V2 or >25000 UMIs for 3’ V2 chemistry) or with a high complexity (>6000 genes for 3’ V3, 5’ V1 and 5’ V2 or >4000 genes for 3’ V2 chemistry). After cell QC, we also removed low quality libraries (<250 cells), low quality samples (<500 cells or <500 median genes recovered) as well as cells from a library if this patient recovered a low total number of cells (<50 cells). In addition, we eliminated genes that were detected in less than 20 cells in less than 5 patients, keeping a total of 22838 genes. Lastly, we computed the cell cycle score using the gene list provided by the function *cc.genes.updated.2019()* from the Seurat library^110^ (v4.3.0.1), and defined the different cell cycle ‘phases’ (G1, G2M, and S). Before the dataset clean-up, we predicted doublets with Scrublet^111^ using the function *scanpy.external.pp.scrublet()* from the *scanpy* library (v1.9.8), which provides a score to flag putative doublets, but without filtering them out at this stage. Consequently, during the clustering and annotation step, the clusters co-expressing gene markers from different lineage/population and high doublet score were assessed to determine whether a specific cluster could be classified as a group of doublets and subsequently excluded. After this step, the **CORE** dataset was split into **Main** and data from **Unseen patients**, as explained above.

Quality control on the data from **unseen studies** was performed independently. We applied the same approach described above but, we filtered only bad quality libraries and cells.

### Data processing for annotation

#### Annotation strategy

To identify all the immune cell types and states present in the human blood, we employed a recursive top-down approach inspired by previous work done by La Manno et al.^112^ and Massoni-Badosa et al.^113^. Starting with 4,918,140 cells and 817 patients from the **Main** dataset, we divided the annotation into several stages. Briefly, we first grouped all cells into the primary compartments within our study. Subsequently, each compartment was processed aiming to detect potential doublets, low quality cells and cells resembling platelets or erythrocytes (cells with high expression of hemoglobin genes). Additionally, we also placed back some clusters of cells into their corresponding cell lineages, when wrongly clustered due to similar profiles (e.g. T cells found into the NK cell group, or vice versa). Then, we identified the clusters resembling specific biological cell profiles (cell subtypes) obtaining a final number of 64 different subpopulations, excluding *Doublets* and *LowQuality_cells*, that we defined as annotation *Level2*. Those cell subtypes were grouped into 15 cell populations that we defined as annotation *Level1*. At the end, our inflammation atlas contains 4,435,922 cells. For each group identified in the initial stage (cell lineages), we applied the following tasks, namely: normalization, feature selection, integration, clustering and annotation. In the following, we will always refer to the parameters of the initial stage, while the specifics of the subsequent steps (from lineages to cell types), along with the annotation labels and the marker genes used to define them, can be found **Supplementary Table 3**.

#### Data normalization

Following standard practices, filtered cells were normalized by total counts over all genes and multiplied by a scaling factor of *10*^4^ (*scanpy.pp.normalize_total(target_sum = 10*^4^)). Then, the normalized count matrix *X* was log-transformed as log_e_(X + 1) (*scanpy.pp.log1p())*.

#### Feature selection

Gene selection was performed by identifying the Highly Variable Genes (HVG). Before doing so, we excluded genes related to mitochondrial and ribosomal organules. Also, we skipped T and B cell receptor (TCR/BCR) genes, including joining and variable regions, since they are not useful to describe cell identities, but rather to capture patient specific clonally expanded cell populations within an inflammatory-related condition. Lastly, we excluded Major Histocompatibility Complex (MHC) genes. In order to reduce the influence of a study’s specific composition, and prevent biases in the gene selection task, we preferred genes that are highly variable in as many studies as possible. Therefore, similarly to Sikkema et al.^114^, we first considered each study independently and computed the HVGs using the Seurat implementation^115^ (*scanpy.pp.highly_variable_genes(min_disp=0.3, min_mean=0.01, max_mean=4)*). Then, we ranked genes based on the number of studies in which they are among the HV. Finally, for the initial stage, we determined the minimum number of studies required to compose a HVGs list of over 3000 genes. Applying this strategy, we selected a total of 3236 genes being highly variable in at least 6 studies. In the following steps, we requested more than 2000 HVGs; the minimum number of studies required and the total of selected genes depends on Step and the cells under study. In order to identify Red Blood Cell (RBC) and Platelets, we kept genes associated with erythrocytes, such as hemoglobin subunits, and PPBP (platelet related) in the HVG list. Since such genes are known to be related with ambient RNA when found in other cell types, we subsequently removed them after having annotated the above cell types.

#### Data integration

Our dataset includes single-cell data obtained from multiple studies including different chemistry protocols, inflammatory status samples and a broad range of other clinical features (e.g., age and sex). While this is a strength point of our atlas, such high levels of heterogeneity induced by technical confounding factors and unwanted biological variability resulted in challenging integration tasks before clustering and annotating cell populations. Therefore, we employed scVI^116^, a Variational AutoEncoder (VAE) approach that proves to be one of the most effective integration methods in complex scenarios, particularly when the annotation information is missing^24^. scVI takes as input the raw count matrix to generate an integrated, low-dimensional, embedding space, where the cell states are preserved and the batch effects are reduced. Moreover, scVI’s embedding space can be exploited to cluster and annotate cells based on either known or cluster specific marker genes. Details on the scVI parameters used in each annotation step can be found in **Supplementary Table 7**.

#### Cell clustering

In order to cluster cells into cell types with the Leiden algorithm ^117^, we first built the K-Nearest Neighbors (KNN) graph using scVI’s latent embeddings and *k=20* as the number of neighbors (*scanpy.pp.neighbors(n_neighbors=k)*). We then applied the Leiden algorithm using a resolution of *r=0.1* (*scanpy.tl.leiden(resolution=r)*). The *k* and *r* used in every other step for every lineage can be found in **Supplementary Table 3.**

#### Cell annotation

Cell clusters were manually annotated by immunology experts by comparing the expression levels of canonical gene markers. Moreover, the final step of annotation was performed using the clusters markers obtained performing a Differential Expression Analysis (DEA) among clusters (see **Supplementary Table 3)**. First, we ranked genes to characterize each cluster (*scanpy.tl.rank_genes_groups()*), by considering normalized RNA counts with the Wilcoxon sum rank test. Then, we selected those genes with Log2 Fold Change (log_2_FC) > 0.25, False Discovery Rate (FDR) adjusted p-value < 0.05, and if they were present in at least 25% of cells. Importantly, cells belonging to RBC and Platelet populations were excluded from all the downstream analyses, except for label transfer performed as a step during patient classifier tasks (as explained below).

#### External annotation validation

We compared our independent annotations with the ones available in the largest public datasets (i.e. SLE from **Perez *et al*.**^27^, COVID-Flu-Sepsis from **COMBAT** (Ahren *et al*.)^99^, COVID from **Ren *et al***.^100^ and healthy controls from **Terekhova *et al*.**^101^). To quantify the overlap of cells among groups, we computed the Adjusted Rand index (ARI)^118^ to measure the similarity between our label assignments and the ones performed by the original authors. Further details are available in the **Supplementary Table 2**.

#### Centralized dataset annotation

All previously described steps were applied to process and annotate the centralized dataset (SCGT00), with the following adjustments: 1) standard HVG selection was performed as the dataset included only a single study; 2) the dataset was integrated using “ *patientPool”* as the batch key; and 3) cell annotation was conducted up to *Level1*, recovering the same cell types as in the main inflammation atlas, as this was necessary for the patient classifier. Here, starting with 855,417 cells and 152 patients included in the reference dataset, we recovered 15 cell populations (*Level 1)*, excluding *Doublets* and *LowQuality_cells*. Details on the scVI parameters used in each annotation step can be found in **Supplementary Table 7**, whereas details on the clustering and annotation steps are detailed in **Supplementary Table 3**.

### Feature selection post annotation

#### Gene selection

To improve the quality of downstream analysis to characterize the inflammation landscape it is necessary to perform a gene selection in order to remove dataset specific genes and reduce the batch effect. First, we performed data normalization (as described above) and then removed all the genes that are not expressed (raw count > 0) in at least 1 cell in each study, along with the genes associated to mitochondrial, ribosomal, TCR/BCR, MHC, hemoglobin and platelet cell types. This step retained a total of 14127 genes. Then, we identified three sets of genes: (i) the HVGs, (ii) the Differentially Expressed Genes (DEGs) between healthy and each inflammatory status, and (iii) Cytopus^119^, a manually curated immune-specific gene list.

#### Highly variable genes (HVGs)

Similarly to the feature selection approach described in the annotation section, we selected a total of 3283 HVGs, by using a threshold of at least 3000 genes. In practice, we first ranked the genes based on the number of studies in which they are concurrently highly variable (*scanpy.pp.highly_variable_genes(min_disp=0.3, min_mean=0.01, max_mean=4, batch_key=’libraryID’))*, and then chose a minimum number of studies of 5.

#### Differentially Expressed Genes (DEGs) between healthy and each disease

We obtained a list of DEGs after grouping single-cell gene expression profiles into pseudobulks. Therefore, we first combined the expression profiles of individual cells to produce pseudobulks for every patient and cell-type (annotation level 1), removing groups with no more than 20 cells, using the Python implementation of decoupleR^42^ (v.1.6.0) (*decoupler.get_pseudobulk(min_cells=20, sample_col=’sampleID’, groups_col=’Level1’, layer=’counts’, mode=’sum’)*). Then, we applied the edgeR’s^120^ (v.4.0.16) quasi-likelihood functions to search for DEGs between healthy patients and each other’s inflammatory conditions, by considering one cell-type at a time. Since not all the cell-types were detected in each patient, we didn’t perform the pairwise comparison if one disease had less than 3 pseudobulks. More in detail, for each pairwise comparison we first removed genes with a low expression value (*filterByExpr(y, group = disease)*). Second, we normalized by library size the aggregated raw counts (*calcNormFactors(y, logratioTrim = 0.3)*). Third, we corrected for the main confounding factors, i.e., chemistry protocol, sex and binned age, considering an additive model. One patient was excluded from the analysis due to missing age information. We defined the design of our comparison using the following patsy-style [https://patsy.readthedocs.io/en/latest/formulas.html] formula: ‘*∼0 + C(disease) + C(chemistry) + C(sex) + C(binned_age)’’*. Fourth, we estimated a Negative Binomial (NB) dispersion for each gene using (*estimateDisp()*) which we feed into a gene-wise NB Generalized Linear Model (*glmQLFit(robust=TRUE)*) to test for differentially expressed genes with a quasi-likelihood F-test (*glmQLFTest()*). Lastly, results obtained from each comparison were merged together and the F-test p-values were corrected using the Benjamini-Hochberg FDR procedure implemented in R (*p.adjust(method = ‘BH’)*). Given the corrected p-values and the log_2_FC we selected 6868 DEGs with p-value < 0.01 and absolute log_2_FC > 1.5.

#### Curated immune-specific genes

To be able to track the full spectrum of inflammatory processes, including immune activation and progression, we curated 9 inflammation related functions defined in the literature (1364 genes present in our dataset; **Supplementary Table 4**)^35–41^ and complemented them with a published list of cell type-specific signatures derived from immunological knowledge based on single-cell studies (Cytopus^119^). Specifically, we retrieved all global gene sets for the leukocyte category and the following inflammatory-related cell-type specific factors: Naive and Non-Naive CD4 T cells (*CD4T_TFH_UP*, *CD4T_TH1_UP*, *CD4T_TH2_UP*, *CD4T_TH17_UP*, *Tregs_FOXP3_stabilization*); Naive and NonNaive CD8 T cells (*CD8T_exhaustion*, *CD8T_tcr_activation*); B cells (*B_effector*); Monocytes (IFNG *response*, *IL4-IL13 response*); and DC (DC antigen-crosspresentation).

#### Aggregation of gene sets

We generate the relevant gene set by doing the union of HVGs, DEGs, and the manually curated list. The final number of unique genes is 8253.

### Datasets integration and gene expression correction via scANVI

Atlas-level analysis requires a careful preprocessing of the gene expression profiles to deal with the heterogeneity of the studies, the batch effect and the missing or noisy observations ^121^. scANVI^23^ is one of the existing methods capable of addressing these challenges and has been proven effective on atlas-level benchmarks compared to other integration methods. We validated its performance on our data by using the metrics from the *scib-metrics* package^24^ (v.0.5) (see **Extended Data Fig. 1**).

#### scANVI integration

scANVI is an extension of the scVI model, employed previously for data integration, that also leverages the information of the cell type annotation. We first trained an scVI VAE (*scvi.model.SCVI*), and then trained scANVI (*scvi.model.SCANVI*) starting from the pre-trained scVI model (see parameters in **Supplementary Table 7**). Both models corrected for the *chemistry* batch, while also considering *libraryID*, *studyID*, *sex*, and *binned age* as covariates. After training, we generated the normalized corrected counts by sampling from scANVI’s negative binomial posterior (*SCANVI.get_normalized_expression*). The batch effect was mitigated by sampling and averaging each cell’s expression as if it originated from each chemistry protocol by setting the *transform_batch* parameter to the list of chemistry protocols present in our atlas. To parallelize the gene expression sampling, we implemented a Nextflow (v.24.04) pipeline^122^ that processed each sample independently and aggregated the obtained anndata objects afterwards.

### Comparison of cell-type composition

#### Compositional cell-type analysis

To estimate the changes in the proportions of cell populations across conditions, we applied the scCODA Python package^123^ (v0.1.9), a Bayesian modeling tool that takes into account the compositional nature of the data to reduce the risk of false discoveries. scCODA allows us to infer changes between conditions while considering other covariates, corresponding to the disease status in our setting. scCODA searches for changes between a reference cell type, assumed to be constant among different conditions, and the other cell types. We selected as the reference population the one that showed the lower variance across conditions, excluding rare cell populations (i.e., *Progenitors, pDCs* and *Cycling cells*). This resulted in the selection of DC as the reference cell type for all diseases. scCODA takes as input the count of cells belonging to each cell type in each patient and returns the list of cell types proportion changes with the corresponding corrected p-values (through the False Discovery Rate procedure; FDR). A *patsy-style* formula was used to build the covariate matrix, specified with ‘healthy’ as baseline and sex and binned age as covariates ( *C(disease, Treatment(’healthy’)) + C(sex) + C(binned_age)*), since we are interested in detecting changes between a normal and diseased status. We reported only changes with a corrected p-value < 0.05 and a log _2_FC > 0.2.

### Comparison of gene expression profiles

#### Gene factor inference

In order to expand the list of curated immune-related genes following a data-driven approach, we employed Spectra^25^ (v.0.2.0), a matrix factorization algorithm that enables us to identify a minimal set of genes related to specific functions in the data, i.e., factors. Spectra takes as input cell-type labels to infer global and cell-type specific factors that decompose the overall gene expression matrix and each cell-type submatrices, respectively. Given our list of curated gene sets, we considered the Cytopus ones as global factors, while we regarded all the remaining as cell-types specific factors. The Spectra model was fitted with default parameters with the exception of *lambda*, which was set equal to 0.001. Considering the prohibitive computational resources required for applying Spectra on our single-cell data, we fed the algorithm with the metacell aggregated expression matrix, as described in the paragraph below. Spectra returned a list of 135 factors that are a linear combination of the gene expression from the original matrix. The coefficients included in the matrix can be then used as a proxy of the gene relevance in a given factor.

#### Metacell generation

We generated metacells using SEACells^124^ (v.0.3.3), which aggregates cells by exploiting their distances in a low-dimensional embedding space. Starting from the normalizing data, we selected the top 3000 highly variable genes using the *highly_variable_genes* function in scanpy, with the *seurat* flavor. To define SEAcell’s input embedding space, we calculated the first 50 Principal Components (PCs) and selected those PCs that explain 90% of the total variance observed across all 50 PCs. To avoid biases due to batch effect and other confounding factors, we executed SEAcell for each sample independently. In particular, we generated a number of metacells equal to the number of cells of each patient divided by 50. SEAcell was executed in parallel with Nextflow ^122^. We further filtered the obtained metacells by computing the proportion of the most abundant annotation label ( *Level1*) in each SEAcell group, and then removed the ones with a purity lower than 0.75. Overall, we defined 71108 metacells. Given the assignment of cells to each metacell, we generated each metacell’s gene expression profile by averaging the corresponding cells’ scANVI normalized and corrected expression profiles. Since scANVI returns counts sampled from a negative-binomial distribution, we also log-scaled the obtained metacell profiles.

#### Inflammation-related signature definition

Spectra provided a total of 135 factors, that include a refined gene list for each gene set we used as input. Thus, we need to assign those factors to our original gene sets for retrieving the corresponding biological function. For doing so, we performed enrichment analysis with Univariate Linear Models (UML) available in the Python implementation of decoupleR^42^, to estimate the factors associated with each gene set. We considered the gene coefficients returned by Spectra as response variable and as explanatory variables a vector of weights equal to 1 or 0 if the gene were included in the gene set or not respectively. UML returns an *estimate* and a *p-value* for each enrichment. We correct those p-values for multiple comparison by computing the False Discovery Control with Benjamini-Hochberg procedure, implemented in *scipy* (v.1.12.0) Python library^125^. We kept 125 factors with a positive estimate and an adjusted p-value lower than 0.05. Finally, we assign to each factor the biological function that corresponds to the gene set that provided the highest estimated score.

#### Inflammation-related signature scores

To compare immune-relevant activation profiles across diseases and cell types, we applied an enrichment signature scoring procedure, considering the factors obtained with Spectra^25^. In particular, we applied an Univariate Linear Model (ULM) from decoupleR^42^. First, we generated pseudobulks stratified by cell type (annotations *Level1* or *Level2*) and patients, discarding groups with less than 10 cells. We averaged the scANVI corrected gene expression matrix of each cell belonging to a given pseudobulk, and then log-transformed and scaled the expression values to zero mean and unit variance, to reduce the impact of highly expressed genes. We fitted decoupleR’s ULM by considering pseudobulk expression profiles as the response variable and the gene coefficient returned by Spectra as the explanatory one. We assessed the scores for the 119 cell-type-specific factors only in their corresponding cell-type. The output of the model is a t-student statistic for each combination of pseudobulk and factor, which is used as a proxy for the corresponding biological function activity: positive values are associated with more active functions in a given sample, and vice-versa. To identify the up-or down-regulated biological functions across inflammatory conditions, we compared the activation score between *healthy* and each disease, considering only comparisons that include at least 3 observations in both conditions. To take into account the batch effect induced by studies and chemistry protocols, that still affects the data (see **Extended Data Fig. 1b,c** *scbi* metrics and PCA), we applied a Linear Mixed Effect Model (LMEM). In particular, we fitted the function *mixedlm()* from *statsmodels* Python Library (v.0.14.1) with the formula*: “’Q(“{factor}”) ∼ C(disease, Treatment(reference=“healthy”)) + ‘f’C(chemistry)’*, grouping by *“studyID”*. We corrected the p-value obtained for multiple testing using FDR considering all the comparisons when tested at *Level1*, and within each *Level1* population when tested at *Level2*.

#### Gene Regulatory Network (GRN) Analysis

Pseudobulk matrices were calculated by averaging the corrected and standardized count matrices by cell type and sample. We compute differential expression analysis for each cell type in each disease using healthy individuals as reference. Linear Mixed Effect Model (LMEM) were used to model the expression levels of each gene independently, considering the disease as a fixed effect while modeling the ID of the study as a random effect. We used the mixedlm() function of statsmodels (v.0.14.0) Python package to run the analysis. To associate each cell-type specific “*IFN-induced”* factor with a given Transcription Factor (TFs) regulator, we integrated these signatures with the CollecTRI Gene Regulatory Network^126^ by matching target genes to identify common genes between TF regulons and Spectra signatures. Therefore, each “*IFN-induced”* signature was thus linked to a subset of TF regulons. The activity of each TF was calculated using only the common genes between each TF and each signature, employing the UML from decoupleR^42^ and the z-values obtained from the differential expression analysis. To ensure robustness, only regulons with at least 10 gene targets were considered. This pipeline was applied across “*IFN-induced”* factors and diseases, focusing on the activity in the cell type where the Spectra signature was identified. Negative activities (t-stat < 0) and non-significant results (p-value > 0.05) were filtered out. This analysis identified STAT1 and SP1 as the sole transcription factor regulators of the defined cell types.

To explore disease distribution according to “*IFN_induced*” Spectra signatures, the MOFA^127^ (version 0.3.2) framework was utilized. We used diseases as samples, cell types as views (representing the Spectra signatures), unique gene targets from SP1 and STAT1 as features and z-values from DEG of these targets as values. The convergence mode was set to fast, and the dropR2 argument was set to 0.0001. For the comparison of Flare and Non Flare patients from Systemic Lupus Erythematosus (SLE), non-corrected log1p normalized single-cell expression matrix from *Perez et al*.^27^ was used to further investigate SP1 and STAT1 regulon activities across both categories. Pseudobulk profiles were calculated by averaging by cell type, considering only cell types (annotation *Level 2)* with a minimum of 10 cells and groups that include at least 3 patients in both conditions (Flare vs Non Flare). Prior to calculating TFs activities across samples, we standardized the gene expression data on SLE patients based on healthy individuals. Specifically, for each gene, the mean and standard deviation were calculated from the healthy group, and these statistics were then used to scale the gene expression values across SLE patients. Only gene targets identified in the previous step were used to calculate enrichment using the UML method. Finally, the activity of STAT1 and SP1 was calculated at *Level2* using CollecTRI^126^.

### Immune gene importance evaluation

In this section, we will introduce our pipeline used to obtain a gene importance metric by interpreting cell-type-specific classifiers for disease prediction. All the steps described below were carried out separately for each cell type (excluding RBC, Platelets, Progenitors and Cycling cells). Specifically, the classification task was performed with Gradient Boosted Decision Trees (GBDTs) implemented in the XGBoost library^54^ (py-xgboost-gpu: v2.0.3). Furthermore, interpretability was performed using SHapley Additive exPlanation (SHAP) values^56^ (v.0.45.1), a powerful approach assigning an importance to each gene by also taking into account their interactions.

#### Feature Selection

To focus our analysis on cell type specific inflammatory related signatures we considered only genes relevant in annotated Spectra factors and we further reduce the list by removing cell identity genes (ex. *CD3E*, *MS4A1*) as well as non protein coding genes. This filtering gave a final number 935 genes.

#### Data Processing

We split our data in three parts: the training set, and the validation and testing set, used for hyperparameter tuning and performance evaluation, respectively. We balanced the splits by disease, ensuring that each sample’s cells were included in the same set. Initially, we partitioned the data into five splits using the function *sklearn.model_selection.StratifiedGroupKFold*. Three of these splits were assigned to the training set, while one was designated for validation and one for testing. Accounting for both stratification by disease and patient partitioning might lead to an uneven distribution of cells among diseases. To address this, we assigned splits with a well-balanced distribution of cells to the training and testing sets first.

#### XGBoost Fitting

XGBoost (*xgboost.XGBClassifier)* hyperparameters were tuned using the Optuna library^55^ (v.3.6.0), which employs a Tree-structured Parzen Estimator (TPE) sampling algorithm^128^ to navigate the hyperparameters space. The performance of each model configuration was estimated using the weighted F1 score on the validation set. To reduce the computational cost, we both pruned unpromising hyperparameters and early-stopped the training when no improvement is achieved over 20 steps before the upper bound of 1500. We considered 50 configurations of XGBoost, taking into account the hyperparameters detailed in the **Supplementary Table 7.** Using the best configuration and its corresponding number of training steps (equivalent to the number of estimators), we retrained the best model on the union of the training and validation sets. This time, we did not apply early stopping and increased the number of training steps by 20%, to account for the larger number of training samples.

#### SHAP interpretability

To interpret the decision of the selected XGBoost classifier, we employed the widely-adopted Shapley values^129^ through the SHAP library. SHAP values were computed with *shap.TreeExplainer* using the *observational* “*three_path_dependant*” approach, which is expected to be more representative of the data itself and its causal dependencies than the *interventional* one^130^. Given the potential resource-intensive nature of handling all SHAP values for every cell and disease, especially in terms of storage, we computed their mean and variance across all samples in batches using the Weldford online algorithm^131^. Given a specific cell type *ct*, we have a SHAP value for every gene in every cell, and for each disease: a matrix of real values *sha p^ct^* (*c, g, d*), where *c, g*, and *d* identify the cell, gene, and disease, respectively. The average contribution of a gene *g* for a disease *d* can be computed as *Dsha p^ct^* (*g, d*)=*mea n* ∨ *sha p^ct^* ( *c, g, d*)∨, where *C* is the set of cells.

### Gene Selection Validation

To validate our ensemble of important genes through SHAP values, we tested if our selection generalizes to unseen studies. First, we defined a gene set ***GS*** that included genes expressed in at least 5% of the cells. Then, for each of the 8 conditions included in **Unseen Studies**, we selected from ***GS*** the top *k* ranked genes by SHAP importance. We then trained XGBoost in a nested cross-validation setting on data from **Unseen Studies**, where we performed both hyperparameter tuning and performance evaluation, using only the selected genes as input features. Next, we computed Weighted F1 (WF1) and Balanced Accuracy Score (BAS) to test the performance considering *k=*5 and *k=10*. Given our selection ***S*** of genes, with size |***S***|, we also tested 20 sets of |***S***| randomly selected genes from ***GS***, excluding the ones in ***S*** (i.e., not top-ranked according to SHAP). Lastly, we compared the performances of the models trained on those subsets of genes against XGBboost trained on the whole set of gene ***GS***. The analysis was repeated for each cell-type independently.

### Patient classifier pipeline

In this section, we describe the pipeline used to validate the inflammation atlas as a diagnostic tool. In the following analysis the terms ‘patient’ and ‘sample’ are equivalent, since after Data splitting we kept only one sample foreach patient. The pipeline consists in i) integrating an annotated reference dataset with data integration tools that provide batch-corrected embeddings, ii) mapping a query dataset into the reference to obtain its corrected embeddings, iii) transfer the cell annotation labels from the reference, iv) defining a patient embedding space, and v) training a classifier to predict the patient conditions from the embeddings.

Starting from a large annotated reference dataset, we applied four state-of-the-art integration methods, described below, to obtain a batch corrected embedding. We considered different chemistry protocols as the main source of batch effect, thus we corrected for the *chemistry* covariate. Then, an independent query dataset was mapped into the corrected embeddings. This step provides both batch correction and allows us to transfer cell annotation labels from the reference to the query dataset. To define patient-wise embeddings, we averaged each patient’s cell embeddings by cell type, resulting in an embedding for each cell-type and each patient. To predict the inflammatory conditions of the patients in the independent query dataset, we fit one classifier for each cell type on the reference patient embeddings. Then, we predicted the inflammatory condition of the query patients by returning the most frequent condition among the predictions of every cell-type-specific classifier.

We validated our pipeline considering three different settings. In the first one, we performed a cross-validation on the **Main** dataset, where each left-out split is considered as a query dataset and the remaining as the reference. Moreover, we tested our diagnostic tool on data from **unseen patients** and **unseen studies**, this time using the whole **Main** as a reference.

### Integration Methods

In this section we will explain each data integration method, while the tested configurations of hyperparameters can be found in **Extended Data Fig. 7** and **Supplementary Table 7**.

#### scGEN

scGen is defined by two main components: a Variational AutoEncoder (VAE) and a latent space arithmetic method. The VAE estimates a posterior distribution of latent variables through the encoder, from which we can reconstruct the expression matrices via the decoder (*scGen_model.batch_removal()*). Similarly to commonly employed VAEs, scGen approximates the posterior through a variational distribution, modeled by the encoder and defined as a multivariate Gaussian. When the scGen’s VAE has been fitted on the reference dataset, latent space arithmetic is employed to correct for the batch effect induced by the chemistry protocol used. Within each cell type, scGen first selects the mean *μ_max_*of the most populated batch, and then corrects each batch with mean *μ*_0_ by adding *δ* =*μ_max_ − μ*_0_ to each cell’s embedding. Importantly, the cell type has to be inferred when not known. The final corrected count matrix will correspond to the generated count matrix from the arithmetic-corrected embeddings. Following scGen’s tutorials, we will refer as corrected embeddings to the ones obtained given the corrected expression matrix as input. In order to apply scGen batch correction on the query dataset, we need to also infer the cell types of those cells. This step was performed through label transfer by nearest neighbors, following a similar approach employed in Human Lung Cell Atlas^114^ and introduced in^66^. The idea is to employ (approximate) nearest neighbors through the PyNNDescent^132^ (v0.5.11) (*pynndescent.NNDescent().prepare()*), and infer the most probable cell type in the 10 nearest neighbors (*pynndescent.NNDescent().query())* from the already annotated cells in the reference dataset. To account for the shape of the distribution of the neighbors, a Gaussian kernel was applied instead of using the Euclidean distance. The most probable nearest neighbor cell type is then assigned to annotate new cells.

#### scANVI

We first trained scVI and scANVI on the reference dataset, like the Dataset Integration described before, then we fine-tuned it to the query dataset. Regarding the label transfer, we employed the scANVI *predict()* function with default parameters.

#### Harmony and Symphony

Harmony^70^ and Symphony^69^ are two related methods that integrate a reference and map a query dataset to it, respectively. Harmony takes a PCA embedding of cells as input, along with their batch covariates (*chemistry*). Next, the model represents cell states as soft clusters, where each cell identity is defined as a probabilistic assignment across clusters, with the aim of maximizing diversity among batches within those clusters. Cells are iteratively assigned soft-cluster memberships; those assignments are used as weights in a linear mixture model to remove confounding factors. The result is a new batch-corrected embedded space. The Symphony algorithm starts from the linear model parameters inferred by Harmony to map query cells onto the corresponding embedding space. First, it projects the query gene expression profiles into the same uncorrected low-dimensional space as the reference cells. Next, Symphony computes soft-cluster assignments for the query cells based on their proximity to the reference cluster centroids. Finally, Symphony employs the Harmony mixture model components to estimate and regress out batch effects from the query data. Importantly, the reference cell embedding remains stable during this mapping process. We transferred annotation labels from the reference to the query dataset by exploiting cell proximity in the embedding space using nearest neighbors through s*klearn.classifier.KNeighborsClassifier*, Symphony default choice [https://github.com/potulabe/symphonypy].

#### scPoli

In contrast to other integration methods such as scANVI, scPoli^72^ encodes the condition (*chemistry*, *sampleID*) as a learnable conditional embedding and characterizes each cell-type as a prototype in the latent embedding to facilitate the label transfer. In the reference building phase, we first pre-trained the model given the reference dataset and its conditions, and then fine-tuned to optimize the prototypes. In the reference mapping phase, we freezed the model and learned the new conditional embeddings belonging to the query dataset. The label transfer is performed by simply assigning the cell type belonging to the closest prototype in the latent embedding space. All the methods belong to the scArches^66^ class *scarches.models.scPoli*.

Note that scGen and Harmony/Symphony approaches generate one integrated dataset that is independent from the query data, while scANVI and scPoli require a fine-tuning of the reference model for a given query dataset.

### Disease Classifiers

#### Patient Embeddings Definition

After obtaining the corrected embedding from one of the data integration approaches described previously, we need to aggregate the cell-wise embeddings into patient-wise embeddings. We decided to group at the level of the cell-types, by computing the mean embedding across cells belonging to the same cell type and *sampleID*. Only for scPoli, we generated three different types of patient embeddings: the learned patient embeddings (*sample*), the averaged cell-wise latent embeddings (*cell*), and the concatenation of the two (*cell&sample*).

#### Classifiers Definition and Hyperparameter Tuning

In this phase the aim is to train a classifier for each cell-type on the patient-wise embeddings belonging to the reference dataset. We tested the following classifier types: *sklearn.svm.LinearSVC*, *sklearn.svm.SVC*, and *sklearn.neighbors.KNeighborsClassifier (sklearn v.1.4.1.post1)*. For each classifier type, we trained different configurations defined in **Supplementary Table 7**, and evaluated their performance using a 5-fold cross-validation on the reference patient embeddings. Similarly to what we did to optimize the XGBoost classifier when estimating the immune gene importance, we employed the Optuna library to perform the hyperparameter tuning for each classifier. The best hyperparameters combination was selected according to the weighted F1 score independently of the cell-type.

#### Majority Voting and Evaluation

The best classifier type according to the average performance over all cell-type is then used to train from scratch the corresponding classifier on the whole reference patient embedding. The predicted condition for a patient is simply the majority voting among the classifiers. In case of a tie of different conditions, we conservatively rejected the prediction of the classifiers. Then, the overall metrics weighted F1-score, BAS, and Matthew Correlation Coefficient (MCC), and the disease-wise metrics Precision, Recall, BAS and F1-score were computed by comparing the predicted inflammatory conditions by each classifier type in the query dataset with the available ground truth. All those metrics were computed with the Sklearn python library, when we refer to the weighted version of a given metric we are using average=’weighted’ parameter to take into account the unbalance of the inflammatory condition observations.

Note, if a given query patient does not have any cells annotated for a given cell type, the corresponding prediction was set as ‘Not Available’. This label was not taken into account during the Majority Voting procedure, and considered as a wrong prediction when evaluating the performances of that cell-type.

