## Supplementary material for "Interpretable Inflammation Landscape of Circulating Immune cells": JimenezGracia_et_al_2023_SUPPLEMENTARYMATERIAL_v2

### Extended Data / Supplementary information

#### Extended Data Figure Legends (1-11)

**Extended Data Figure 1. Composition of the inflammation atlas datasets from the Main Core.** (**a**) Barplot showing patient count distribution across different diseases, including healthy condition, stratified by technical variables (*studyID* and *chemistry*) and clinical metadata (*age* and *sex*). The donor without age information is shown in white. (**b**) *SCIB* metric results computed on five different embedding spaces, ranked by their overall performances. (**c**) Heatmaps showing the coefficient of determination $R^2$ from a linear regression between each principal component and one of four confounding factors. The Principal Component Analysis was performed on (**left**) original data (normalized and log-scaled) and (**right**) from scANVI normalized expression (log-scaled). (**d**) Cellular proportions (*Level 1*) across diseases and healthy donors (**Top**). Compositional analysis of *Level 1* populations (excluding Platelets and RBC) between each disease and healthy donors (**Bottom**). The dot size reflects the significance of the result ('Final parameter' != 0), and the color represents the log2FC (Disease vs Healthy).

**Extended Data Figure 2. Marker gene expression for the Inflammation Atlas cell populations.** Dotplot showing the average expression of canonical gene markers (*x-axis*) characterizing each cell population (*y-axis*), both considering (**a**) Level1 and (**b**) Level2 annotation (UTC, Unconventional T cells[133]; ILC, Innate Lymphoid Cells[134]). The dot size reflects the percentage of cells in a cluster expressing each gene, and the color represents the average expression level.

**Extended Data Figure 3**. **Inflammation-related signatures across diseases and cell types**. (**a**) Agglomerative hierarchical clustering with complete linkage, performed considering the euclidean distance among columns, based on the corrected immune-related signature activity score computed by disease and cell type (*Level 1*). (**b**) Barplot showing factor scores for each disease obtained from MOFAcell analysis. (**c**) Boxplot displaying the activity of STA1 and SP1 across cell types (annotation *Level2*) in SLE patients from *Perez et al.*[27]. Asterisk (*) identifies significant differences with respect to other lineage populations (Wilcoxon rank-sum test, FDR Adjusted p-value < 0.05); location refers to the direction of change (on top, means upregulated whereas below, refers to downregulated). (**d**) Dotplot showing the uncorrected average expression of the 50 genes defining the IFN Type 1 and 2 signature (*x-axis*) in Non-Naive CD8 T cell population (*Level 1*) across IMID diseases and health (SCGT00 study, *y-axis*). (**e**) Dotplot showing the uncorrected average expression of the 10 top ranked genes defining the IFN Type 1 and 2 signature (*x-axis*) across subpopulation of Non-Naive CD8 T cells (*Level 2*) in IMID diseases and health (SCGT00 study). For panels (**d**) and (**e**), the dot size reflects the percentage of cells of each disease expressing each gene, and the color represents the average expression level.

**Extended Data Figure 4. Confusion matrices of predicted inflammatory condition by cell type.** Normalized confusion matrices, one for each cell-type, displaying proportion of predictions belonging to each True Condition. Diagonal values correspond to the Recall metric. XGBoost was trained on the (a) scANVI batch corrected and log-scaled cell expression profiles, and **(b)** original normalized and log-scaled cell expression profiles.

**Extended Data Figure 5. Functional biomarker discovery using interpretable machine learning analysis.** (**a**) Gene list ranked top-to-bottom by importance (absolute SHAP value), coupled with max-normalized expression levels computed per celltype (*Level1*) and considering selected diseases. From **left** to **right**, reporting top ranked genes for Naive CD4 T cells in RA disease as well as for Monocytes and pDC in SLE patients. (**b**) Rank by importance (absolute SHAP value) of the CYBA gene in every combination of celltype (*Level1*) and disease. (**c**) Scatter plot of max-normalized gene expression against SHAP values computed for *CYBA* gene on Monocyte population and specific diseases (COPD and Asthma, from left to right). (**d**) Rank by importance (absolute SHAP value) of *IFITM1* gene in every combination of celltype (*Level1*) and disease. (**e**) Scatter plot of max-normalized gene expression against SHAP values computed for *IFITM1* gene on ILC populations and disease (Cirrhosis, COPD, and Asthma, from **top-left**, **top-right**, and **bottom** respectively). In Panels **(a)**, **(b)**, and (**d**) we first dropped the genes expressed in less than 5% of the selected cell population. In Panels (**c**) and (**e**), we limited the visualization to up to 60,000 cells, sampling an equal percentage from each patient corresponding to 5% and 15% of the total cells for Monocytes and ILC, respectively. Cells belonging to samples with or without the given condition are marked in orange or blue, respectively.

**Extended Data Figure 6. Sample-wise distribution of SHAP-values for *CYBA* and *IFITM1*.** Averaged sample-wise SHAP values for each inflammatory condition, i.e., the mean SHAP value of the corresponding cells on the following combinations of cell-type and gene: (Mono, *CYBA*), (CD8 Non-Naive, *IFITM1*), (CD4 Non-Naive, *IFITM1*), and (ILC, *IFITM1*). The orange and blue groups correspond to the samples with or without the corresponding condition (disease), respectively.

**Extended Data Figure 7. Extended patient classifier workflow schema**. (**a**) Definition of reference and query datasets, for Scenarios 1, 2, 3, and centralized dataset (from left to right). (**b**) Integration of the reference dataset and mapping of the query dataset to define the patient-wise embeddings, stratified by cell-type. (**c**) Patient classifier pipeline composed by the hyperparameter tuning of each classifier family, the selection of the best classifier family and the final evaluation of the left-out query dataset. (d) Schema of the three experiments performed.

**Extended Data Figure 8. Additional performance evaluation metrics. Scenario 1.** (**a**) Boxplots showing the distribution of Balanced Accuracy Score (balanced by true disease support), F1, Precision and Recall computed during 5-fold cross validation, considering Majority Vote prediction on the left-out split for each inflammatory condition. The average number of samples among 5 splits, with the corresponding ground truth labels, are also reported. (**b**) Boxplots showing the distribution of Balance Accuracy Score (**top**), and Matthew Correlation Coefficient (**bottom**) computed during 5-fold cross validation, considering Majority Vote and cell-type prediction, on the left-out split. (**c**) Heatmap reporting Recall and Precision (from **left** to **right**) computed by aggregating the prediction performed by each cell-type on each left-out split during 5-fold cross validation.

**Extended Data Figure 9. Additional performance evaluation metric in Scenario 2 and 3.** (**a**) Table with all the data integration method configurations and corresponding performance metrics computed on data from unseen patients (Scenario 2) and unseen studies (Scenario 3). The best performing classifiers (SVM with different kernels and kNN) in terms of Weighted F1 are reported. (**b-c**) Performance evaluation from Scenario 2 and 3, respectively, showing (**left**) the distribution of Weighted Recall and Weighted Precision for all the configurations of each data integration approach, and (**right**) the mean and standard-deviation of each data integration method, including 100 random label assignments. Arrows highlight the scANVI configuration applied in Scenario 1.

**Extended Data Figure 10. Additional performance evaluation metric in the Centralized dataset analysis.** (**a**) Pointplot showing the Balance Accuracy Score (**top**), and Matthew Correlation Coefficient (**bottom**) computed, considering Majority Vote, 100 random disease assignments, and cell-type prediction, on the samples from left out pools. (**b**) Heatmap reporting Recall and Precision obtained on the samples from left out pools by each cell-type for each disease included in the centralized dataset.

**Extended Data Figure 11. Patient classifier performance in Scenarios 2 and 3. (a,c,e,g)** Result obtained with the best parameter configuration for each integration and mapping method, considering W-F1 score computed on prediction of samples from unseen patients. **(b,d,f,h)** Result obtained with the best parameter configuration for each integration and mapping method, considering W-F1 score computed on prediction of samples from unseen studies. In Panels (**a**) to (**h**): (**top-left**) Pointplot of weighted F1-scores for Majority vote and each cell-type. (**bottom-left**) F1-score for each combination of cell-type and disease, columns ordered for similarities. (**right**) Normalized confusion matrices displaying proportion of predictions belonging to each true condition. Diagonal values correspond to the Recall metric. Corresponding Majority Vote Weighted F1 score (WF1) and Balanced Accuracy Score (BAS) were reported. Note, scPoli configurations where embedding space = *sample* were not considered.

Ext F1

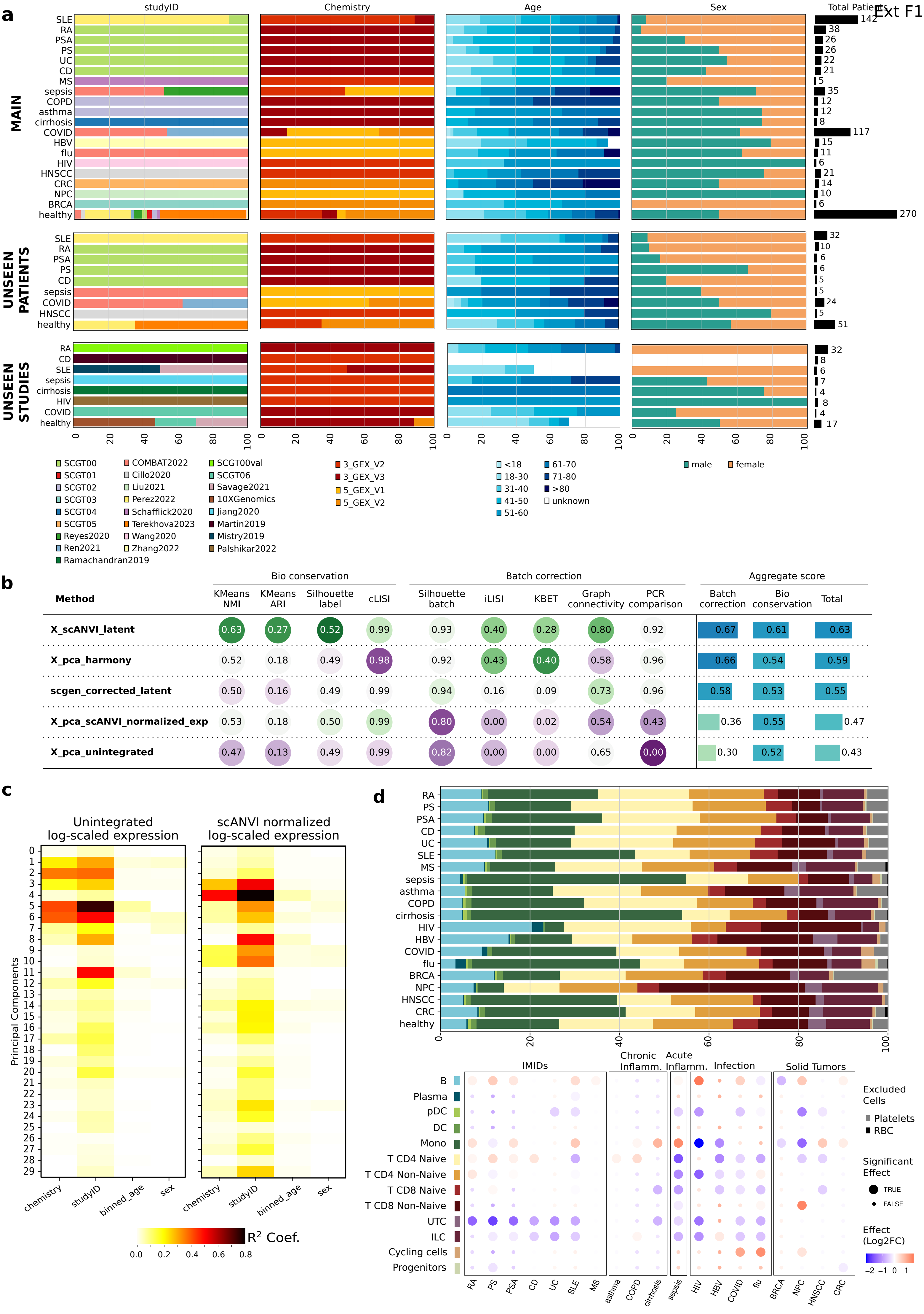

a
studyID
Chemistry
Age
Sex
Total Patients
MAIN
UNSEEN PATIENTS
UNSEEN STUDIES
SLE RA PSA PS UC CD MS sepsis COPD asthma cirrhosis COVID HBV flu HIV HNSCC CRC NPC BRCA healthy
142 38 26 26 22 21 5 35 12 12 8 117 15 11 6 21 14 10 6 270
32 10 6 6 5 5 24 5 51
32 8 6 7 4 8 4 17
SCGT00 SCGT01 SCGT02 SCGT03 SCGT04 SCGT05 Reyes2020 Ren2021 Ramachandran2019
COMBAT2022 Cillo2020 Liu2021 Perez2022 Schafflick2020 Terekhova2023 Wang2020 Zhang2022
SCGT00val SCGT06 Savage2021 10XGenomics Jiang2020 Martin2019 Mistry2019 Palshikar2022
3_GEX_V2 3_GEX_V3 5_GEX_V1 5_GEX_V2
<18 18-30 31-40 41-50 51-60 61-70 71-80 >80 unknown
male female
b
Method
Bio conservation: KMeans NMI, KMeans ARI, Silhouette label, cLISI
Batch correction: Silhouette batch, iLISI, KBET, Graph connectivity, PCR comparison
Aggregate score: Batch correction, Bio conservation, Total
X_scANVI_latent 0.63 0.27 0.52 0.99 0.93 0.40 0.28 0.80 0.92 0.67 0.61 0.63
X_pca_harmony 0.52 0.18 0.49 0.98 0.92 0.43 0.40 0.58 0.96 0.66 0.54 0.59
scgen_corrected_latent 0.50 0.16 0.49 0.99 0.94 0.16 0.09 0.73 0.96 0.58 0.53 0.55
X_pca_scANVI_normalized_exp 0.53 0.18 0.50 0.99 0.80 0.00 0.02 0.54 0.43 0.36 0.55 0.47
X_pca_unintegrated 0.47 0.13 0.49 0.99 0.82 0.00 0.00 0.65 0.00 0.30 0.52 0.43
c
Unintegrated log-scaled expression
scANVI normalized log-scaled expression
Principal Components
chemistry studyID binned_age sex
$R^2$ Coef.
0.0 0.2 0.4 0.6 0.8
d
IMIDs
Chronic Inflamm.
Acute Inflamm.
Infection
Solid Tumors
B Plasma pDC DC Mono T CD4 Naive T CD4 Non-Naive T CD8 Naive T CD8 Non-Naive UTC ILC Cycling cells Progenitors
Excluded Cells
Platelets
RBC
Significant Effect
TRUE
FALSE
Effect (Log2FC)
−2 −1 0 1

Ext F2

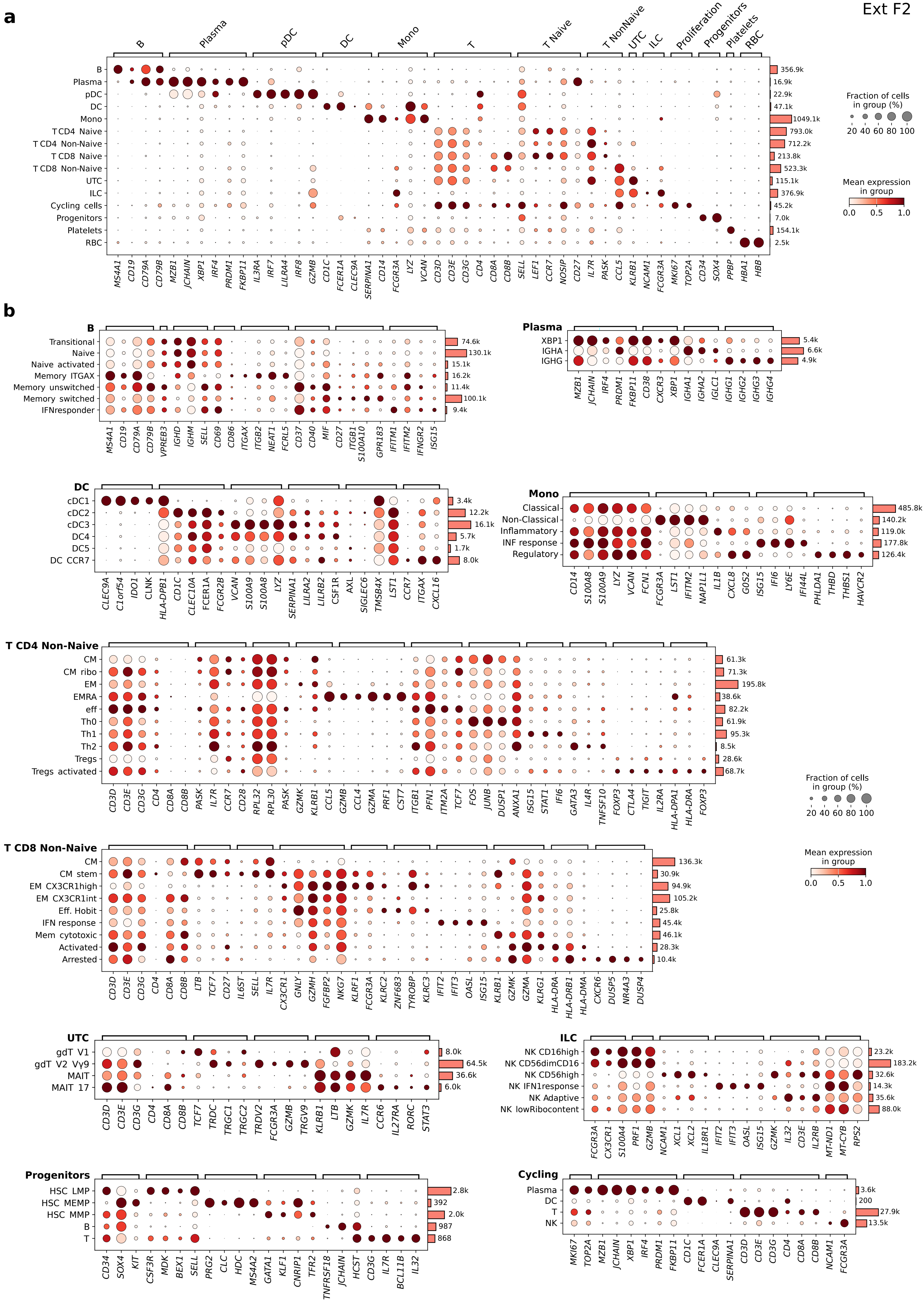

a
B Plasma pDC DC Mono T T Naive T NonNaive UTC ILC Proliferation Progenitors Platelets RBC
B
Plasma
pDC
DC
Mono
T CD4 Naive
T CD4 Non-Naive
T CD8 Naive
T CD8 Non-Naive
UTC
ILC
Cycling cells
Progenitors
Platelets
RBC
356.9k 16.9k 22.9k 47.1k 1049.1k 793.0k 712.2k 213.8k 523.3k 115.1k 376.9k 45.2k 7.0k 154.1k 2.5k
MS4A1 CD19 CD79A CD79B MZB1 JCHAIN XBP1 IRF4 PRDM1 FKBP11 IL3RA IRF7 LILRA4 IRF8 GZMB CD1C FCER1A CLEC9A SERPINA1 CD14 FCGR3A LYZ VCAN CD3D CD3E CD3G CD4 CD8A CD8B SELL LEF1 CCR7 NOSIP CD27 IL7R PASK CCL5 KLRB1 NCAM1 FCGR3A MKI67 TOP2A CD34 SOX4 PPBP HBA1 HBB
Fraction of cells in group (%)
20 40 60 80 100
Mean expression in group
0.0 0.5 1.0
b
B
Transitional
Naive
Naive activated
Memory ITGAX
Memory unswitched
Memory switched
IFNresponder
74.6k 130.1k 15.1k 16.2k 11.4k 100.1k 9.4k
MS4A1 CD19 CD79A CD79B VPREB3 IGHD IGHM SELL CD69 CD86 ITGAX ITGB2 NEAT1 FCRL5 CD37 CD40 MIF CD27 ITGB1 S100A10 GPR183 IFITM1 IFITM2 IFNGR2 ISG15
Plasma
XBP1
IGHA
IGHG
5.4k 6.6k 4.9k
MZB1 JCHAIN IRF4 PRDM1 FKBP11 CD38 CXCR3 XBP1 IGHA1 IGHA2 IGLC1 IGHG1 IGHG2 IGHG3 IGHG4
DC
cDC1
cDC2
cDC3
DC4
DC5
DC CCR7
3.4k 12.2k 16.1k 5.7k 1.7k 8.0k
CLEC9A C1orf54 IDO1 CLNK HLA-DPB1 CD1C CLEC10A FCER1A FCGR2B VCAN S100A9 S100A8 LYZ SERPINA1 LILRA2 LILRB2 CSF1R AXL SIGLEC6 TMSB4X LST1 CCR7 ITGAX CXCL16
Mono
Classical
Non-Classical
Inflammatory
INF response
Regulatory
485.8k 140.2k 119.0k 177.8k 126.4k
CD14 S100A8 S100A9 LYZ VCAN FCN1 FCGR3A LST1 IFITM2 NAP1L1 IL1B CXCL8 G0S2 ISG15 IFI6 LY6E IFI44L PHLDA1 THBD THBS1 HAVCR2
T CD4 Non-Naive
CM
CM ribo
EM
EMRA
eff
Th0
Th1
Th2
Tregs
Tregs activated
61.3k 71.3k 195.8k 38.6k 82.2k 61.9k 95.3k 8.5k 28.6k 68.7k
CD3D CD3E CD3G CD4 CD8A CD8B PASK IL7R CCR7 CD28 RPL32 RPL30 PASK GZMK KLRB1 CCL5 GZMB CCL4 GZMA PRF1 CST7 ITGB1 PFN1 ITM2A TCF7 FOS JUNB DUSP1 ANXA1 ISG15 STAT1 IFI6 GATA3 IL4R TNFSF10 FOXP3 CTLA4 TIGIT IL2RA HLA-DPA1 HLA-DRA FOXP3
Fraction of cells in group (%)
20 40 60 80 100
T CD8 Non-Naive
CM
CM stem
EM CX3CR1high
EM CX3CR1int
Eff. Hobit
IFN response
Mem cytotoxic
Activated
Arrested
136.3k 30.9k 94.9k 105.2k 25.8k 45.4k 46.1k 28.3k 10.4k
CD3D CD3E CD3G CD4 CD8A CD8B LTB TCF7 CD27 IL6ST SELL IL7R CX3CR1 GNLY GZMH FGFBP2 NKG7 KLRF1 FCGR3A KLRC2 ZNF683 TYROBP KLRC3 IFIT2 IFIT3 OASL ISG15 KLRB1 GZMK GZMA KLRG1 HLA-DRA HLA-DRB1 HLA-DMA CXCR6 DUSP5 NR4A3 DUSP4
Mean expression in group
0.0 0.5 1.0
UTC
gdT V1
gdT V2 Vγ9
MAIT
MAIT 17
8.0k 64.5k 36.6k 6.0k
CD3D CD3E CD3G CD4 CD8A CD8B TCF7 TRDC TRGC1 TRGC2 TRDV2 FCGR3A GZMB TRGV9 KLRB1 LTB GZMK IL7R CCR6 IL27RA RORC STAT3
ILC
NK CD16high
NK CD56dimCD16
NK CD56high
NK IFN1response
NK Adaptive
NK lowRibocontent
23.2k 183.2k 32.6k 14.3k 35.6k 88.0k
FCGR3A CX3CR1 S100A4 PRF1 GZMB NCAM1 XCL1 XCL2 IL18R1 IFIT2 IFIT3 OASL ISG15 GZMK IL32 CD3E IL2RB MT-ND1 MT-CYB RPS2
Progenitors
HSC LMP
HSC MEMP
HSC MMP
B
T
2.8k 392 2.0k 987 868
CD34 SOX4 KIT CSF3R MDK BEX1 SELL PRG2 CLC HDC MS4A2 GATA1 KLF1 CNRIP1 TFR2 TNFRSF18 JCHAIN HCST CD3G IL7R BCL11B IL32
Cycling
Plasma
DC
T
NK
3.6k 200 27.9k 13.5k
MKI67 TOP2A MZB1 JCHAIN XBP1 IRF4 PRDM1 FKBP11 CD1C FCER1A CLEC9A SERPINA1 CD3D CD3E CD3G CD4 CD8A CD8B NCAM1 FCGR3A

Ext F3

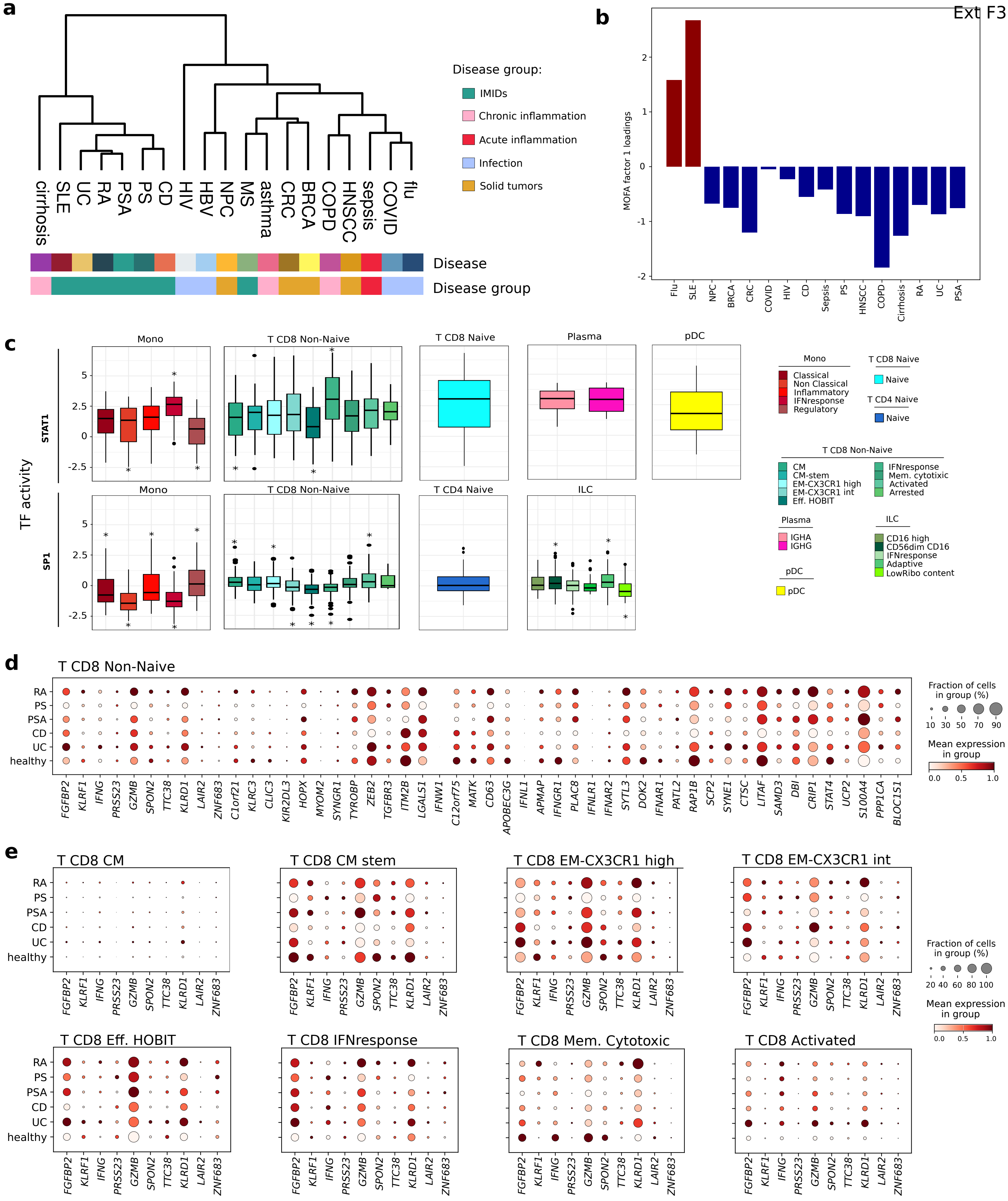
a
b
c
d
e
Disease group:
IMIDs
Chronic inflammation
Acute inflammation
Infection
Solid tumors
cirrhosis
SLE
UC
RA
PSA
PS
CD
HIV
HBV
NPC
MS
asthma
CRC
BRCA
COPD
HNSCC
sepsis
COVID
flu
Disease
Disease group
MOFA factor 1 loadings
Flu
SLE
NPC
BRCA
CRC
COVID
HIV
CD
Sepsis
PS
HNSCC
COPD
Cirrhosis
RA
UC
PSA
TF activity
STAT1
SP1
Mono
T CD8 Non-Naive
T CD8 Naive
Plasma
pDC
T CD4 Naive
ILC
Classical
Non Classical
Inflammatory
IFNresponse
Regulatory
Naive
CM
CM-stem
EM-CX3CR1 high
EM-CX3CR1 int
Eff. HOBIT
Mem. cytotoxic
Activated
Arrested
IGHA
IGHG
CD16 high
CD56dim CD16
Adaptive
LowRibo content
T CD8 Non-Naive
RA
PS
PSA
CD
UC
healthy
FGFBP2
KLRF1
IFNG
PRSS23
GZMB
SPON2
TTC38
KLRD1
LAIR2
ZNF683
C1orf21
KLRC3
CLIC3
KIR2DL3
HOPX
MYOM2
SYNGR1
TYROBP
ZEB2
TGFBR3
ITM2B
LGALS1
IFNW1
C12orf75
MATK
CD63
APOBEC3G
IFNL1
APMAP
IFNGR1
PLAC8
IFNLR1
IFNAR2
SYTL3
DOK2
IFNAR1
PATL2
RAP1B
SCP2
SYNE1
CTSC
LITAF
SAMD3
DBI
CRIP1
STAT4
UCP2
S100A4
PPP1CA
BLOC1S1
Fraction of cells
in group (%)
Mean expression
in group
T CD8 CM
T CD8 CM stem
T CD8 EM-CX3CR1 high
T CD8 EM-CX3CR1 int
T CD8 Eff. HOBIT
T CD8 IFNresponse
T CD8 Mem. Cytotoxic
T CD8 Activated

Ext F4

a

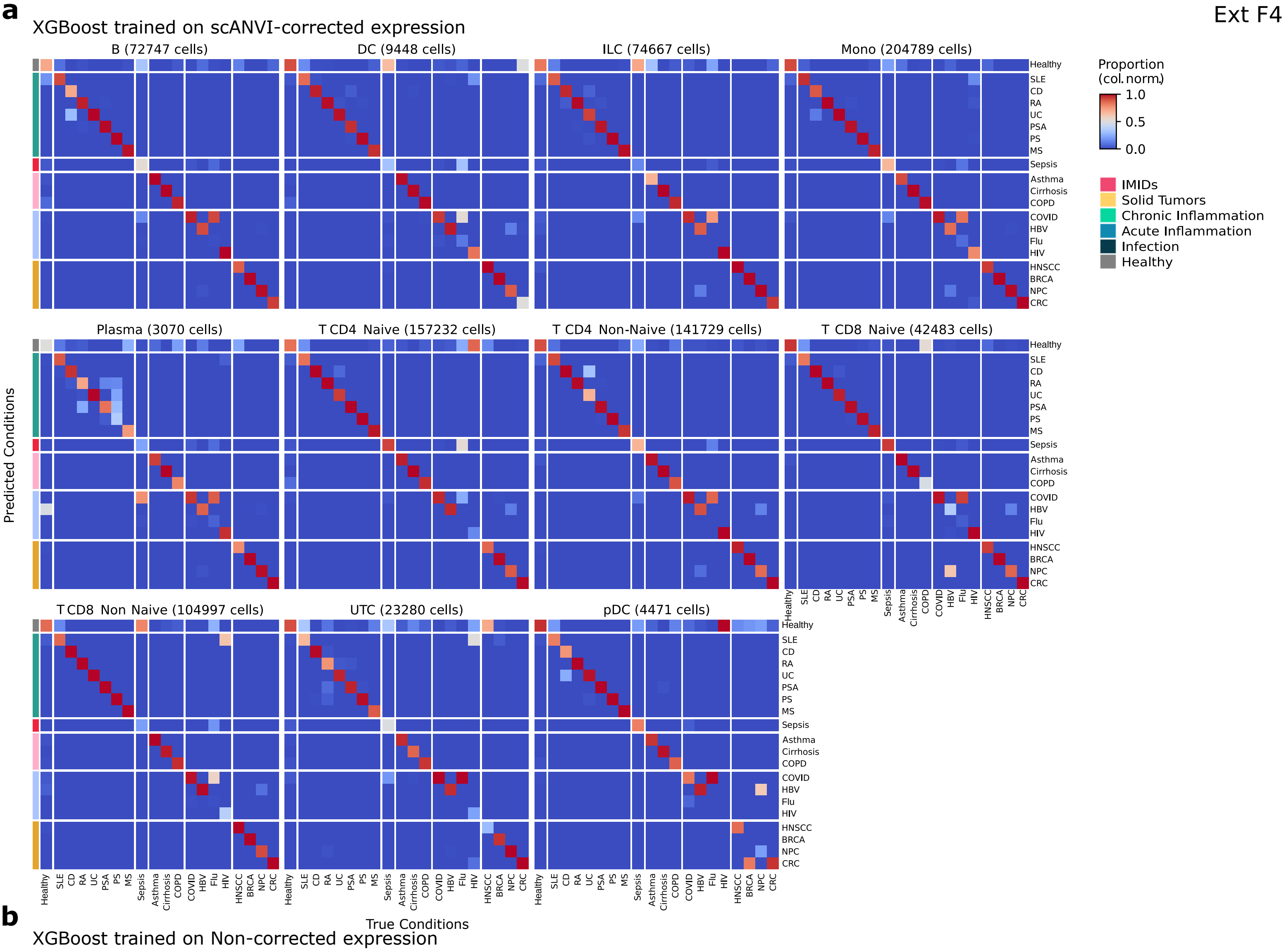
XGBoost trained on scANVI-corrected expression
B (72747 cells)
DC (9448 cells)
ILC (74667 cells)
Mono (204789 cells)
Plasma (3070 cells)
T CD4 Naive (157232 cells)
T CD4 Non-Naive (141729 cells)
T CD8 Naive (42483 cells)
T CD8 Non Naive (104997 cells)
UTC (23280 cells)
pDC (4471 cells)
Predicted Conditions
True Conditions
Healthy
SLE
CD
RA
UC
PSA
PS
MS
Sepsis
Asthma
Cirrhosis
COPD
COVID
HBV
Flu
HIV
HNSCC
BRCA
NPC
CRC
Proportion
(col. norm.)
1.0
0.5
0.0
IMIDs
Solid Tumors
Chronic Inflammation
Acute Inflammation
Infection
Healthy

b

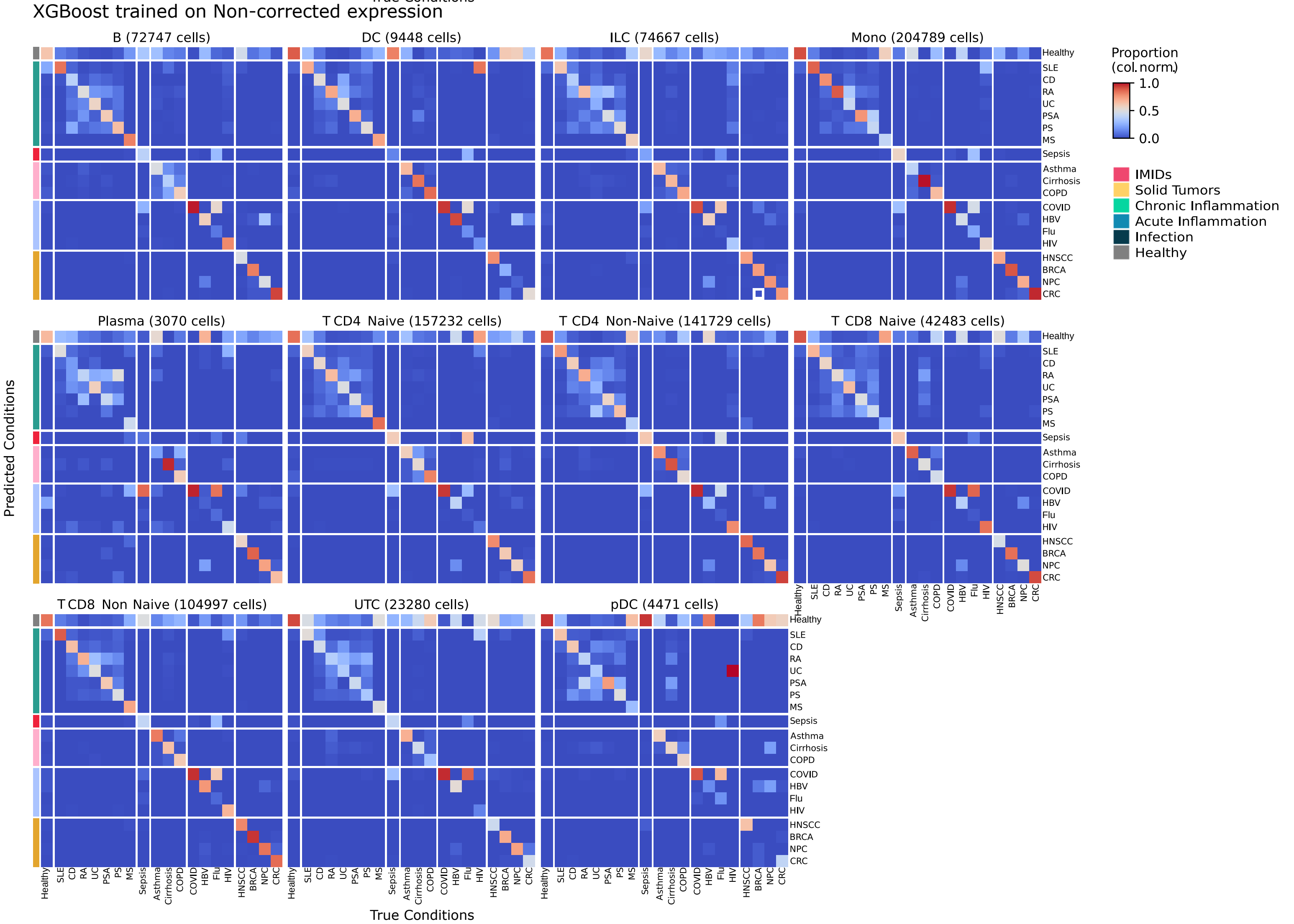
XGBoost trained on Non-corrected expression
B (72747 cells)
DC (9448 cells)
ILC (74667 cells)
Mono (204789 cells)
Plasma (3070 cells)
T CD4 Naive (157232 cells)
T CD4 Non-Naive (141729 cells)
T CD8 Naive (42483 cells)
T CD8 Non Naive (104997 cells)
UTC (23280 cells)
pDC (4471 cells)
Predicted Conditions
True Conditions
Healthy
SLE
CD
RA
UC
PSA
PS
MS
Sepsis
Asthma
Cirrhosis
COPD
COVID
HBV
Flu
HIV
HNSCC
BRCA
NPC
CRC
Proportion
(col. norm.)
1.0
0.5
0.0
IMIDs
Solid Tumors
Chronic Inflammation
Acute Inflammation
Infection
Healthy

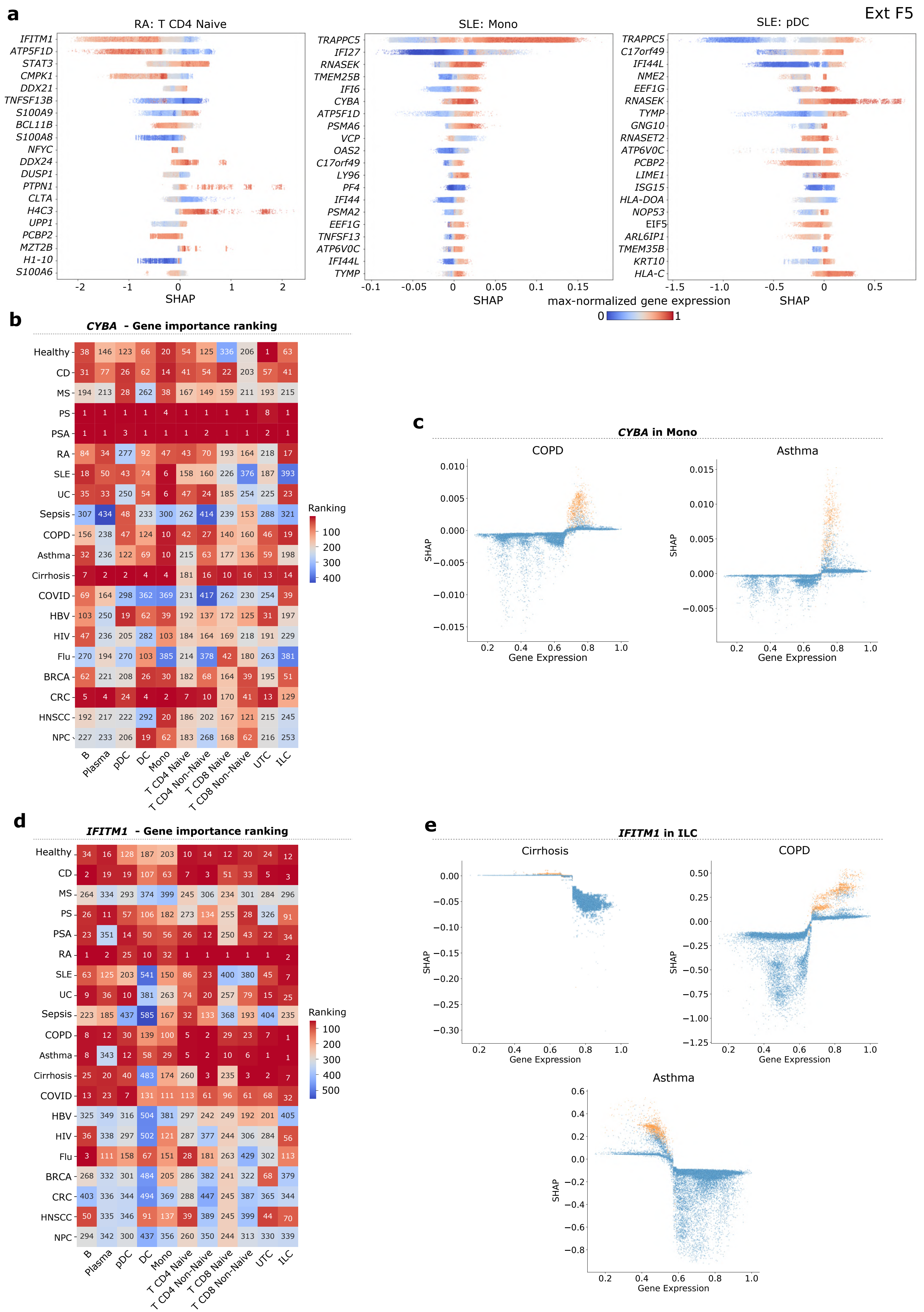

Ext F5
a
RA: T CD4 Naive
IFITM1
ATP5F1D
STAT3
CMPK1
DDX21
TNFSF13B
S100A9
BCL11B
S100A8
NFYC
DDX24
DUSP1
PTPN1
CLTA
H4C3
UPP1
PCBP2
MZT2B
H1-10
S100A6
-2 -1 0 1 2
SHAP
SLE: Mono
TRAPPC5
IFI27
RNASEK
TMEM25B
IFI6
CYBA
ATP5F1D
PSMA6
VCP
OAS2
C17orf49
LY96
PF4
IFI44
PSMA2
EEF1G
TNFSF13
ATP6V0C
IFI44L
TYMP
-0.1 -0.05 0 0.05 0.1 0.15
SHAP
SLE: pDC
TRAPPC5
C17orf49
IFI44L
NME2
EEF1G
RNASEK
TYMP
GNG10
RNASET2
ATP6V0C
PCBP2
LIME1
ISG15
HLA-DOA
NOP53
EIF5
ARL6IP1
TMEM35B
KRT10
HLA-C
-1.5 -1.0 -0.5 0 0.5
SHAP
max-normalized gene expression
0 1
b
CYBA - Gene importance ranking
Healthy 38 146 123 66 20 54 125 336 206 1 63
CD 31 77 26 62 14 41 54 22 203 57 41
MS 194 213 28 262 38 167 149 159 211 193 215
PS 1 1 1 1 4 1 1 1 1 8 1
PSA 1 1 3 1 1 1 2 1 1 2 1
RA 84 34 277 92 47 43 70 193 164 218 17
SLE 18 50 43 74 6 158 160 226 376 187 393
UC 35 33 250 54 6 47 24 185 254 225 23
Sepsis 307 434 48 233 300 262 414 239 153 288 321
COPD 156 238 47 124 10 42 27 140 160 46 19
Asthma 32 236 122 69 10 215 63 177 136 59 198
Cirrhosis 7 2 2 4 4 181 16 10 16 13 14
COVID 69 164 298 362 369 231 417 262 230 254 39
HBV 103 250 19 62 39 192 137 172 125 31 197
HIV 47 236 205 282 103 184 164 169 218 191 229
Flu 270 194 270 103 385 214 378 42 180 263 381
BRCA 62 221 208 26 30 182 68 164 39 195 51
CRC 5 4 24 4 2 7 10 170 41 13 129
HNSCC 192 217 222 292 20 186 202 167 121 215 245
NPC 227 233 206 19 62 183 268 168 62 216 253
B Plasma pDC DC Mono T CD4 Naive T CD4 Non-Naive T CD8 Naive T CD8 Non-Naive UTC ILC
Ranking
100
200
300
400
c
CYBA in Mono
COPD
0.010
0.005
0.000
−0.005
−0.010
−0.015
SHAP
0.2 0.4 0.6 0.8 1.0
Gene Expression
Asthma
0.015
0.010
0.005
0.000
−0.005
SHAP
0.2 0.4 0.6 0.8 1.0
Gene Expression
d
IFITM1 - Gene importance ranking
Healthy 34 16 128 187 203 10 14 12 20 24 12
CD 2 19 19 107 63 7 3 51 33 5 3
MS 264 334 293 374 399 245 306 234 301 284 296
PS 26 11 57 106 182 273 134 255 28 326 91
PSA 23 351 14 50 56 26 12 250 43 22 34
RA 1 2 25 10 32 1 1 1 1 1 2
SLE 63 125 203 541 150 86 23 400 380 45 7
UC 9 36 10 381 263 74 20 257 79 15 25
Sepsis 223 185 437 585 167 32 133 368 193 404 235
COPD 8 12 30 139 100 5 2 29 23 7 1
Asthma 8 343 12 58 29 5 2 10 6 1 1
Cirrhosis 25 20 40 483 174 260 3 235 3 2 7
COVID 13 23 7 131 111 113 61 96 61 68 32
HBV 325 349 316 504 381 297 242 249 192 201 405
HIV 36 338 297 502 121 287 377 244 306 284 56
Flu 3 111 158 67 151 28 181 263 429 302 113
BRCA 268 332 301 484 205 286 382 241 322 68 379
CRC 403 336 344 494 369 288 447 245 387 365 344
HNSCC 50 335 346 91 137 39 389 245 399 44 70
NPC 294 342 300 437 356 260 350 244 313 330 339
B Plasma pDC DC Mono T CD4 Naive T CD4 Non-Naive T CD8 Naive T CD8 Non-Naive UTC ILC
Ranking
100
200
300
400
500
e
IFITM1 in ILC
Cirrhosis
0.00
−0.05
−0.10
−0.15
−0.20
−0.25
−0.30
SHAP
0.2 0.4 0.6 0.8 1.0
Gene Expression
COPD
0.50
0.25
0.00
−0.25
−0.50
−0.75
−1.00
−1.25
SHAP
0.2 0.4 0.6 0.8 1.0
Gene Expression
Asthma
0.6
0.4
0.2
0.0
−0.2
−0.4
−0.6
−0.8
SHAP
0.2 0.4 0.6 0.8 1.0
Gene Expression

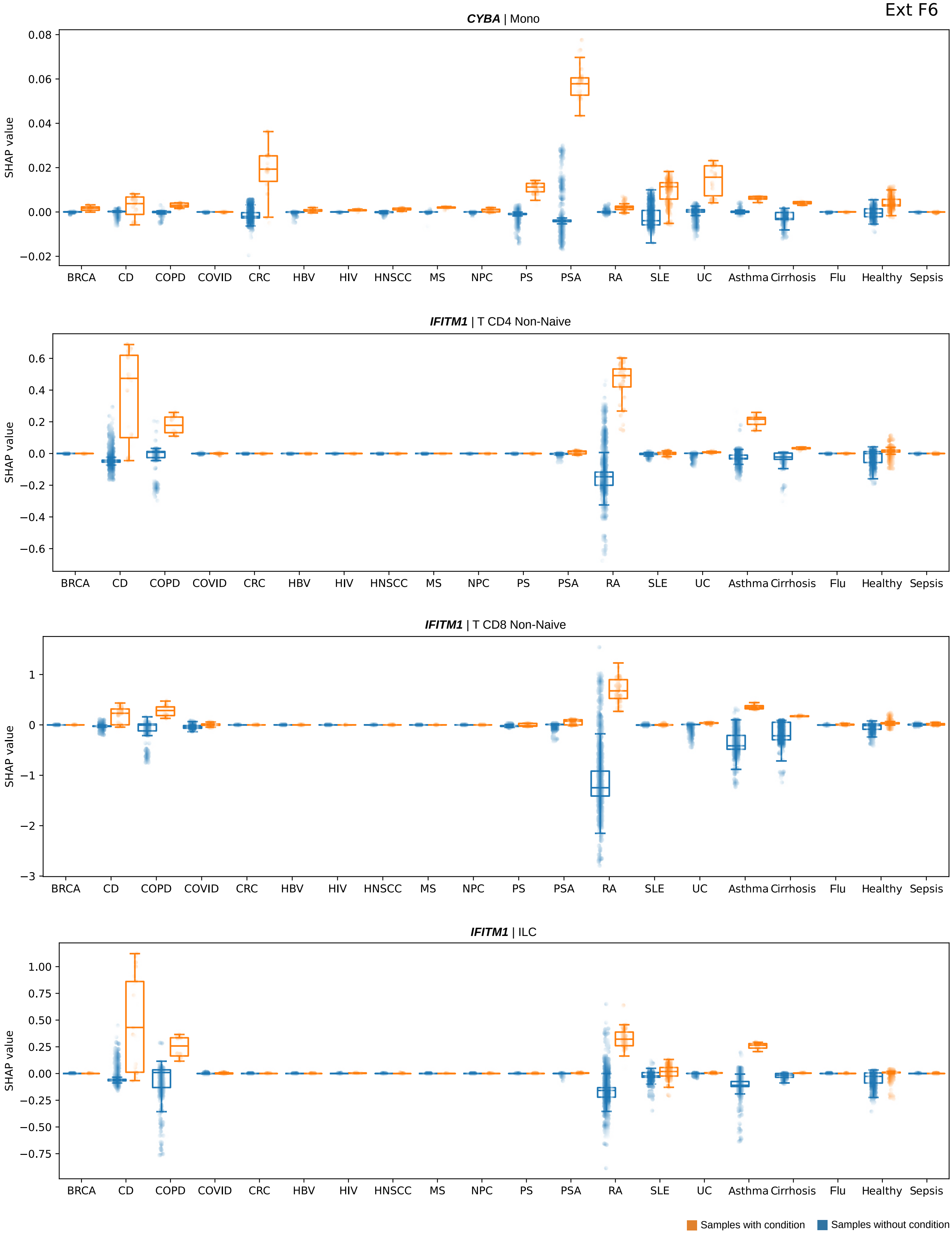
CYBA | Mono
SHAP value
0.08
0.06
0.04
0.02
0.00
−0.02
IFITM1 | T CD4 Non-Naive
SHAP value
0.6
0.4
0.2
0.0
−0.2
−0.4
−0.6
IFITM1 | T CD8 Non-Naive
SHAP value
1
0
−1
−2
−3
IFITM1 | ILC
SHAP value
1.00
0.75
0.50
0.25
0.00
−0.25
−0.50
−0.75
BRCA
CD
COPD
COVID
CRC
HBV
HIV
HNSCC
MS
NPC
PS
PSA
RA
SLE
UC
Asthma
Cirrhosis
Flu
Healthy
Sepsis
Samples with condition
Samples without condition

Ext F7

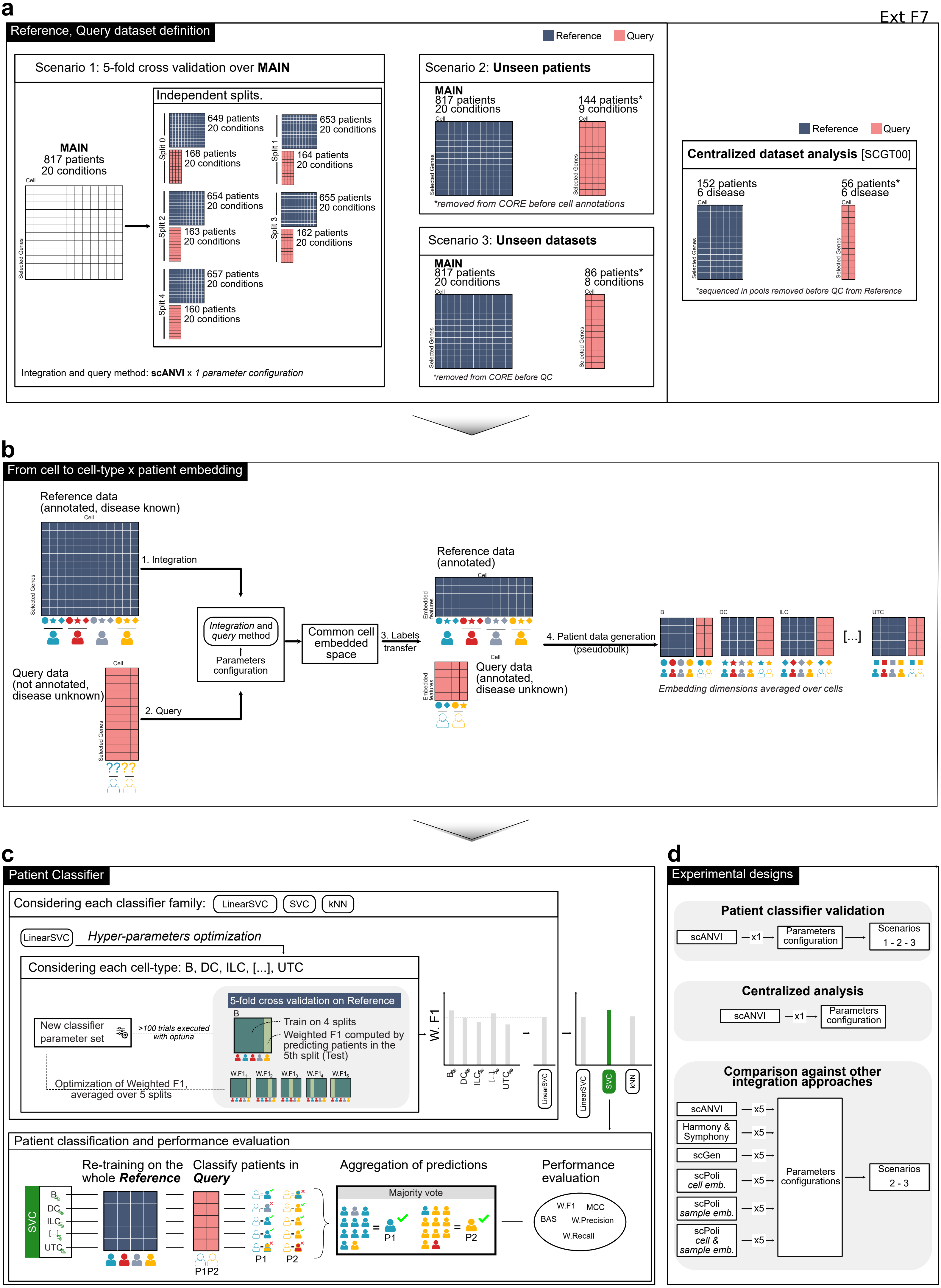
a
Reference, Query dataset definition
Reference
Query
Scenario 1: 5-fold cross validation over MAIN
MAIN
817 patients
20 conditions
Cell
Selected Genes
Independent splits.
Split 0
649 patients
20 conditions
168 patients
20 conditions
Split 1
653 patients
20 conditions
164 patients
20 conditions
Split 2
654 patients
20 conditions
163 patients
20 conditions
Split 3
655 patients
20 conditions
162 patients
20 conditions
Split 4
657 patients
20 conditions
160 patients
20 conditions
Integration and query method: scANVI x 1 parameter configuration
Scenario 2: Unseen patients
MAIN
817 patients
20 conditions
144 patients*
9 conditions
*removed from CORE before cell annotations
Scenario 3: Unseen datasets
MAIN
817 patients
20 conditions
86 patients*
8 conditions
*removed from CORE before QC
Centralized dataset analysis [SCGT00]
152 patients
6 disease
56 patients*
6 disease
*sequenced in pools removed before QC from Reference
b
From cell to cell-type x patient embedding
Reference data
(annotated, disease known)
1. Integration
Integration and query method
Parameters configuration
Common cell embedded space
3. Labels transfer
Query data
(not annotated, disease unknown)
2. Query
Reference data
(annotated)
Embedded features
Query data
(annotated, disease unknown)
4. Patient data generation (pseudobulk)
B
DC
ILC
UTC
[...]
Embedding dimensions averaged over cells
c
Patient Classifier
Considering each classifier family:
LinearSVC
SVC
kNN
Hyper-parameters optimization
Considering each cell-type: B, DC, ILC, [...], UTC
New classifier parameter set
>100 trials executed with optuna
5-fold cross validation on Reference
Train on 4 splits
Weighted F1 computed by predicting patients in the 5th split (Test)
Optimization of Weighted F1, averaged over 5 splits
W. F1
Patient classification and performance evaluation
Re-training on the whole Reference
Classify patients in Query
Aggregation of predictions
Majority vote
P1
P2
P1P2
Performance evaluation
W.F1
MCC
BAS
W.Precision
W.Recall
d
Experimental designs
Patient classifier validation
scANVI
x1
Parameters configuration
Scenarios 1 - 2 - 3
Centralized analysis
scANVI
x1
Parameters configuration
Comparison against other integration approaches
scANVI
Harmony & Symphony
scGen
scPoli cell emb.
scPoli sample emb.
scPoli cell & sample emb.
x5
Parameters configurations
Scenarios 2 - 3

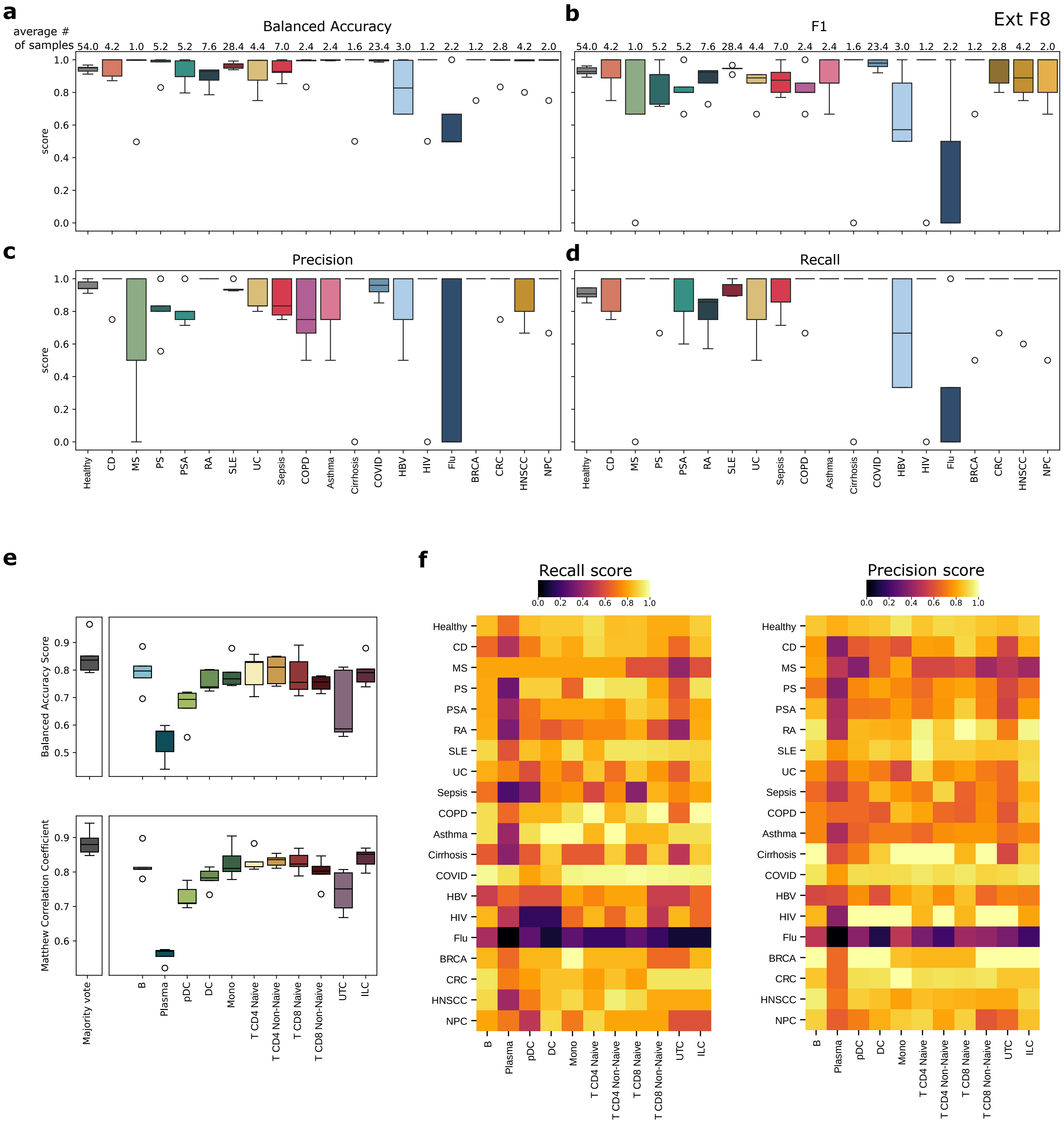
Ext F8
a
Balanced Accuracy
average # of samples
54.0 4.2 1.0 5.2 5.2 7.6 28.4 4.4 7.0 2.4 2.4 1.6 23.4 3.0 1.2 2.2 1.2 2.8 4.2 2.0
score
b
F1
c
Precision
d
Recall
Healthy
CD
MS
PS
PSA
RA
SLE
UC
Sepsis
COPD
Asthma
Cirrhosis
COVID
HBV
HIV
Flu
BRCA
CRC
HNSCC
NPC
e
Balanced Accuracy Score
Matthew Correlation Coefficient
Majority vote
B
Plasma
pDC
DC
Mono
T CD4 Naive
T CD4 Non-Naive
T CD8 Naive
T CD8 Non-Naive
UTC
ILC
f
Recall score
Precision score

a

| Method | Emb Space | Emb Dim | PEmb Dim | annot | wF1 | | BAS | | MCC | | wPRE | | wREC | | wSPE | | bestClf | |
|---|---|---|---|---|---|---|---|---|---|---|---|---|---|---|---|---|---|---|
| | | | | | Un. p. | Un. s. | Un. p. | Un. s. | Un. p. | Un. s. | Un. p. | Un. s. | Un. p. | Un. s. | Un. p. | Un. s. | Un. p. | Un. s. |
| **HS** | **cell** | **20** | **0** | **NU** | 0.84 | 0.47 | 0.69 | 0.24 | 0.8 | 0.38 | 0.87 | 0.46 | 0.84 | 0.5 | 0.97 | 0.93 | SVC | SVC |
| **scPoli** | **cellSample** | **200** | **100** | **1&2** | 0.66 | 0.46 | 0.42 | 0.24 | 0.62 | 0.45 | 0.64 | 0.42 | 0.71 | 0.55 | 0.89 | 0.89 | SVC | SVC |
| **scPoli** | **sample** | **200** | **100** | **1&2** | 0.6 | 0.46 | 0.36 | 0.24 | 0.55 | 0.46 | 0.6 | 0.42 | 0.65 | 0.56 | 0.86 | 0.9 | SVC | SVC |
| **HS** | **cell** | **30** | **0** | **NU** | 0.81 | 0.45 | 0.69 | 0.24 | 0.76 | 0.37 | 0.88 | 0.47 | 0.81 | 0.48 | 0.93 | 0.93 | SVC | SVC |
| **HS** | **cell** | **50** | **0** | **NU** | 0.86 | 0.42 | 0.76 | 0.24 | 0.83 | 0.34 | 0.9 | 0.47 | 0.86 | 0.44 | 0.95 | 0.92 | SVC | SVC |
| **scPoli** | **cell** | **100** | **20** | **1&2** | 0.96 | 0.41 | 0.95 | 0.18 | 0.95 | 0.27 | 0.97 | 0.49 | 0.96 | 0.36 | 0.99 | 0.96 | LSVC | LSVC |
| **scPoli** | **cell** | **200** | **100** | **1&2** | 0.94 | 0.4 | 0.88 | 0.16 | 0.92 | 0.24 | 0.96 | 0.55 | 0.94 | 0.34 | 0.99 | 0.97 | LSVC | LSVC |
| **HS** | **cell** | **200** | **0** | **NU** | 0.94 | 0.37 | 0.92 | 0.18 | 0.9 | 0.25 | 0.96 | 0.46 | 0.92 | 0.35 | 0.98 | 0.95 | LSVC | LSVC |
| **scPoli** | **sample** | **20** | **100** | **1&2** | 0.59 | 0.36 | 0.44 | 0.18 | 0.53 | 0.26 | 0.66 | 0.43 | 0.63 | 0.38 | 0.85 | 0.9 | SVC | SVC |
| **scPoli** | **cell** | **50** | **20** | **1&2** | 0.94 | 0.36 | 0.9 | 0.15 | 0.9 | 0.21 | 0.97 | 0.52 | 0.92 | 0.28 | 0.99 | 0.97 | LSVC | LSVC |
| **scPoli** | **sample** | **100** | **100** | **1&2** | 0.63 | 0.35 | 0.5 | 0.18 | 0.56 | 0.28 | 0.71 | 0.44 | 0.65 | 0.38 | 0.87 | 0.9 | SVC | SVC |
| **scGen** | **cell** | **20** | **0** | **2** | 0.96 | 0.34 | 0.91 | 0.15 | 0.94 | 0.23 | 0.97 | 0.45 | 0.95 | 0.33 | 0.99 | 0.94 | SVC | SVC |
| **HS** | **cell** | **100** | **0** | **NU** | 0.85 | 0.33 | 0.78 | 0.19 | 0.8 | 0.25 | 0.9 | 0.46 | 0.84 | 0.35 | 0.94 | 0.92 | SVC | SVC |
| **scPoli** | **cell** | **50** | **100** | **1&2** | 0.97 | 0.32 | 0.96 | 0.11 | 0.95 | 0.17 | 0.99 | 0.49 | 0.96 | 0.24 | 0.99 | 0.98 | LSVC | LSVC |
| **scPoli** | **cell** | **30** | **20** | **1&2** | 0.92 | 0.32 | 0.84 | 0.13 | 0.89 | 0.18 | 0.95 | 0.47 | 0.91 | 0.24 | 0.99 | 0.98 | LSVC | LSVC |
| **scPoli** | **cellSample** | **20** | **100** | **1&2** | 0.64 | 0.31 | 0.49 | 0.14 | 0.57 | 0.19 | 0.68 | 0.43 | 0.66 | 0.3 | 0.88 | 0.91 | SVC | SVC |
| **scPoli** | **sample** | **30** | **20** | **1&2** | 0.84 | 0.3 | 0.77 | 0.14 | 0.8 | 0.19 | 0.88 | 0.45 | 0.84 | 0.29 | 0.96 | 0.93 | SVC | SVC |
| **scGen** | **cell** | **30** | **0** | **2** | 0.96 | 0.3 | 0.92 | 0.12 | 0.93 | 0.17 | 0.98 | 0.45 | 0.94 | 0.26 | 0.99 | 0.95 | SVC | SVC |
| **scPoli** | **cell** | **100** | **100** | **1&2** | 0.96 | 0.29 | 0.9 | 0.15 | 0.93 | 0.17 | 0.98 | 0.48 | 0.94 | 0.24 | 0.99 | 0.95 | LSVC | LSVC |
| **scANVI** | **cell** | **20** | **0** | **2** | 0.95 | 0.29 | 0.9 | 0.11 | 0.91 | 0.15 | 0.97 | 0.45 | 0.93 | 0.23 | 0.99 | 0.96 | SVC | SVC |
| **scPoli** | **cellSample** | **30** | **100** | **1&2** | 0.74 | 0.28 | 0.58 | 0.14 | 0.69 | 0.17 | 0.78 | 0.44 | 0.76 | 0.28 | 0.93 | 0.91 | SVC | SVC |
| **scPoli** | **cell** | **20** | **100** | **1&2** | 0.92 | 0.27 | 0.79 | 0.12 | 0.87 | 0.14 | 0.96 | 0.48 | 0.9 | 0.2 | 0.99 | 0.97 | SVC | SVC |
| **scGen** | **cell** | **100** | **0** | **2** | 0.98 | 0.27 | 0.95 | 0.12 | 0.96 | 0.17 | 0.99 | 0.46 | 0.97 | 0.24 | 1.0 | 0.95 | SVC | SVC |
| **scPoli** | **sample** | **30** | **100** | **1&2** | 0.7 | 0.27 | 0.56 | 0.15 | 0.65 | 0.19 | 0.75 | 0.44 | 0.72 | 0.29 | 0.89 | 0.91 | SVC | SVC |
| **scPoli** | **sample** | **20** | **20** | **1&2** | 0.91 | 0.26 | 0.87 | 0.15 | 0.89 | 0.19 | 0.93 | 0.45 | 0.91 | 0.28 | 0.97 | 0.92 | SVC | SVC |
| **scANVI** | **cell** | **200** | **0** | **2** | 0.97 | 0.25 | 0.96 | 0.12 | 0.95 | 0.16 | 0.98 | 0.47 | 0.96 | 0.22 | 0.99 | 0.96 | LSVC | SVC |
| **scGen** | **cell** | **200** | **0** | **2** | 0.97 | 0.25 | 0.92 | 0.12 | 0.95 | 0.15 | 0.99 | 0.45 | 0.96 | 0.23 | 1.0 | 0.94 | SVC | SVC |
| **scGen** | **cell** | **50** | **0** | **2** | 0.98 | 0.24 | 0.95 | 0.11 | 0.96 | 0.14 | 0.99 | 0.45 | 0.97 | 0.22 | 1.0 | 0.94 | SVC | SVC |
| **scPoli** | **cellSample** | **30** | **20** | **1&2** | 0.91 | 0.24 | 0.91 | 0.11 | 0.88 | 0.14 | 0.93 | 0.46 | 0.9 | 0.21 | 0.97 | 0.95 | SVC | SVC |
| **scPoli** | **cell** | **20** | **50** | **1&2** | 0.89 | 0.23 | 0.74 | 0.09 | 0.82 | 0.11 | 0.94 | 0.44 | 0.85 | 0.17 | 0.99 | 0.96 | SVC | SVC |
| **scPoli** | **cell** | **30** | **100** | **1&2** | 0.91 | 0.23 | 0.83 | 0.08 | 0.86 | 0.11 | 0.95 | 0.49 | 0.89 | 0.15 | 0.99 | 0.98 | LSVC | LSVC |
| ➤ **scANVI** | **cell** | **30** | **0** | **2** | 0.98 | 0.23 | 0.95 | 0.12 | 0.96 | 0.17 | 0.98 | 0.5 | 0.97 | 0.21 | 0.99 | 0.97 | SVC | SVC |
| **scPoli** | **cellSample** | **20** | **20** | **1&2** | 0.89 | 0.21 | 0.84 | 0.1 | 0.84 | 0.11 | 0.92 | 0.43 | 0.88 | 0.2 | 0.98 | 0.93 | SVC | SVC |
| **scPoli** | **cellSample** | **100** | **100** | **1&2** | 0.68 | 0.2 | 0.62 | 0.14 | 0.61 | 0.16 | 0.75 | 0.44 | 0.69 | 0.24 | 0.91 | 0.9 | SVC | SVC |
| **scANVI** | **cell** | **100** | **0** | **2** | 0.96 | 0.2 | 0.93 | 0.11 | 0.93 | 0.14 | 0.97 | 0.47 | 0.94 | 0.2 | 0.99 | 0.96 | SVC | SVC |
| **scPoli** | **sample** | **50** | **20** | **1&2** | 0.8 | 0.18 | 0.7 | 0.12 | 0.74 | 0.12 | 0.87 | 0.47 | 0.79 | 0.19 | 0.94 | 0.95 | SVC | SVC |
| **scPoli** | **cellSample** | **50** | **100** | **1&2** | 0.71 | 0.16 | 0.52 | 0.11 | 0.64 | 0.11 | 0.83 | 0.45 | 0.72 | 0.19 | 0.93 | 0.93 | SVC | SVC |
| **scANVI** | **cell** | **50** | **0** | **2** | 0.96 | 0.16 | 0.91 | 0.11 | 0.93 | 0.13 | 0.97 | 0.46 | 0.94 | 0.19 | 0.99 | 0.95 | SVC | SVC |
| **scPoli** | **cellSample** | **50** | **20** | **1&2** | 0.83 | 0.16 | 0.74 | 0.09 | 0.78 | 0.1 | 0.89 | 0.45 | 0.82 | 0.16 | 0.96 | 0.95 | SVC | SVC |
| **scPoli** | **sample** | **50** | **100** | **1&2** | 0.69 | 0.16 | 0.55 | 0.12 | 0.63 | 0.11 | 0.76 | 0.44 | 0.71 | 0.2 | 0.9 | 0.92 | SVC | SVC |
| **scPoli** | **cellSample** | **20** | **50** | **1&2** | 0.83 | 0.14 | 0.77 | 0.11 | 0.78 | 0.11 | 0.88 | 0.44 | 0.82 | 0.19 | 0.94 | 0.92 | SVC | SVC |
| **scPoli** | **sample** | **20** | **50** | **1&2** | 0.64 | 0.13 | 0.4 | 0.09 | 0.57 | 0.08 | 0.74 | 0.44 | 0.64 | 0.15 | 0.95 | 0.94 | LSVC | LSVC |
| **scPoli** | **sample** | **100** | **20** | **1&2** | 0.83 | 0.12 | 0.79 | 0.11 | 0.78 | 0.1 | 0.87 | 0.44 | 0.83 | 0.19 | 0.94 | 0.91 | SVC | SVC |
| **scPoli** | **cell** | **20** | **20** | **1&2** | 0.93 | 0.12 | 0.86 | 0.06 | 0.9 | 0.04 | 0.95 | 0.41 | 0.92 | 0.08 | 0.99 | 0.96 | SVC | SVC |
| **scPoli** | **cellSample** | **100** | **20** | **1&2** | 0.86 | 0.1 | 0.87 | 0.09 | 0.83 | 0.05 | 0.89 | 0.24 | 0.86 | 0.15 | 0.95 | 0.9 | SVC | SVC |
| **random** | **na** | **nan** | **nan** | **NaN** | 0.2 | 0.1 | 0.09 | 0.1 | -0.0 | -0.0 | 0.21 | 0.21 | 0.19 | 0.11 | 0.81 | 0.89 | rand | rand |

Un. p. = Unseen patient dataset
Un. s. = Unseen studies dataset
NU = Not Utilized

**Annotation (annot)**
1&2 = Level 1 and 2
2 = Level 2

**Performance metrics**
wF1 = weighted F1
BAS = Balanced accuracy score
MCC = Matthew Correlation Coef.
wPRE = weighted Precision
wREC = weighted Recall
wSPE = weighted Specificity

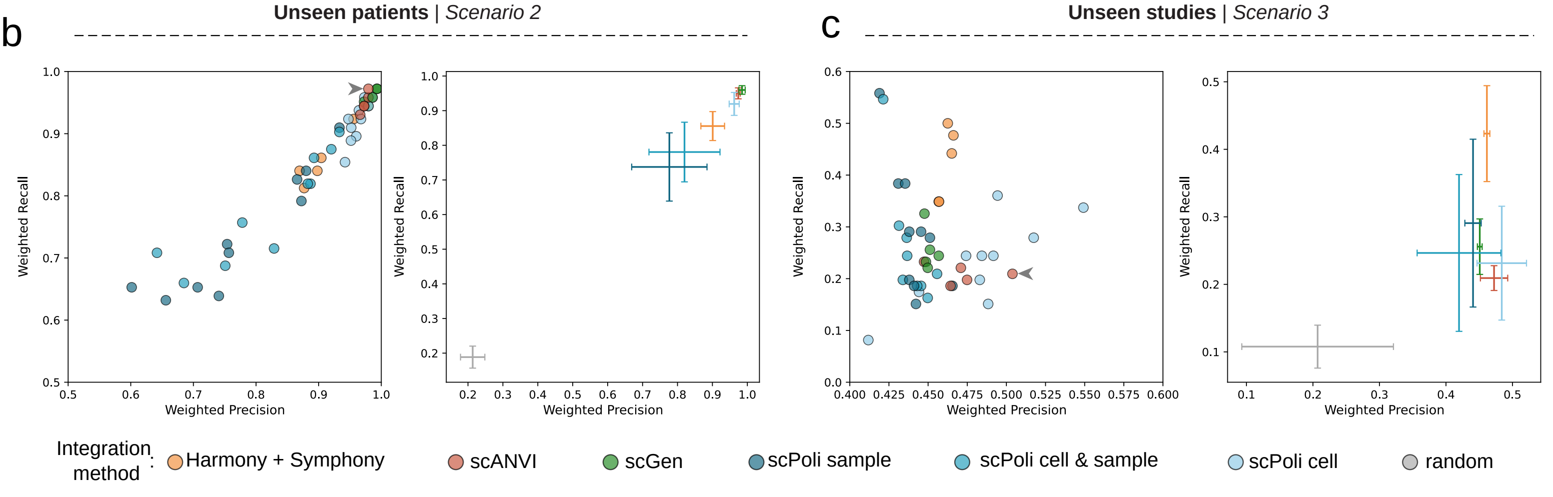

Ext F10

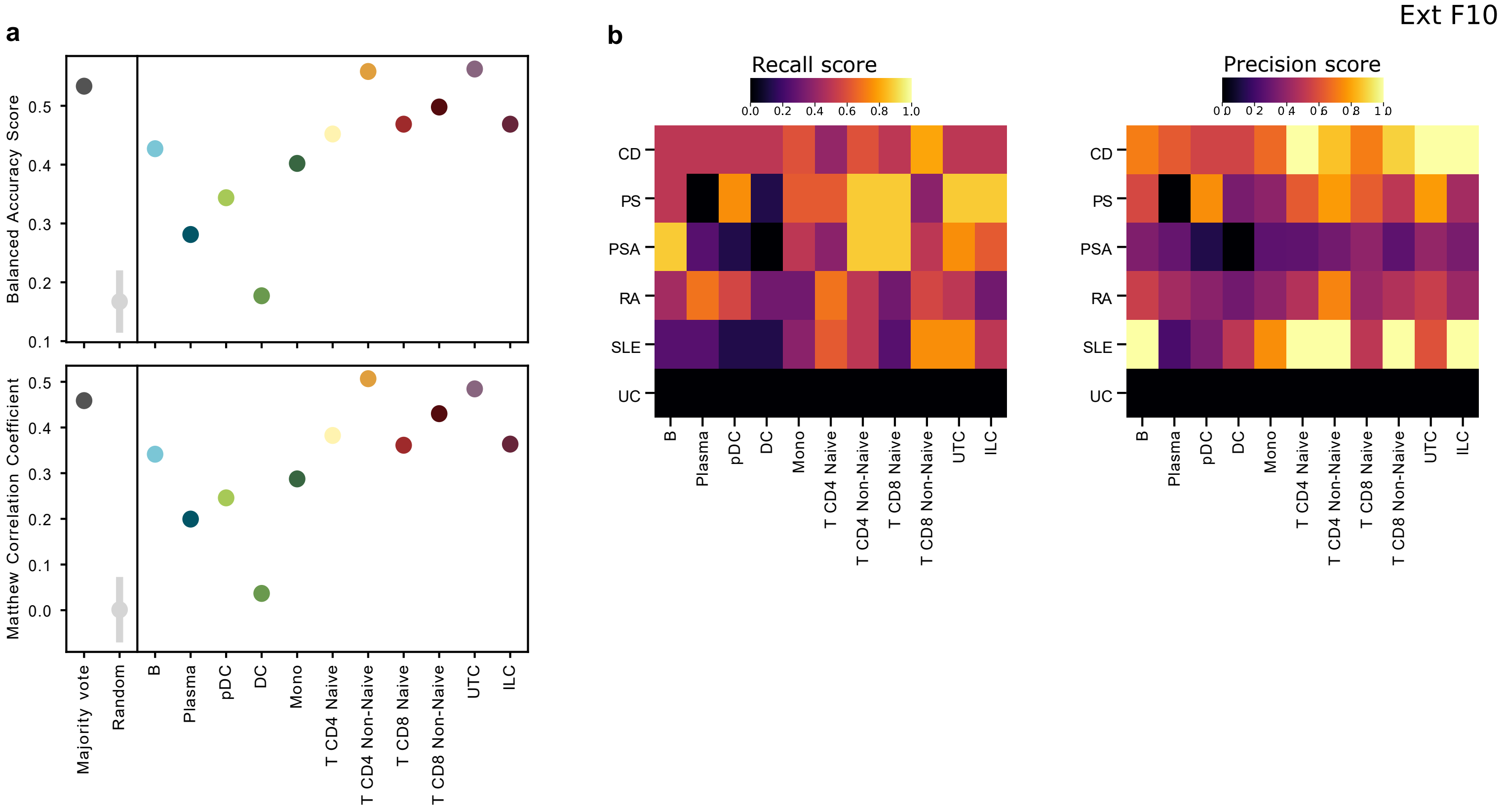
a
Balanced Accuracy Score
Matthew Correlation Coefficient
Majority vote
Random
B
Plasma
pDC
DC
Mono
T CD4 Naive
T CD4 Non-Naive
T CD8 Naive
T CD8 Non-Naive
UTC
ILC
b
Recall score
Precision score
CD
PS
PSA
RA
SLE
UC

Ext F11

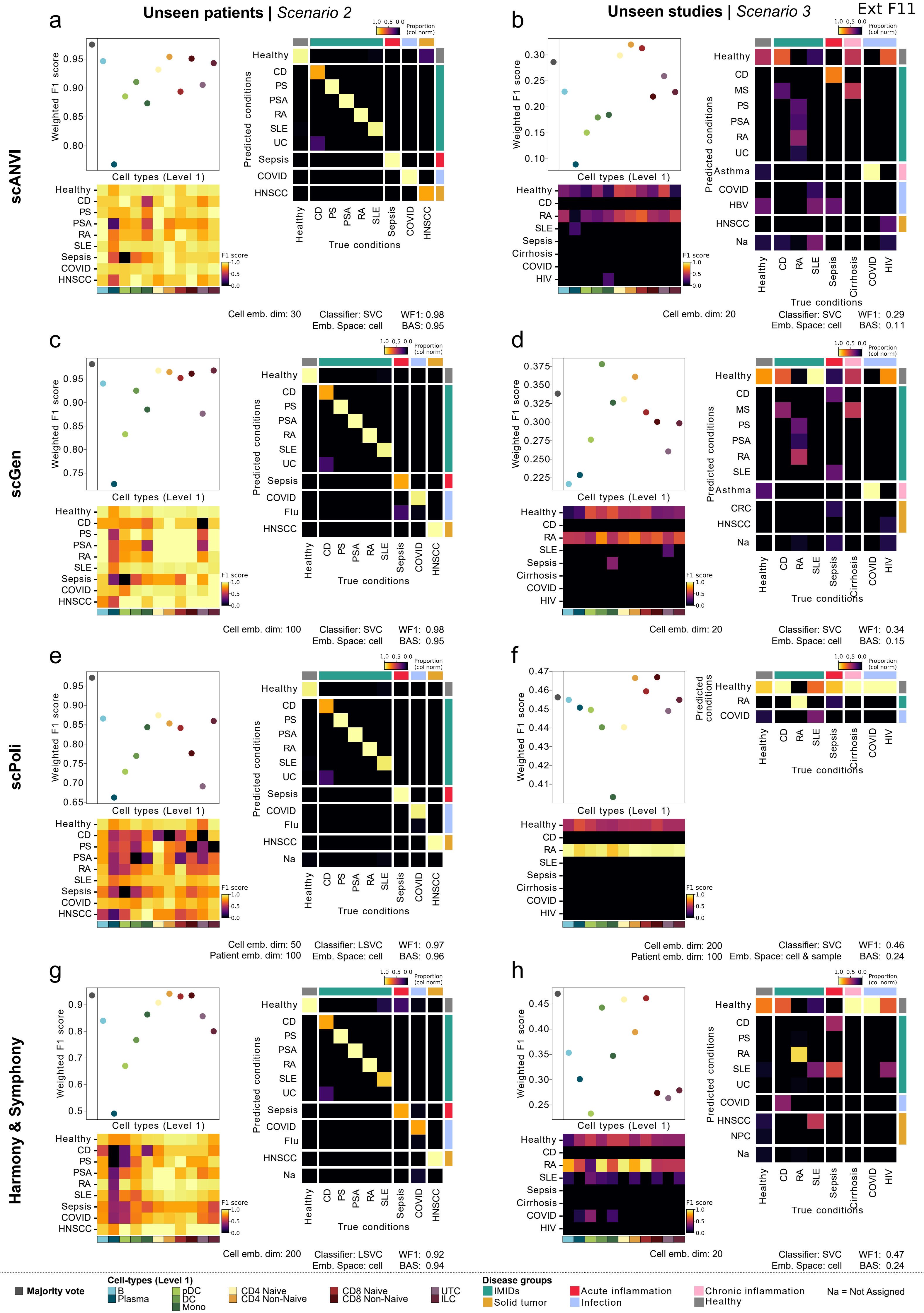

a
Unseen patients | Scenario 2
b
Unseen studies | Scenario 3
scANVI
scGen
scPoli
Harmony & Symphony
Weighted F1 score
Cell types (Level 1)
Predicted conditions
True conditions
Proportion (col norm)
F1 score
Healthy
CD
PS
PSA
RA
SLE
UC
Sepsis
COVID
Flu
HNSCC
MS
Asthma
HBV
Na
Cirrhosis
HIV
CRC
NPC
Cell emb. dim: 30
Classifier: SVC
Emb. Space: cell
WF1: 0.98
BAS: 0.95
Cell emb. dim: 20
Classifier: SVC
Emb. Space: cell
WF1: 0.29
BAS: 0.11
c
d
Cell emb. dim: 100
Classifier: SVC
Emb. Space: cell
WF1: 0.98
BAS: 0.95
Cell emb. dim: 20
Classifier: SVC
Emb. Space: cell
WF1: 0.34
BAS: 0.15
e
f
Cell emb. dim: 50
Patient emb. dim: 100
Classifier: LSVC
Emb. Space: cell
WF1: 0.97
BAS: 0.96
Cell emb. dim: 200
Patient emb. dim: 100
Classifier: SVC
Emb. Space: cell & sample
WF1: 0.46
BAS: 0.24
g
h
Cell emb. dim: 200
Classifier: LSVC
Emb. Space: cell
WF1: 0.92
BAS: 0.94
Cell emb. dim: 20
Classifier: SVC
Emb. Space: cell
WF1: 0.47
BAS: 0.24
Majority vote
Cell-types (Level 1)
B
Plasma
pDC
DC
Mono
CD4 Naive
CD4 Non-Naive
CD8 Naive
CD8 Non-Naive
UTC
ILC
Disease groups
IMIDs
Solid tumor
Acute inflammation
Infection
Chronic inflammation
Healthy
Na = Not Assigned

### Supplementary Table Legends (1-7)

**Supplementary Table 1. Dataset overview of human PBMCs samples**. This table contains general information regarding the datasets and the clinical information of the samples included in the current study. (**Sheet 1: byStudyID**) Details on the dataset (studyID), where the data has been generated (in-house or public), the 10X Genomics chemistry, the publication and the dataset reference (in case of public data), and if we have remapped the FASTQ files. In all cases, we provide the CellRanger and Reference Genome version used. Additionally, for each disease, we provide the number of donors collected before the quality control. (**Sheet 2: byDisease_splitted**) Summary of the number of patients per disease and stratified by subsets (Main, unseen patients or unseen studies), considering sex and binned age categories. (**Sheet 3: bySampleID_afterQC**) Details regarding the technical and clinical metadata per sample; for the missing metadata information (NA is displayed). (**Sheet 4: SCGT00_CentralizedDataset**) Details of samples from a unified, centralized study of the patient cohort, processed by sample pools (patientPool) and stratification into Reference and Query subsets.

**Supplementary Table 2. Comparison between the inflammation atlas and dataset specific (published) annotations**. This table contains the Adjusted Rand Index (ARI) between each cell identifier grouped in our atlas (*Level 2*) and each cell identifier assigned by the external annotation extracted from the published single-cell atlas, using the highest resolution available in their publication. Also, we provide the contingency tables between external annotation (reference) and grouping our internal annotation to match their level of resolution. From **Sheet 1-4**, public single-cell atlas such as Perez *et al.*[27], Ahren *et al.*[99], Terekhova *et al.*[101] and Ren *et al.*[100], respectively.

**Supplementary Table 3. Overview of the clustering and annotation strategy, parameters applied and cell type marker genes.** (**Sheet 1: Annotation Strategy**) This table contains the clustering details (*scVI* number of latents, number of neighbors and resolution parameter) as well as annotation labels (Level 1 and 2) obtained after recursive integration, and clustering steps. (**Sheet 2: Level1_all**) List of gene markers obtained from the differential expression analysis (DEA) to annotate populations at Level 1. (**Sheet 3-11: Level2_CellTypeLevel1**) Each sheet contains the list of gene markers obtained from the DEA to annotate Level 2 populations. (**Sheet 12: Annotation Strategy SCGT00_CentralizedDataset**) This table contains the clustering details (*scVI* number of latents, number of neighbors and resolution parameter) as well as annotation labels (*Level 1*) obtained after recursive integration, and clustering steps.

**Supplementary Table 4. Gene signatures describing the full spectrum of an inflammatory process**. (**Sheet 1: Spectra input gene sets**) This table contains the input gene sets stratified by cell type and global. These gene sets include (a) a list of cell type-specific signatures derived from Spectra - Cytopus[25,119] as well as (b) our curated list of nine inflammatory signatures based on existing literature [35–41], that were integrated into the gene sets of each Level 1 cell type. (**Sheet 2: Spectra Factor markers**) This table contains selected gene markers for each Spectra factor. For additional information on the strategy to select markers see ***Methods***.

**Supplementary Table 5. Evaluation of Spectra inflammation-related signatures.** This table contains the results of the Linear Mixed Effect Model (LMEM) implemented to evaluate the Spectra immune-related signature estimates for each disease compared to healthy patients, while correcting for *chemistry* and *studyID*. (**Sheet 1: Level1 annotation**) LMEM model to evaluate Spectra signatures considering cell types annotated at Level1. (**Sheet 2: Level2 annotation**) LME model to evaluate Spectra signatures considering cell types annotated using fine-grained annotation, Level2.

**Supplementary Table 6. Gene regulatory Network and TF activity.** This table contains the results related with the Gene Regulatory Network analysis of the IFN-induced Spectra factors. (**Sheet 1: CollecTRI-Spectra**): Integration of the CollecTRI Gene Regulatory Network with Spectra factors based on the target genes. source_collectri = Transcription Factor (TF); weight = Binding weights for the TF-gene interactions; source_net: ID of Spectra signature; Factor_celltype: cell type where the Spectra signature was identified; Factor_function: pathway enriched in the Spectra signature; value: score of Spectra signature. (**Sheet 2: TF activity**): Results showing the activity of TFs associated with each Spectra signature. We calculated the activity of each TF by cell type and by disease, using only common genes between each Spectra signature and each TF. Activity and p-values are reported after running the ULM method from decoupleR[42]. (**Sheet 3: pval SLE_Level2**): Results for the comparison of the TF activity across Level2 subpopulations and correction with Benjamini-Hochberg. (**Sheet 4: MOFA loadings**): Gene loadings of MOFAcell results. (**Sheet 5: pval SLE_Flare**): Results for the comparison of the TF activity across Flare and non Flare SLE patients from Perez *et al.*[27] using a T-test and correction with Benjamini-Hochberg.

**Supplementary Table 7**. **Parameters Utilized in Integration, Mapping, and Classification Tasks.** This table reports the parameters employed for executing integration, mapping, and classifier optimization tasks. (**Sheet1: Data Integration for annotation**): Parameters used with scVI for integrating the Main and Centralized dataset at each annotation step, the parameters used with scANVI for integrating the Main dataset post cell annotation, and the parameters used to extract the batch-corrected expression from the scANVI model. (**Sheet 2: Patient Classifier**): Combination of parameters used for integrating a Reference and mapping a Query dataset under each scenario implemented to evaluate the performance of the patient classifier pipeline, Centralized dataset analysis, and the comparison of integration methods. (**Sheet 3: Hyperparameter optimization**): Parameters and their corresponding range of values that were explored to optimize the Cell-wise and Patient-wise classifiers.
